# A spatial code governs olfactory receptor choice and aligns sensory maps in the nose and brain

**DOI:** 10.1101/2025.05.02.651738

**Authors:** David H. Brann, Tatsuya Tsukahara, Cyrus Tau, Dennis Kalloor, Rylin Lubash, Lakshanyaa Thamarai Kannan, Nell Klimpert, Mihaly Kollo, Martín Escamilla-Del-Arenal, Bogdan Bintu, Thomas Bozza, Sandeep R. Datta

## Abstract

Although topographical maps organize many peripheral sensory systems, it remains unclear whether olfactory sensory neurons (OSNs) choose which of the ∼1100 odor receptors (ORs) to express based upon their spatial location in the olfactory epithelium (OE) or instead ORs are scattered randomly. Here we reveal that each OR is expressed at a precise mean position along the OE dorsoventral axis, thereby instantiating a receptor map. This patterning reflects the differential use, by precursors and mature OSNs, of a coherent gene expression program controlled by a spatially-varying retinoic acid gradient; this program — which includes key transcription factors and axon guidance genes — translates position into a spatially appropriate distribution of OR choices and aligns the epithelial map of OR identity with the glomerular map present in the olfactory bulb. These results identify a transcriptional code that distinguishes and spatially organizes the vast array of sensory channels that comprise the olfactory system.

## Introduction

The nervous system uses space to map sensation^1,2^. The cochlea, for example, preferentially encodes characteristic frequencies at different positions along its unrolled length; this tonotopic map is faithfully propagated through patterned projections to the brainstem, thalamus, and cortex, thereby creating a series of auditory maps in which physical space is used to represent a key stimulus attribute^3^. A similar cascade of maps — originating in the sensory periphery and running through a hierarchy of higher brain centers — supports touch and vision^4,5^. In the olfactory system, odors are detected by mature olfactory sensory neurons (OSNs) in the nasal olfactory epithelium (OE), each of which expresses only one of the 1,172 functional odor receptors (ORs) encoded in the mouse genome^6,7^. However, the extent to which OR expression is spatially organized in the OE into a stereotyped peripheral sensory map remains unclear^8–11^. Addressing this foundational question is critical for our understanding of the sense of smell, as the answer has significant implications for how OSNs choose their receptors, encode information about odors, and find their targets in the higher brain.

Three challenges have vexed our understanding of whether OSNs are spatially patterned in the epithelium. First, we have only limited knowledge of how epithelial position might modulate the differentiation of precursors into mature, OR-expressing OSNs^12^. While there are a small number of known OSN cell lineages (each associated with the expression of ORs belonging to a different receptor gene class)^13,14^, whether spatial position restricts the possible fates of OSN precursors or influences OR choice remains uncertain^15^. Second, understanding the relationship between gene expression and physical location is difficult in the OE, because the underlying cartilaginous turbinates transform an otherwise smooth and continuous epithelial sensory sheet into a complex labyrinth^16^, challenging analysis of spatial order. Third, most of what we know about spatial patterning of OR expression arises from in situ hybridization and related techniques, which can query only a few genes at once and which are notoriously difficult to align across epithelial samples derived from different individuals^17–20^; although recent microdissection-based efforts have been made to deduce the locations of ORs, the resulting spatial estimates are indirect^21,22^.

Despite these constraints, prior experiments have suggested a working model in which individual receptors are assigned to one of between four and nine dorsoventral “zones” in the OE, but are randomly chosen for expression by individual OSNs in each zone^17–21,23^; as a consequence, every receptor is thought to be expressed uniformly within each zone. Because the OE continuously regenerates during adulthood from a population of basally-localized stem cells, this zonal model implies that pre-choice OSN precursors must acquire a zone-specific cell identity in order to limit OR choices to the appropriate subset^24^. Consistent with this possibility, OSN precursors appear fated to choose ORs that are limited to a single zone, as the OR “switches” occasionally observed in late-stage precursors are restricted to ORs belonging to the same epithelial zone^25–27^. In addition, a handful of marker genes have been identified that distinguish the dorsal-most zone from the rest of the epithelium^28,29^; furthermore, OSNs expressing Class II ORs — which constitute the majority of ORs in the genome — are mostly restricted to ventral zones, whereas OSNs expressing the much smaller number of Class I ORs receptors are strongly biased towards the dorsal-most zone^25,30^.

However, to date no genes have been identified that distinguish the different ventral zones, and it is therefore unclear how OSN precursors might restrict OR choice to the appropriate set of ∼150–250 ORs per zone^31^. Any such restrictions must ultimately influence the mechanisms that enable OSN precursors to choose which single OR to express, which are not yet fully understood but include: the low-level co-expression of ORs in OSN precursors; intra- and inter-chromosomal interactions between “Greek Island” enhancers and OR genes, which ultimately drives high-level expression of a single OR allele; and heterochromatin-mediated silencing of ORs that were not ultimately chosen for expression^7,32–37^. It has recently been suggested that the zone in which a given OSN precursor resides might limit the set of ORs available for co-expression^32^, with OSNs in ventral zones only having receptors from its own and more dorsal zones available for choice, although precisely how zones are translated into a set of co-expression- and choice-qualified ORs remains unknown.

Information from OSNs is projected to insular structures called glomeruli in the olfactory bulb, the first waystation for olfactory processing in the brain; all OSNs of a given subtype (i.e., those expressing the same OR) innervate one of a small number of glomeruli whose spatial location is largely invariant from animal to animal, thereby creating a stereotyped spatial map of odor identity in the brain^38–41^. Any spatial biases apparent in patterns of OR expression in the nose, therefore, need to be aligned with the detailed map of OR identity apparent in the olfactory bulb^42,43^. The axon guidance genes *Nrp2* and *Robo2* may participate in this process, as they are expressed in complementary gradients in the epithelium and causally regulate the targeting of OSN axons to their appropriate zonal targets along the dorsoventral axis in the bulb^12,44–46^. However, it is not known whether these gradients systematically vary across OSN subtypes, whether this variation is sufficient to ensure proper glomerular targeting, or how the expression of these or any other axon guidance-related genes is coordinated with OR choice^47,48^. Because current models propose that ORs are chosen randomly within each broad zone, a number of mechanisms have also been suggested to enable OSNs to precisely segregate their axons into the correct, OR-specific glomerulus in the bulb based upon the identity of the OR they have chosen. These include OR-dependent regulation of axon guidance gene expression, homophilic interactions among axons expressing the same OR, and repulsive interactions between those expressing other ORs^42,49–52^.

Here we show that the nose harbors a receptor map, one built from ∼1100 distinguishable peaks of OR expression whose mean positions are reliably stereotyped across mice. This patterning is the consequence of a coherent gene expression program expressed in both precursors and mature OSNs that is comprised of ∼250 genes, including key transcription factors, retinoic acid signaling components, and axon guidance molecules. Expression levels of the genes that make up this program systematically vary as a function of dorsoventral spatial position in the epithelium and distinguish all OSN subtypes; the use of this gene expression program (which reflects gradients in retinoic acid signaling in the mesenchyme underneath the OE proper) influences and predicts both OR choice and the spatial location of target glomeruli, and thus not only builds a receptor map in the nose but also aligns this map with the second order map of odor identity apparent in the olfactory bulb. ORs are therefore not chosen at random within a small number of zones; instead, spatial gradients of gene expression are transformed into discrete OR choices and glomerular targets, thereby coordinately organizing the first two stages of the peripheral olfactory system.

## Results

We hypothesized that olfactory sensory neurons (OSNs) contain transcriptional signatures that reflect their spatial location in the olfactory epithelium. If each OSN subtype (defined as the collection of OSNs expressing the same OR) is associated with a specific zone, then one might expect that the OSNs from each zone express one or more zone-specific marker genes (Figure 1A). To test this hypothesis, we used single cell RNA sequencing (scRNA-seq) to characterize the transcriptomes of nearly 5 million total cells in the mouse olfactory epithelium across hundreds of replicates, which yielded a dataset of 2.3 million mature OSNs that collectively expressed 1,100 ORs with a median of 1,300 cells per OSN subtype (Figure S1A, see Methods).

**Figure 1.**
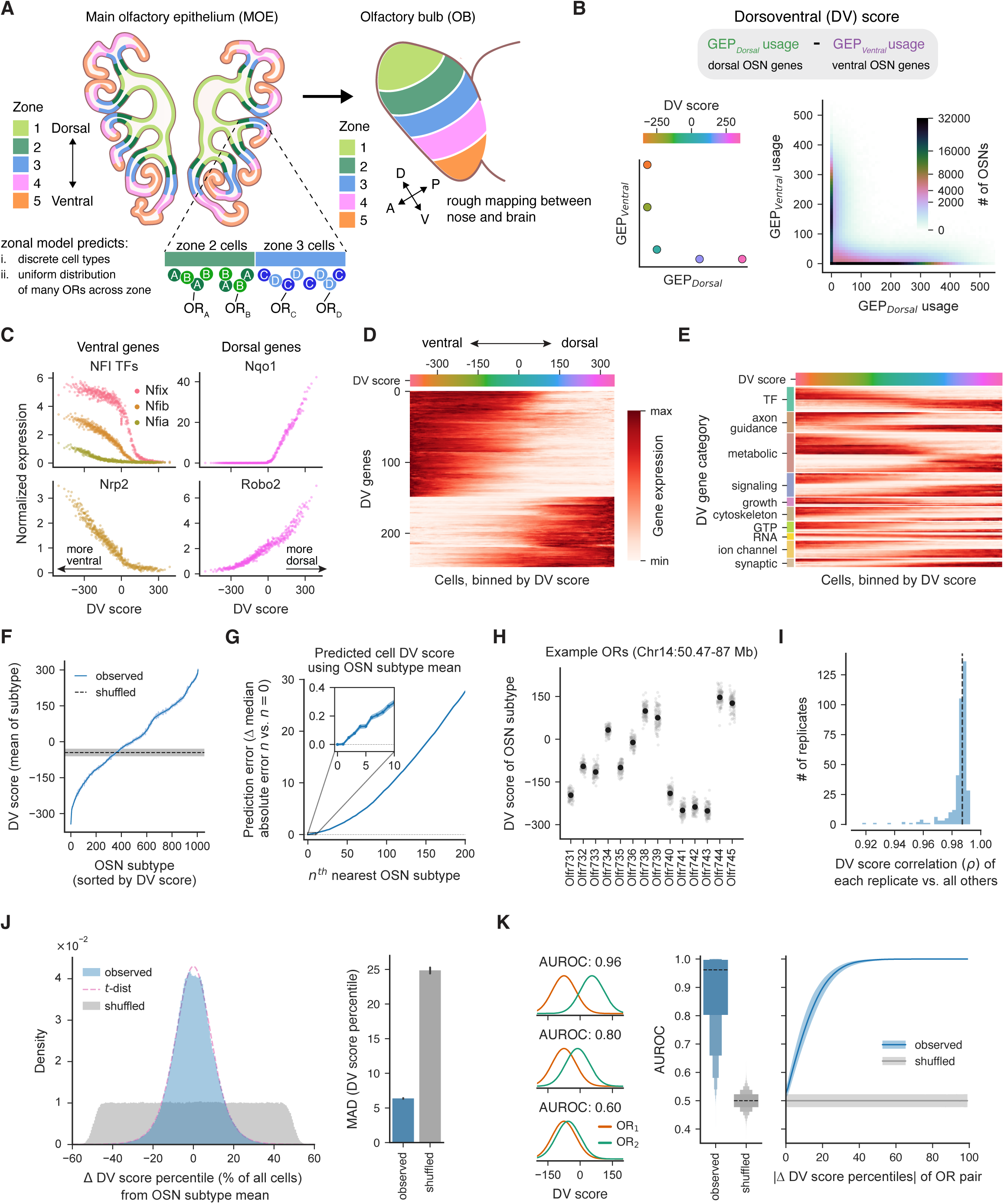
Each OSN subtype is associated with a unique dorsoventral (DV) score. A. Schematic depicting the zonal model for OR choice, and how epithelial zones correspond with positions in the olfactory bulb. B. Gene expression from single-cell RNA-seq (scRNA-seq) data was decomposed into usages for gene expression programs (GEPs) using constrained non-negative matrix factorization (as described in recent work54, see Methods). Schematic (left) depicts how the dorsoventral (DV) score (the difference between GEP_Dorsal_ and GEP_Ventral_) was calculated for each OSN (data shown on right, colormap power transformed for legibility). C. Normalized gene expression of the indicated genes as a function of cellular DV scores. Cells were binned into equal-frequency bins and each dot depicts the average expression in each bin. D. Heatmap depicting the expression of 248 DV-related genes whose expression was correlated with the DV score (see Methods), grouped via hierarchical clustering and plotted as a function of the DV score. E. Similar to (C) but DV genes were organized by their assigned gene category (see Supplemental Table 1 for functional assignments for individual genes). F. DV scores, sorted by the mean for each OSN subtype (defined as the set of OSNs singularly expressing a given OR), for the 1007 OSN subtypes with at least 150 cells in our dataset. G. Error in predicting the DV score for each OSN based upon its expressed OR (using cross-validated linear regression models), based upon the DV score mean (in the training data) of either the same OSN subtype or the means of OSN subtypes with neighboring DV scores. H. The DV scores of OSN subtypes expressing the indicated ORs, which are all located as part of a genomic cluster on chromosome 14. Black dots indicate the mean DV score, and gray dots depict the means when cells from each OSN subtype were split by replicate into 100 separate bins. I. Histogram of the correlation of the mean DV scores for each OSN subtype for each individual replicate and the mean for each OSN subtype from all other replicates. Dotted line depicts the median correlation across replicates. J. (Left) The distribution across all OSNs of the difference of each cell’s DV score from the mean of its respective OSN subtype. Equal numbers of cells were subsampled for each OSN subtype, and the distribution was plotted as a function of the percentile-normalized DV score, and this distribution was fit with a t-distribution. “Shuffled” depicts the distribution when OR labels are shuffled across cells. (Right) The median absolute deviation (MAD) of the respective distributions depicted on the left. Error bars depict bootstrapped 95% confidence intervals of the mean. K. (Left to right) Schematic showing the area under the receiver operating characteristic curve (AUROC) for several example synthetic distributions of DV scores associated with individual cells expressing one of two ORs; the distribution of AUROCs for all pairs of OSN subtypes (black dashed line indicates median, boxes represent 25th/75th percentile, 12.5th/87.5th percentile, and so forth across 1007C2 = 506,521 pairs); and the AUROC as a function of the difference in DV scores (as a percentile of all OSN subtypes) of the pair (with lines and shading reflecting the mean and interquartile range for the pairs at each percentile). Shuffles depict the AUROCs when OR labels are shuffled across cells.

Despite the large size of our dataset, we were unable to identify genes expressed in mature OSNs that uniquely marked each of the proposed zones in the ventral epithelium (Figure S1B). However, our analysis did recover several previously identified genes (e.g., *Nqo1*, *Nfia*) whose expression differed between the dorsal-most and the more ventral zones, suggesting a broad transcriptional bifurcation between the dorsal and ventral epithelium^32,53^.

To better characterize this distinction, we subjected our scRNA-seq data to consensus non-negative matrix factorization (cNMF), which identifies gene expression programs (GEPs) that are each composed of ∼100-200 individual genes whose expression levels co-vary across different OSNs^54,55^. cNMF identified two GEPs that we provisionally named GEP_Dorsal_ and GEP_Ventral_, as they included a handful of genes known to be restricted to the dorsal or ventral epithelium^54^, as well as many others including transcription factors, axon guidance molecules, and cytoskeletal regulatory proteins (Figure 1A, Supplemental Table 1). The overall level of expression of the genes participating in a given GEP can be summarized in a single number, the GEP “usage”; because individual OSNs expressed genes belonging to GEP_Dorsal_ or GEP_Ventral_ but not both, we could further summarize the usage of these two GEPs through a single metric called the dorsoventral (DV) score (i.e., the usage of GEP_Dorsal_ subtracted from that of GEP_Ventral_, Figure 1B).

As expected, known dorsal marker genes like *Nqo1* were expressed in OSNs with high DV scores, whereas ventral marker genes like the *NFI* family transcription factors were expressed in OSNs with low DV scores (Figure 1C). However, *Nqo1* expression was not uniform across those OSNs that expressed *Nqo1*, as would be expected if they were organized into a homogenous dorsal “zone”. Instead *Nqo1* expression tonically increased as a function of DV scores, suggesting that *Nqo1* expression varies in a gradient-like fashion across OSNs; a similar expression gradient was observed in ventral cells expressing *NFI*. Notably, both *Nqo1* and *NFI* genes themselves participate in the DV score, suggesting that other GEP_Dorsal_ and GEP_Ventral_ genes may be similarly organized into smooth transcriptional gradients. Indeed, expression of the axon guidance gene *Robo2* (as well as the other ∼100 genes participating in GEP_Dorsal_) tonically increased as a function of OSN DV scores, while expression of the axon guidance gene *Nrp2* (as well as the other ∼150 genes participating in GEP_Ventral_) tonically decreased^12,44–46^ (Figures 1C–E). Thus OSNs harbor an organized gradient of transcriptional differences, one imposed by systematic variation in the use of GEP_Dorsal_ and GEP_Ventral_.

Each OSN subtype was associated with a unique average DV score, and DV scores varied smoothly across OSN subtypes from lowest (“most ventral”) to highest (“most dorsal”) (Figures 1F–H and S1C). Although DV scores could be used to categorize OSN subtypes as dorsal (DV score ≥ 60, Figure S1D-E, see Methods) or ventral, we did not observe any additional clustering of DV scores, consistent with OSN gene expression varying continuously rather than discretely as one might expect in a zonal model (Figure S1F). The relationship between each OSN subtype and its associated mean DV score was remarkably stable, as the rank ordering of OSN subtypes based on their DV scores was nearly invariant (rho = 0.987) across the hundreds of independent samples in our sequencing dataset, which encompasses both sexes and a variety of odor environments and experimental manipulations (Figures 1H–I and S1G–H). Despite the reliable relationship between the identity of each OSN subtype and its mean DV score, the individual neurons belonging to a given OSN subtype had different DV scores, which were tightly clustered around the subtype mean with a sharp peak and extended tails (Figures 1J and S1I–J). This distribution was sufficiently peaked that individual OSNs belonging to different subtypes could be effectively distinguished based upon their DV scores alone (median auROC = 0.96, Figure 1K).

Thus, despite variation in DV scores among individual OSNs expressing the same OR, the mean DV score associated with each OSN subtype is stereotyped across mice, is sufficient to differentiate each OSN subtype from all others, and summarizes the coordinated expression of several hundred genes, including a subset of genes known to signify whether an OSN is located in the dorsal or ventral olfactory epithelium.

### DV scores predict the unique spatial location of each OSN subtype and OR

Given the continuous variation observed in DV gene expression across OSN subtypes, we hypothesized that the specific dorsoventral location adopted by each OSN is encoded by its DV score. Because each OSN subtype is associated with a specific mean DV score, this proposal makes the strong prediction that the mean location of each OSN subtype — and therefore OR — along the DV axis should be unique, predictable and stereotyped across mice, rather than being randomized within one of a small number of discretized zones. To test this hypothesis, we used multiplexed error-robust fluorescence in situ hybridization (MERFISH) to directly measure the spatial positions of hundreds of OR and DV genes throughout the epithelium^56^ (Figures 2A–B and S2A–B, see Methods).

**Figure 2.**
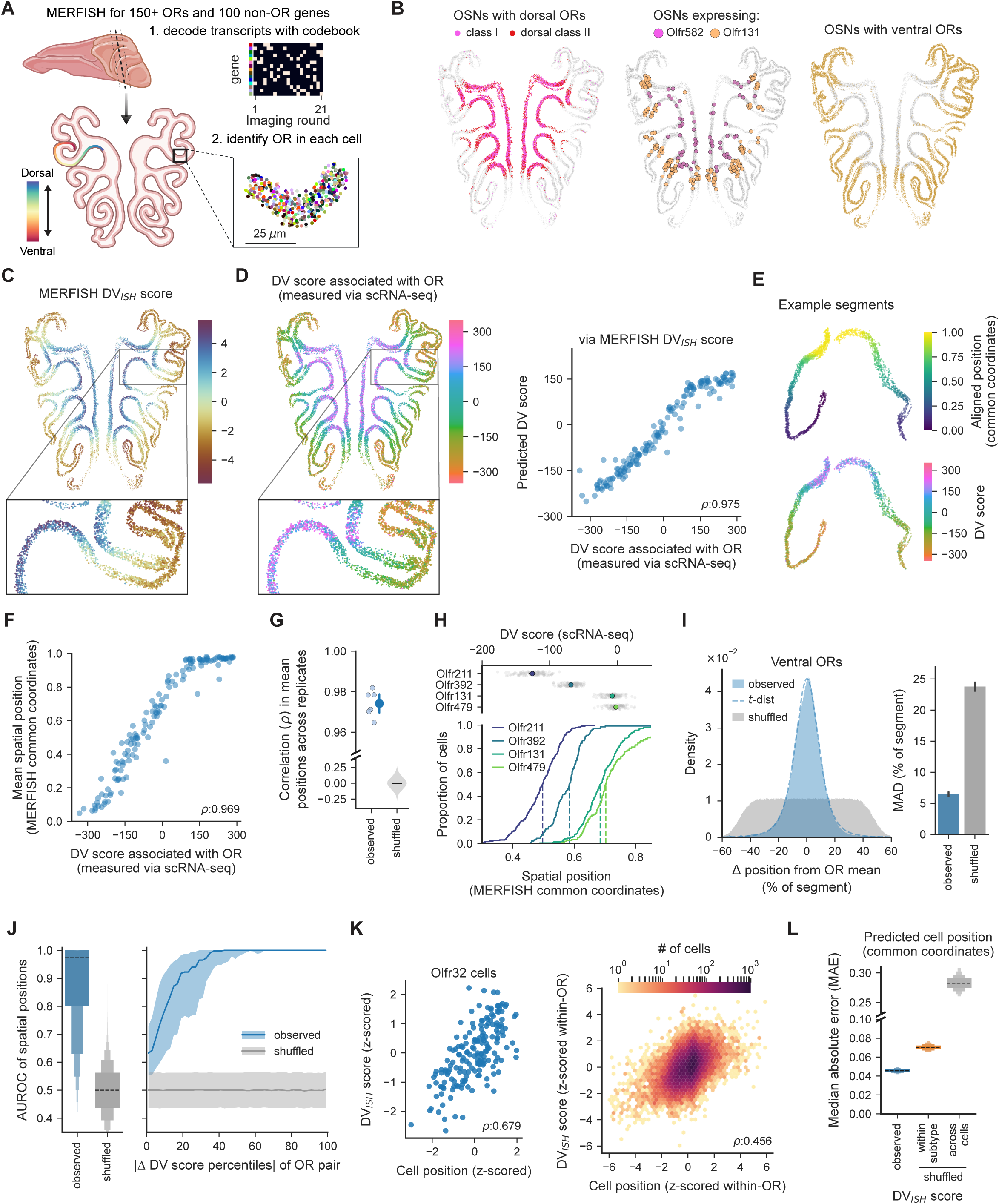
Each OSN subtype occupies a unique region of the epithelium. A. Schematic depicting the use of MERFISH to detect over 150 ORs and ∼100 non-OR genes (including DV genes) in individual coronal sections of the MOE. For reference, the dorsoventral axis is colored along an example epithelial segment. B. From left to right, OSNs expressing dorsal ORs, OSN expressing either the dorsal OR Olfr582 or the ventral OR Olfr131, and OSNs expressing all ventral class II ORs. Note that for visualization purposes, in the middle panel dots have been placed over OSN locations; dots are ∼100x normal size and not weighted by density of the underlying data, and so this representation highlights the tails of the OR distribution. C. OSNs colored by their DV_ISH_ score, defined as the first principal component of the MERFISH-measured expression of DV genes in each OSN. Note, only a subset of the DV genes identified by scRNA-seq were measured via MERFISH. D. (Left) OSNs colored by the DV score associated with their expressed OR, as measured via scRNA-seq. (Right) Predictions of the DV score of each OSN subtype (as identified via scRNA-seq) using regression models trained on the DV_ISH_ scores for each OSN. E. Example of two individual segments showing (top) the common coordinate aligned position (see Methods for alignment details) and (bottom) the DV scores associated with the expressed ORs. F. The mean aligned spatial position for each OSN subtype, as a function of its DV score. G. The correlation of the mean aligned spatial positions across OSN subtypes across pairs of sections (from four individual sections), compared to the correlation when positions were shuffled 10,000 times across OSNs within a section. H. Cumulative distributions of the aligned common coordinate position of cells from four example OSN subtypes (bottom), along with the DV scores as identified via scRNA-seq for each subtype (on top). I. The distribution of the differences in the spatial position of individual OSNs expressing ventral class II receptors (as a percentile of all positions) after subtracting the mean spatial position of its respective OSN subtype. Shuffles depict the distribution when OR labels are shuffled across cells. (Right) The mean absolute deviation (MAD) of the respective distributions, with the mean and bootstrapped 95% confidence interval of the mean. J. The distribution of area under the receiver operating characteristic curves (AUROCs) for the spatial positions of OSNs for all pairs of OSN subtypes (left) and plotted as a function of the difference in DV scores of the pair (right), with lines and shading reflecting the mean and interquartile range for the pairs at each percentile. Shuffles depict AUROCs when OSN subtype labels are shuffled across cells. K. (Left) For OSNs expressing Olfr32, the z-scored DV_ISH_ score as a function of the z-scored spatial position of each cell. (Right) The DV_ISH_ score for all OSNs as a function of its position, where both positions and DV_ISH_ scores were z-scored within each OSN subtype. L. The median absolute error (MAE) of the predicted spatial position of each OSN in the aligned common coordinates, using regression models trained on each cell’s DV_ISH_ score. Predictions were compared to data in which the DV_ISH_ score was shuffled within each OSN subtype, or shuffled across all cells.

Consistent with past in situ hybridizations, individual ORs were expressed in a spatially restricted manner, with the expression of the most-dorsal receptors being broader than more-ventral receptors (Figure 2B). However, simply coloring each OSN based upon its associated patterns of DV gene expression (the DV_ISH_ score, determined directly via MERFISH) revealed a striking and previously unappreciated fine organization to the epithelium: in each turbinate DV_ISH_ scores smoothly and predictably varied from dorsal to ventral along each segment (Figures 2C and S2C). Furthermore, identifying the OR expressed within each OSN and coloring it based upon its DV score (as measured via scRNA-seq) also yielded a similar smooth and continuous gradient (Figures 2D and S2D). We observed a similar relationship between physical space and OSN subtype-specific DV scores when the epithelium was “unrolled” by aligning all segments to a common spatial axis (Figures 2E–G and S2E, see Methods).

Furthermore, the reliable relationships between spatial position, DV scores, and OSN subtype identities enabled accurate predictions of the spatial location of each OSN subtype both in our own data and in five independently obtained datasets in which the position of OSN subtypes could be precisely or approximately assigned (Figures 2F, S2F–H). In contrast, we did not observe any apparent spatial clustering of OSN subtypes in our MERFISH data, although a model with two clusters could indeed recover the broad dorsal and ventral regions (Figure S2I). Notably, in the dorsal-most region the relationship between the mean position of a given OSN subtype and its DV score is less apparent (Figures 2D, 2F), as might be expected given the breadth of the spatial distribution of individual receptors in this epithelial region.

While the mean location of each ventral OSN subtype was distinct, predictable, and stereotyped, the constituent OSNs that belong to a given subtype occupied a distribution of locations (with a sharp peak and long tails encompassing ∼5–15 percent of the epithelium), with the extent of overlap in the spatial distributions of different ventral OSN subtypes reflecting the degree to which their DV scores were similar or different (Figures 1J, 2H–J and S2J). Note that this peaky spatial distribution of OR expression is fundamentally incompatible with zonal models that predict the uniform expression of a given OR within a particular zone. Dorsal OSN subtypes had spatial distributions that were also distinguishable, but their positions are more overlapping and less discriminable than their ventral counterparts (Figure S2J), consistent with the dorsal-most zone being less spatially organized than the rest of the epithelium. Importantly, variation among DV_ISH_ scores for each OSN within a subtype largely reflected spatial location rather than noise, as variation in DV scores among OSNs expressing the same OR reflected relatively more dorsal or ventral positioning (Figure 2K); the reliable relationship between DV scores and the OSN location also enabled accurate predictions of the position of individual OSNs based upon DV scores alone (Figure 2L).

Together, these data reveal that the precise location of each OSN is well-captured by its DV score and that each OSN subtype occupies a unique and restricted spatial distribution in the olfactory epithelium — thereby creating a map with ∼1100 overlapping but distinguishable receptor peaks. This pattern is the consequence of each OR having a preferred spatial position in the epithelium, and those OSNs that reside at each spatial position having a preferred OR.

### Dorsoventral identity is independent of the expressed OR and downstream activity

Two possibilities could explain the tight link between each OR and its associated DV score: the chosen OR could determine the DV score of each OSN subtype or, alternatively, OR choice could be a downstream consequence of each cell’s spatial identity, which is reflected by its DV score. However, OSN DV scores were essentially unchanged after three distinct activity manipulations: naris occlusion, exposure to novel olfactory environments, and knocking out CNGA2, the key ion channel that transforms OR activation into calcium entry and spiking activity (Figures S3A–B)^57^. Indeed epithelial position appears to be functionally decoupled from the activity patterns of OSNs, as odor-evoked activity driven by both acute and chronic exposure to a wide range of monomolecular and home-cage odors triggered patterns of neural activation that were spatially distributed across the entire epithelium (Figure S3C–D); one notable exception was the spatially localized pattern of activity elicited by acute exposure to acids, which largely activate class I ORs in the dorsal region of the epithelium^58^.

To test whether the identity of the expressed OR itself determines DV scores, we took advantage of the OMP-P2 mouse line (Figure S3E), in which the P2 receptor is ectopically expressed in most mature OSNs through a process that frequently silences the previously-chosen OR in each cell^59^. P2 OSNs from OMP-P2 mice adopted a much wider than normal distribution of DV scores (Figures S3F–G), suggesting that DV scores are the consequence of spatial position rather than receptor identity^59^. Consistent with this model, analysis of the ∼15% of OSNs in which P2 was found to be co-expressed with another OR (and which therefore were likely caught in the process of switching from their original OR to P2) revealed that DV scores matched the original OR rather than P2 (Figure S3H–I).

#### Transcriptional signatures of dorsoventral identity are present in OSN precursors

These results suggest that dorsoventral position *per se* imparts an indelible transcriptional identity to each cell. Because OSN precursors reside in an organized layer beneath their mature progeny, it is possible that information about dorsoventral position is present in the olfactory stem cells and intermediate neuronal progenitors (INP) that continuously generate mature OR-expressing OSNs in adult mice (Figure 3A)^60^. To ask whether OSN stem cells or progenitors harbor positional information, we extracted from our scRNA-seq dataset 400,000 differentiating cells before, during, and immediately after OR choice (Figure 3B, see Methods). Ordering these OSN precursors (which exclude late immature and functional OSNs) in pseudotime recapitulated known expression patterns of differentiation-related marker genes and transcription factors that precede OR expression (Figures 3C–D and S4A)^60,61^.

**Figure 3.**
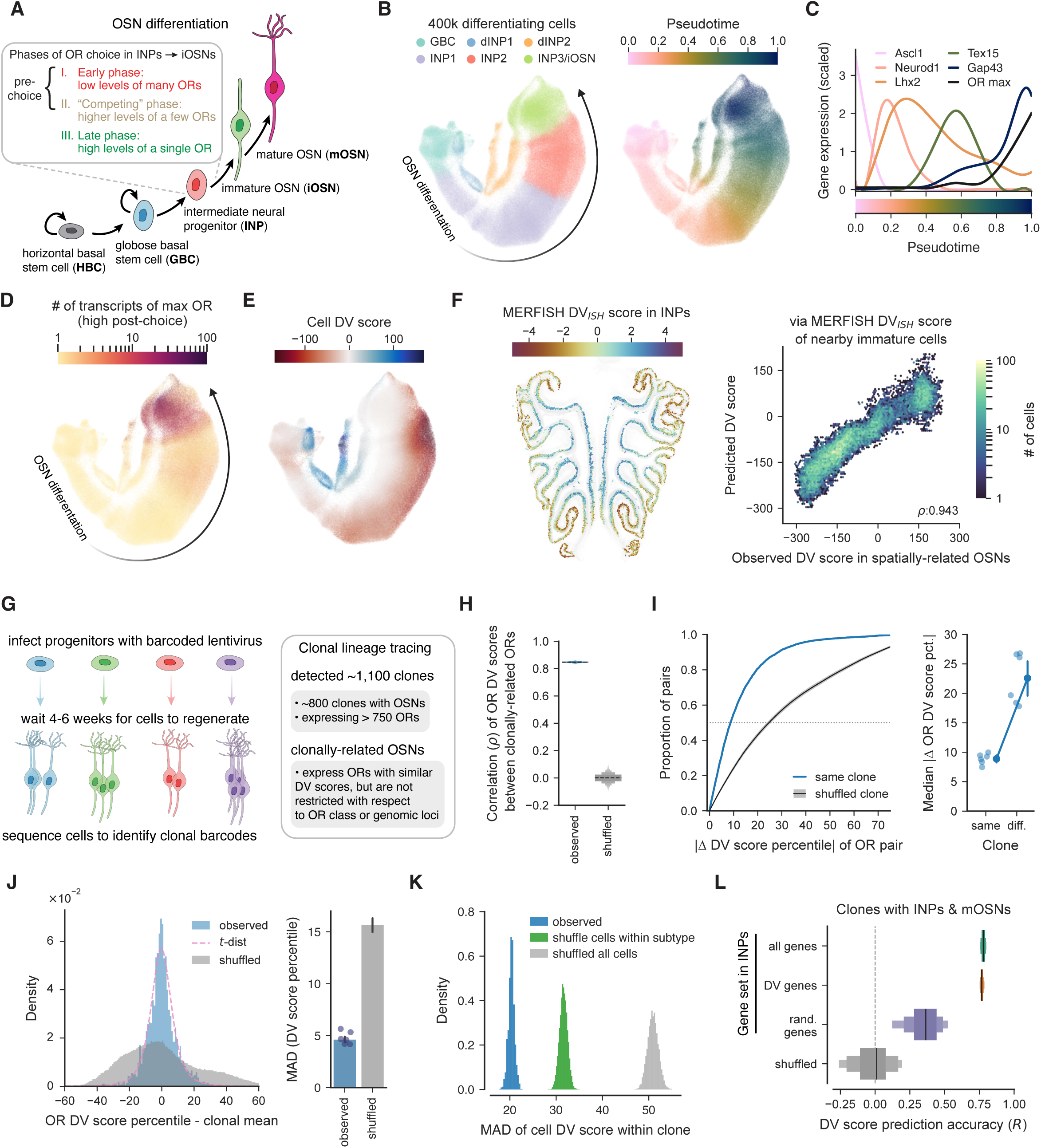
Epithelial position restricts OSN precursor fates and OR choice. A. Schematic depicting the stages of OSN differentiation (which occurs throughout life as OSNs continuously regenerate) as well as the stages of OR co-expression that precede the stable expression of a single OR. B. (Left) UMAP plot depicting 400k precursors and differentiating OSNs, using an integrated dataset from over 360 replicates. Cells are colored by their cluster (dINP = dorsal INP). Note, the majority of iOSNs and all mOSNs are excluded in this dataset. (Right) Shading of UMAP plot based upon pseudotemporal ordering of differentiating OSNs. C. Expression levels of key genes related to OSN differentiation (scaled by their standard deviation), as a function of pseudotime. D. The levels of expression (measured as the number of unique molecular identifiers (UMIs)) of the highest-expressed OR in each cell. E. The DV score for each differentiating cell, calculated using the weights for each gene for GEP_Dorsal_ and GEP_Ventral_ as identified in mature OSNs. Note that the distribution of DV scores is compressed relative to what is observed via scRNA-seq because precursors express a subset of the DV genes expressed in mature OSNs. F. (Left). INP cells, colored by their DV_ISH_ score, as measured via MERFISH. (Right) Predictions (y-axis), via elastic-net regularized models, of the DV score associated with each mature OSN using the DV_ISH_ score of nearby immature cells, as a function of the observed DV scores for each mature OSN subtype (x-axis). G. Schematic depicting the *in vivo* clonal lineage tracing experiments, in which OSN progenitors are labeled with a lentiviral barcode, and then clonally-related progeny are identified using scRNA-seq. H. The correlation in the DV scores associated with the ORs found in clonally-related OSNs. Pairs of OSNs were subsampled across 10,000 restarts and the distributions for the observed data and for data with shuffled clonal labels are shown. I. (Left) The cumulative distribution of the distance in the DV scores associated with each OR for pairs of clonally-related ORs, or for data with shuffled clonal labels, with the mean and bootstrapped 95% confidence interval across shuffles. (Right) The median distance in DV scores of ORs for pairs of OSNs from the same or different clones. Dots indicate individual mice, and error bars indicate the mean and 95% confidence interval of the mean across mice. J. (Left) The distribution of DV scores associated with each OR within a given clone subtracted from the mean of the DV scores associated with each OR within the clone. Shuffles depict the distribution when clonal labels are shuffled across cells. (Right) The median absolute deviation (MAD) of the respective distributions, with the mean and bootstrapped 95% confidence interval of the mean. K. The MAD of the DV scores of clonally-related mature OSNs, compared to data in which clonal labels were shuffled across cells of the same OSN subtype or across all cells. L. Performance of support vector regressor models trained on the indicated gene set in INP cells to predict the DV score associated with the OR detected in each clonally-related mOSN. Shuffles depict the results for DV genes when the DV scores of the ORs expressed in mOSNs was shuffled, and distributions depict the results across 100 restarts.

Many (but not all) DV genes were expressed throughout OSN differentiation, enabling us to compute DV scores for all OSN precursor cell types (see Methods, Supplemental Table 1). We observed a gradient of DV scores across OSN precursors, regardless of their stage of differentiation, which reflected differences in the spatial positioning of precursors within the epithelium itself (Figures 3E–F and S4B–D). Principal components analysis revealed that spatial diversity among precursors (as captured by DV score variation) was the major driver of transcriptional variation among OSN precursors at each developmental stage before final maturation (Figure S4E); this sharply contrasts with our prior results demonstrating that the main axis of transcriptional variation in functional mature OSNs reflects odor-evoked activity (Figure S4F)^54^. Thus, as in mature OSNs, spatial position determines expression levels of GEP_Dorsal_ and GEP_Ventral_ genes at all stages of differentiation, thereby transcriptionally diversifying OSN precursors before they commit to expressing a single OR.

### Clonally-related cells express restricted sets of ORs with similar dorsoventral identities

Our sequencing and MERFISH data demonstrate that the earliest olfactory stem cells express DV genes and therefore have access to information about their position within the epithelium; this information in principle could restrict the OSN subtype(s) each stem cell is capable of ultimately generating. To test this model, we performed *in vivo* clonal lineage tracing by first ablating all mature OSNs with the drug methimazole and then infecting basal stem cells intranasally with a DNA barcoded lentiviral library to uniquely mark each stem cell; methimazole prompts stem cells to synchronously regenerate the mature epithelium^62^, thus allowing us via scRNA-seq to ask about the distribution of ORs generated by each marked stem cell (Figure 3G, see Methods).

After regeneration, OSNs expressed single ORs and harbored transcriptomes similar to those of control cells (Figures S5A–D). Most identified clones (800 out of 1,100) included mature OSNs, which collectively expressed more than 750 different ORs (Figure S5E). These clones revealed that there is not a deterministic 1:1 relationship between stem cell positional identity and the type of OSN that is generated, as individual mature OSNs derived from the same clone often expressed different ORs (Figures S5F–H). However, OSN subtypes (and hence ORs) found within the same clone had much more similar DV scores than those derived from different clones, and mature OSNs generated by the same progenitor expressed the same OR at above chance rates (Figures 3H–J, and S5I–J). These observations are consistent with the process of OR choice in OSN progenitors being biased towards a spatially optimal receptor but being variable at the single cell level, thereby generating a specific (and peaked) distribution of OR choices for any progenitor DV score. Interestingly, the DV scores of clonally-related mature OSNs were even more restricted than might be expected based on their chosen ORs (Figure 3K), suggesting a tighter mapping between epithelial space and DV identity than between epithelial space and OR choice.

As was observed for clonally-related mature OSNs, precursors belonging to the same clone had similar DV scores; furthermore, as precursor DV gene expression could effectively predict the DV scores associated with the receptors expressed by mature OSNs in the same clone (Figures 3L and S5K–L). MERFISH revealed that the DV gene spatial gradients apparent in precursors and mature OSNs were aligned with each other, consistent with OSN precursors residing in one location in the epithelium giving rise to mature OSNs with matched DV scores (Figure 3F). Together, these results demonstrate that olfactory stem cells are fate restricted depending upon their specific location within the olfactory epithelium; this fate restriction biases the choice process towards a spatially appropriate OR — thereby generating a patterned map of OR identity in the nose — but is sufficiently stochastic that often OSN progenitors pick ORs more typically associated with OSN subtypes nearby in space, thereby contributing to the spatial distribution observed for each OSN subtype along the dorsoventral axis of the epithelium.

### Retinoic acid signaling varies spatially and influences OR choice

OSNs isolated from epithelia that have been regenerated after methimazole-mediated ablation exhibited the same relationships between DV scores and OR choice as those from untreated epithelia (Figure S5C), suggesting that the mechanisms that translate space into DV gene expression and choice are either intrinsic to basal stem cells or the result of diffusible cues originating from outside the epithelium. We therefore searched for candidate signaling programs originating in the basal epithelial layer or in the underlying mesenchymal tissue that might reflect dorsoventral position. We observed that multiple genes related to retinoic acid (RA) signaling varied spatially both across the dorsoventral axis of the epithelium and across different epithelial cell types^63–65^ (Figure 4A). For example, the enzyme retinaldehyde dehydrogenase 2 (ALDH1A2), which converts retinaldehyde into RA, was expressed in lamina propria mesenchymal cells, which are located just below the basal stem cells (Figures 4B and S6A). Consistent with prior reports^12,65^, *Aldh1a2* expression (as assessed via MERFISH) was organized into a subepithelial gradient along the dorsoventral axis and was negatively correlated with the DV scores of nearby OSN subtypes (Figures 4C). OSN progenitors and mature OSNs also expressed gradients of other genes related to RA signaling, including retinaldehyde dehydrogenase 3 (*Aldh1a3*), the cellular retinoic acid binding protein 1 (*Crabp1*), the retinoic acid receptor *Rarb* and the retinoid metabolic enzyme *Dhrs3* (Figures S6B–C); in addition, OE cells uniformly expressed many genes capable of responding to RA gradients, including *Rara*, *Rarg*, *Rxrb* and *Rere* (Figure 4A).

**Figure 4.**
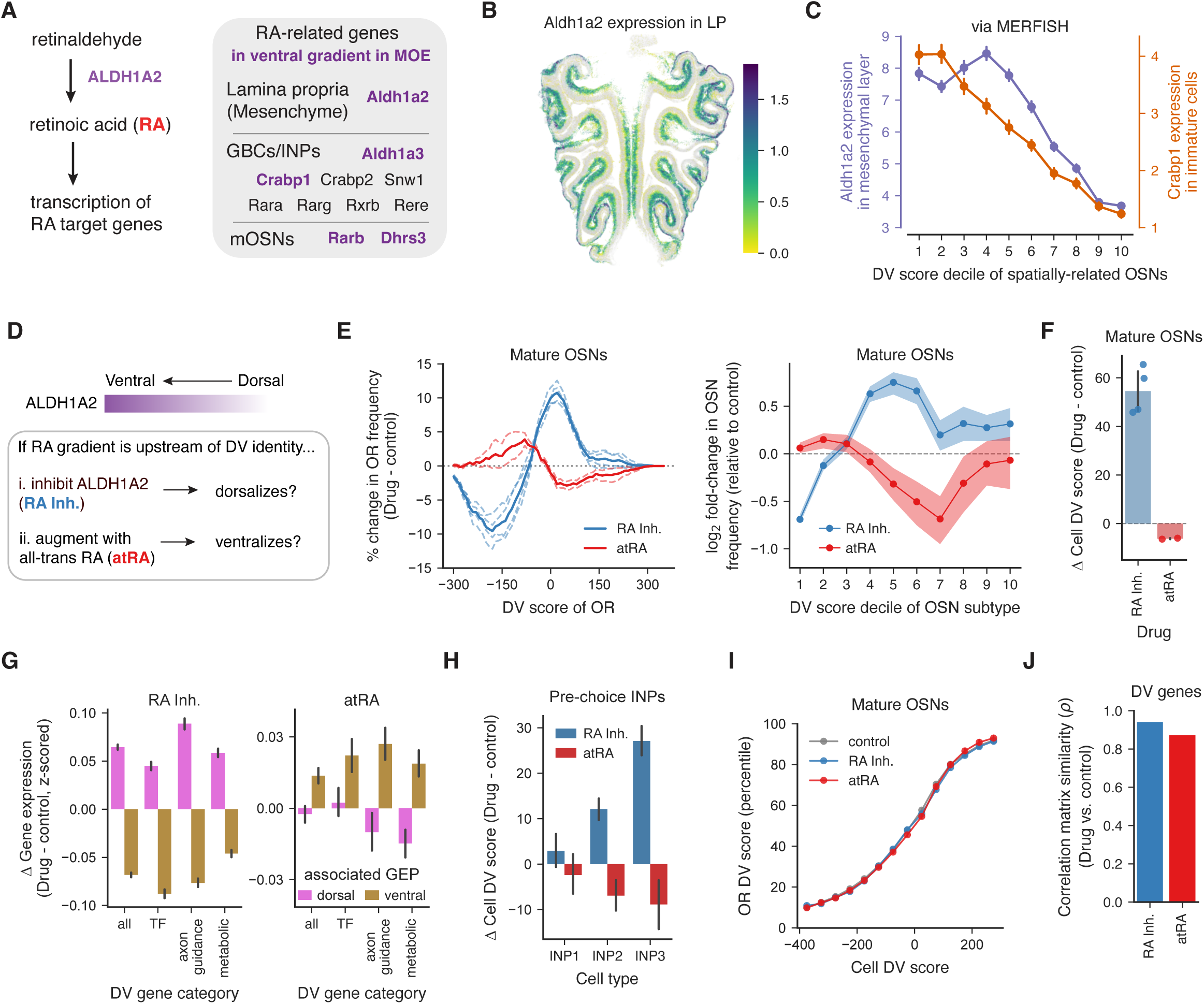
Manipulating retinoic acid signaling biases OSN fates and OR choice. A. (Left) Schematic of enzymes involved in retinoic acid signaling, and (right) examples of RA-related changes with expression in the MOE, with highlighted genes enriched in a ventral gradient. B. The expression of the retinoic acid-producing enzyme *Aldh1a2* in the mesemchymal lamina propria (LP) layer of the epithelium, as measured via MERFISH. C. The spatially-varying expression of *Aldh1a2* in the LP and the cellular retinoic acid binding protein *Crabp1* in immature cells, as a function of the DV score associated with the ORs expressed in spatially-related OSNs. D. Schematic depicting approach to manipulate RA signaling by systemically injecting mice with either the RA inhibitor (RA Inh.) WIN 18,446, which inhibits ALDH1A2, or all-trans retinoic acid (atRA) during OSN regeneration. E. (Left) Change in frequency of ORs with given DV scores relative to control mice injected with vehicle; note that data are obtained after epithelial regeneration post-methimazole treatment. Dashed lines indicate individual replicates. (Right) Log_2_ fold-change in OSN frequency among OSNs expressing ORs with given DV scores (grouped into 10 deciles). Error bars depict bootstrapped 95% confidence intervals. F. Change in the median DV scores (drug – control) for the OSNs from mice given the indicated drugs. Dots depict individual replicates and error bars depict bootstrapped 95% confidence intervals of the mean. G. The change in z-scored expression of the indicated DV genes, in animals treated with either drug, separated by gene category for DV genes associated with either GEP_Dorsal_ or GEP_Ventral_. H. Change in the DV scores for the pre-choice INP cells at the indicated stage. I. The relationship between percentile-normalized OR-associated DV scores and the DV scores of the OSNs themselves, for OSNs from the indicated conditions. J. The similarity in DV gene pairwise correlations between control animals and animals treated with either the RA inhibitor or atRA.

These findings raise the possibility that RA, originating in the mesenchyme, signals dorsoventral position and restricts cell fates in OSN precursors and mature OSNs. To test this hypothesis, we systemically delivered either an inhibitor of ALDH1A2 (WIN 18,446), or an activator of RA signaling (all-trans retinoic acid, atRA) to adult animals during methimazole-induced regeneration and evaluated their effects on the OSN cell identities and OR choices of regenerated cells via scRNA-seq (Figure 4D). As in the lineage tracing experiments, regenerated OSNs expressed single ORs (Figure S6D). However, compared to control mice, we observed bidirectional shifts in the number of “dorsal” or “ventral” OSNs: mice given the RA inhibitor had fewer OSNs expressing ORs with ventral DV scores and more cells expressing ORs with dorsal DV scores, whereas mice given atRA exhibited the opposite phenotype (Figures 4E and S6E). RA-dependent bidirectional shifts in DV scores — and their constituent DV genes — were observed in both precursors and mature OSNs (Figures 4F–H). Critically, these population-level shifts did not alter the relationship between DV scores and the chosen ORs *per se*, which were similar in normal and RA-manipulated epithelia, nor did they change the co-variation structure of the genes that make up the DV score itself (Figures 4I–J).

Sub-epithelial RA therefore informs overlying stem cells and precursors about their dorsoventral position, which is then translated into a particular pattern of DV gene expression and ultimately into the selection of a single OR that on average is optimal for a given spatial position; the fact that RA manipulations alter expression levels of all DV genes in concert — and thereby shift each of the ∼250 gene gradients up or down coherently — demonstrates that these genes are organized into a singular, rheostat-like, transcriptional program that is sensitive to spatial position.

### Dorsoventral location embodies a developmental timing signal for OR choice

How might physical space — presumably acting through RA and DV genes — influence the process of OR choice to create the ∼1100 overlapping and yet distinguishable peaks of receptor expression observed in the epithelium? One possibility is that space influences the palette of ORs that are co-expressed at low levels in precursors before choice ensues^32–34^. Consistent with previous observations, we find that OSN precursors often express at least ∼10– 20 ORs at low levels before ultimately choosing a single OR to express at high levels (Figures 3A, 5A and S4G). Quantification of the average DV score associated with the set of co-expressed ORs revealed that the process of OR co-expression and singular choice was temporally organized in a continuous dorsal-to-ventral fashion during differentiation (Figure 5B). We observed that when OR co-expression begins, it only includes a small subset of the dorsal-most receptors, which are expressed in all precursors regardless of spatial position (as inferred by the DV score of each cell). As differentiation proceeds, two processes happen in parallel. First, the dorsal-most precursors begin expressing the dorsal-most ORs at higher levels; as a result, individual dorsal ORs like Olfr1513 are only ever highly expressed (and subsequently chosen for singular expression) in a small number of dorsal cells (Figure 5B). Second, all of the precursors that are ventral to the dorsal-most set of precursors — which ultimately choose a dorsal OR — stop expressing the most-dorsal ORs and begin weakly co-expressing slightly more ventral ORs like Olfr194 (Figure 5B); as a result, the mean DV score associated with all of the co-expressed ORs at this stage is slightly more ventral. The next-most-dorsal precursor then begins similarly resolving its co-expression into a singular choice, and the process of co-expression, down-regulation and selection progressively repeats itself until the ventral-most ORs are finally co-expressed and subsequently chosen in the ventral-most cells.

**Figure 5.**
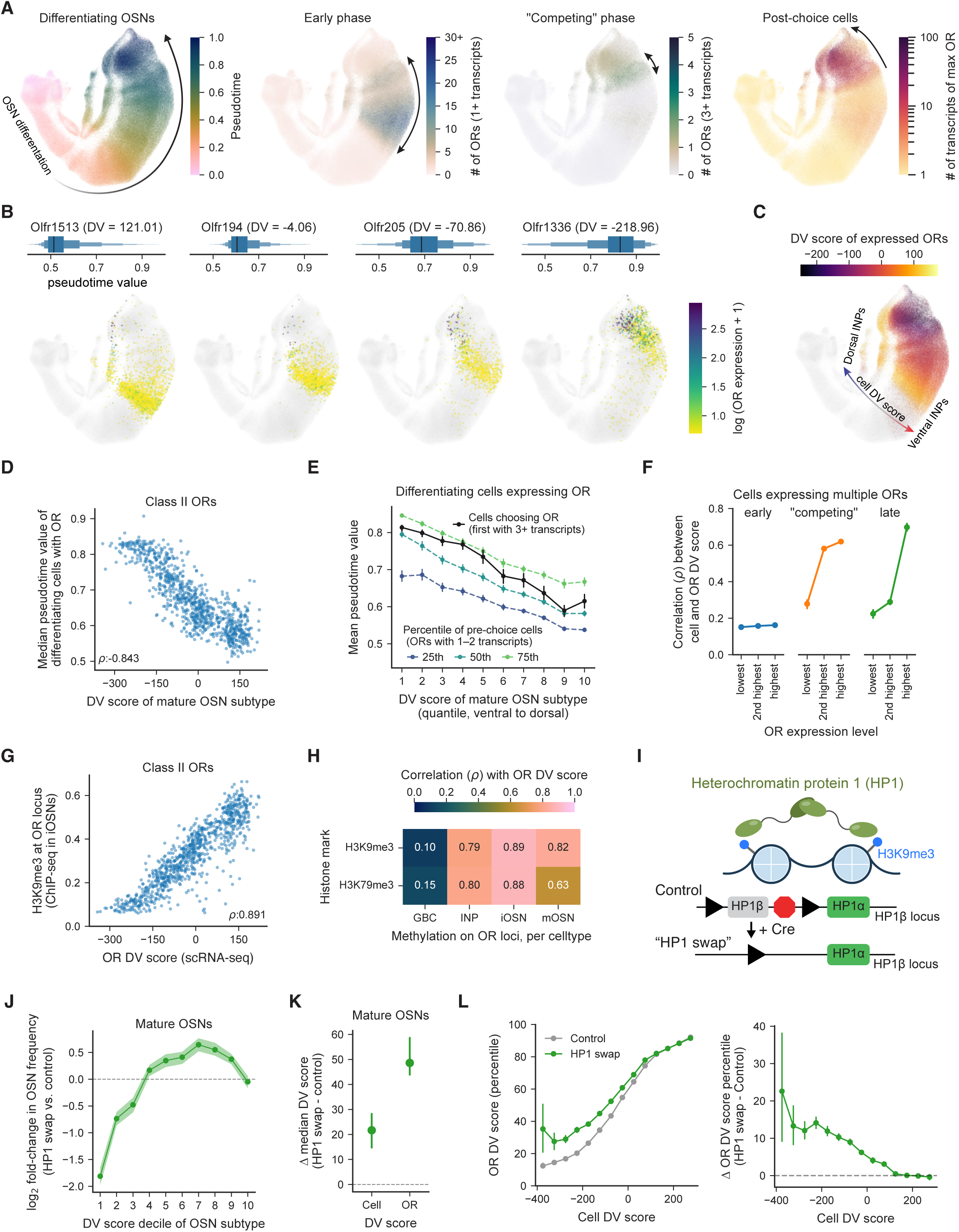
Space, time, and heterochromatin translate DV scores into restricted OR choice. A. OR expression in differentiating cells, captured at different phases of OR choice. (Left to right): the pseudotime of each cell, the levels of expression (measured as the number of receptor-associated unique molecular identifiers (UMIs) present in the scRNA-seq data) of the highest-expressed OR in each cell; the number of ORs expressed with at least one UMI; and the number of ORs with at least three UMIs. B. Examples of differentiating OSNs expressing the indicated ORs, colored by the log of the number of UMIs of each OR in each cell to depict pre- and post-choice. On top, the distribution of pseudotime values among the differentiating OSNs expressing each OR, and the mean DV score of mature OSNs expressing each OR). Cells not expressing each OR are colored gray. C. The mean DV score associated with all co-expressed ORs (as measured in the respective mature OSN subtype) in each cell. DV scores were weighted by OR expression for cells expressing multiple ORs, and cells not expressing any OR are colored gray. Note, in contrast to the OR-associated DV scores, cell DV scores vary across cells in an orthogonal axis present throughout OSN differentiation (see Figure 3E). D. The median pseudotime of OSNs expressing each class II OR (at any level of expression) during OSN differentiation, as a function of the DV score associated with mature OSNs that singularly express that OR. E. The mean pseudotime for cells expressing class II ORs, binned based on the DV scores associated with each OR in mature OSNs and plotted separately based on the expression level of each OR. For ORs expressed at the lowest levels (1–2 UMIs), the 25th, 50th, and 75th percentile of the pseudotime values of the cells expressing that OR are shown and compared to that for the first cells (10th percentile) expressing each OR at post-choice levels (3+ UMIs). F. For cells expressing multiple ORs, the correlation of the DV score for each immature cell with the DV score associated with the OR expressed at the highest, second highest, and lowest level; data are plotted separately for cells expressing multiple ORs at low levels, “competing” ORs at moderate levels, or those beginning to expressing a single OR at higher levels. OR-associated DV scores were determined based on scRNA-seq data from mature OSNs. G. The levels of the heterochromatin mark H3K9me3 measured via ChIP-seq at each Class II OR locus versus the DV score associated with each OR (as measured in mature OSNs via scRNA-seq). ChIP-seq data were previously generated and were reanalyzed32,101. H. The correlation between levels of the repressive heterochromatin marks (either H3K9me3 and K3K79me3) on each OR locus and the DV score associated with each OR (as measured in mature OSNs via scRNA-seq), for each of the indicated cell types. I. Schematic of the interaction between Heterochromatin protein 1 (HP1) and histones modified with the heterochromatin-promoting H3K9me3 mark, and depiction of the “HP1 swap” experiment, in which the loss of HP1β is rescued with HP1α; in these experiments HP1β was conditionally knocked out of neurons using the Foxg1-Cre mouse line. J. Log2 fold-change in OSN frequency among OSNs expressing ORs with given DV scores (grouped into 10 deciles) in HP1 swap relative to control mice. Error bars depict bootstrapped 95% confidence intervals. K. Change in the median DV scores relative to control animals for cells from the HP1 swap mice, and the change in the DV scores associated with the ORs (in control mature OSNs). Error bars depict bootstrapped 95% confidence intervals. L. (Left) The DV score associated with the OR expressed in each cell, plotted as a function of the DV score for the cell itself, from HP1 swap and control mice. (Right) Residuals from regression models predicting the DV score of each OR using the DV score of each cell. Models were trained on data from control animals and applied to HP1 swap animals.

In other words, we observe a sliding window of OR co-expression that systematically evolves as OSN precursors differentiate, with dorsal-most receptors being co-expressed first, and ventral-most receptors being co-expressed last (Figure 5C); this temporal pattern of co-expression is mirrored by a temporal pattern of receptor choice, in which dorsal precursors chose their final dorsal receptors earlier in pseudotime, while ventral precursors commit to their receptor choices later (Figure 5D–E). The delay between dorsal and ventral choice likely reflects the fact that ventral precursor cells must first co-express and silence more-dorsal ORs before subsequently picking a spatially appropriate ventral OR. Note that class I receptors located in the dorsal-most zone violate this pattern of co-expression and repression, consistent with class I OR-expressing OSNs having a distinct developmental trajectory (Figures 3B, S4G).

If indeed OR choice is restricted spatially, the final chosen OR in a mature OSN should ultimately reflect the underlying DV score of its pre-choice precursor. As expected, at an early differentiation stage (i.e., when dorsal ORs are broadly co-expressed) on average DV scores for each precursor and the DV scores associated with its expressed ORs (as determined in mature OSNs) are significantly mismatched. At a later differentiation stage, when many receptors are expressed at a low level and a few are expressed at a high level, the two highest-expressed ORs have associated DV scores that match each cell’s underlying DV score. Finally, at an even later stage of differentiation, when multiple ORs are still weakly co-expressed but only one receptor is expressed at a high level, the single highest-expressed OR is on average well aligned with the DV score (rho = .8, Figures 5F and S4H–I). Note that this correlation is high but not perfect, consistent with OSN precursors frequently choosing suboptimal ORs from a spatially limited distribution of possibilities. Thus, as OSN precursors transition from the multiple to single OR stage, expression levels of “incorrect” ORs systematically declines, ultimately yielding a mature OSN that chooses to express, on average, the OR that is most appropriate given its spatial position.

Importantly, RA-dependent shifts in dorsoventral cell identity (as captured by the DV score) influence the underlying process of OR choice, as RA manipulations that dorsalized (or ventralized) DV scores also dorsalized (or ventralized) the population of ORs that are co-expressed in pre-choice INPs (Figure S6F). Together, these data demonstrate that OSN development proceeds in a spatiotemporal gradient, with dorsal OSNs systematically maturing earlier and ventral OSNs maturing later during differentiation. Dorsoventral positioning therefore not only influences which of the ∼1100 different ORs a given mature OSNs chooses to express but also shapes the process of how OR choice unfolds during differentiation.

### Heterochromatin deposition translates DV scores into OR choice

The predictable relationship between mean DV scores and the chosen OR suggests that DV genes themselves somehow bias the process of OR choice. Previous work has suggested a mechanism in which the NFI transcription factors (NFIA/B/X) — which are included in the DV score — regulate heterochromatin-mediated repression of co-expressed ORs to facilitate OR choice^32,66^. This process is more prevalent in ventral OSNs, as they must sequentially repress the large suite of more-dorsal ORs before settling on an appropriate ventral OR to singularly express^37^. Recent experiments in which different “zones” of the epithelium were manually dissected revealed that mutation of NFI genes causes both overexpression of dorsal ORs in ventral zones and a reduction of heterochromatin in ventral zones, rendering their heterochromatin profiles akin to those from more-dorsal zones^32^.

Importantly, we find that *NFI* genes are expressed in a dorsoventral gradient in our MERFISH data (Figure S2B); exhibit graded expression in both precursors and mature OSNs with respect to DV scores (Figures 1C, S4B, and S4D); and are bidirectionally sensitive to RA manipulations (Figure S6G). These observations suggest that heterochromatin itself (which depends upon NFI activity) is likely also organized into a continuous dorsoventral gradient, one that in principle enables the precise levels of repression in each precursor required to achieve spatially appropriate OR expression. To test this hypothesis, we reanalyzed recent chromatin immunoprecipitation datasets to ask whether each OSN subtype’s DV score indeed predicted whether its cognate OR locus was heterochromatinized, and found a strong monotonic relationship (rho = 0.89) between the DV score associated with each OR and heterochromatin levels at that OR’s locus (Figure 5G); heterochromatin levels were highest for dorsal ORs because such receptors are co-expressed and repressed in nearly all cells, but they were lowest for the ventral ORs, which are only co-expressed in the most ventral cells. The relationship between OR location and heterochromatin was most apparent in immature OSNs in the process of choosing ORs, though similar results were observed in INPs and mature OSNs (Figure 5H). Thus — like OR expression patterns — heterochromatin levels of OR loci (which are controlled by a key DV transcription factor) are not organized zonally but instead systematically descend along a dorsoventral gradient.

If graded levels of heterochromatin regulate which OR is ultimately chosen at a given spatial position, then manipulations of heterochromatin might alter the typical relationship between spatial position and OR choice. One strategy for doing so is suggested by the finding that heterochromatinization of OR loci requires binding by the heterochromatin protein 1β (HP1β), which causally participates in compacting DNA into its repressed, heterochromatic form^67,68^. Because loss of HP1β leads to neonatal lethality, we performed scRNA-seq on “HP1 swap” mice, in which HP1β knockouts are rescued via the expression of HP1α, which also interacts with chromatin but is thought to play a less prominent role in transcriptional silencing (Figures 5I and S6H–I)^69,70^. Mature OSNs from HP1 swap mice expressed ORs at normal levels but were dorsalized relative to control mice, in that there were relatively more OSNs expressing ORs normally localized to the dorsal epithelium and fewer expressing ventral receptors (Figures 5J–K and S6J–L).

However, weakening heterochromatin also changed the relationship between the DV score and the chosen OR such that expressed receptors were more dorsal than would otherwise be expected given each cell’s DV score (Figure 5L and 6M). Similar effects were also observed in pre-choice INPs, where the distribution of weakly-expressed ORs at this stage was more dorsalized than the corresponding patterns of DV gene expression (Figure S6N). The observation that manipulations of heterochromatin alter the relationship between DV gene expression and OR choice demonstrates that heterochromatin is at least in part causally responsible for translating spatial position (as captured by the DV score) into a spatially appropriate distribution of ORs, suggesting a model in which variation in the expression of DV genes (e.g., *Nfia/b/x*) recruits different levels of heterochromatin and thereby implements distinct OR choices.

### Linked genomic context relates spatial positions to OR expression

While variation in heterochromatin levels provides a compelling means for suppressing the expression of spatially inappropriate receptors, it remains unclear how OSN precursors facilitate OR co-expression and ultimately choose single ORs in a manner that respects dorsoventral positioning. Given the spatiotemporal gradient in OR co-expression observed in our atlas of differentiating cells, one would predict that binding sites for transcription factors known to drive OR expression might vary as a function of spatial position. Indeed, ORs with increased number of binding sites for EBF transcription factors (which are required for OR expression) were more likely to be dorsal, consistent with the early expression of dorsal ORs during OSN differentiation (Figure S7A). In contrast, OR genes with binding sites for NFI and RAR transcription factors were more likely to be expressed ventrally, consistent with the graded expression of both *NFI* and RA-related genes observed in ventral OSNs (Figure S7A).

These observations suggest that local genomic context (i.e., upstream and promoter sequences) assigns each OR to its appropriate dorsoventral position in the epithelium. Indeed, analysis of “receptor swap” mice, in which the coding sequence of one receptor replaces that of another receptor at a distinct genomic location^25^, revealed that OSN DV scores (and expression levels of *NFI* and other DV genes) reflect the genomic locus in which the expressed OR resides, whereas activity-dependent gene expression depends upon the specific OR protein being expressed (Figures S7B–E). This observation is consistent with a model in which cells acquire a spatial identity reflected by the expression of DV score genes (as the OR does not determine the DV score), and in which a given OR is rendered competent for expression within precursors of a particular DV score by sequences linked to the OR in the genome. Note that differences in the DV scores associated with any two ORs only weakly relate to OR phylogenetic relationships or how close each OR is to the other in the genome (Figure S7F–I).

To further test the idea that cis-regulatory elements determine the dorsoventral positioning of individual ORs, we characterized DV scores in OSNs from F_1_ hybrid crosses between C57BL/6J mice and wild-derived CAST/EiJ mice (Figure 6A). These two mouse strains have largely homologous sets of ORs but many single nucleotide polymorphisms (SNPs) across their genomes^71^; such SNPs might be expected to occasionally influence enhancers and promoter elements relevant to OR choice, thereby yielding an OR that is, for example, expressed dorsally in one strain and ventrally in the other. As expected, OR choice was monogenic and monoallelic (Figure 6B), thus allowing us to compare the GEP usages of OSNs expressing each OR from each strain in a single F_1_ hybrid mouse. While GEP usage was largely consistent within a strain for all OSN subtypes, a subset of OSN subtypes (and therefore a subset of associated OR alleles) had significant differences in their associated DV scores across strains (Figures 6C and S7J–L); on average there are 8 SNPS in the upstream regions of the OR genes whose associated DV scores differed across strains. Importantly, OR coding sequence variation was unrelated to strain-specific differences in DV scores but instead predictably altered the use of previously-identified OSN gene expression programs that report ligand-dependent and -independent OR activity (Figures 6D and S7J)^54^.

**Figure 6.**
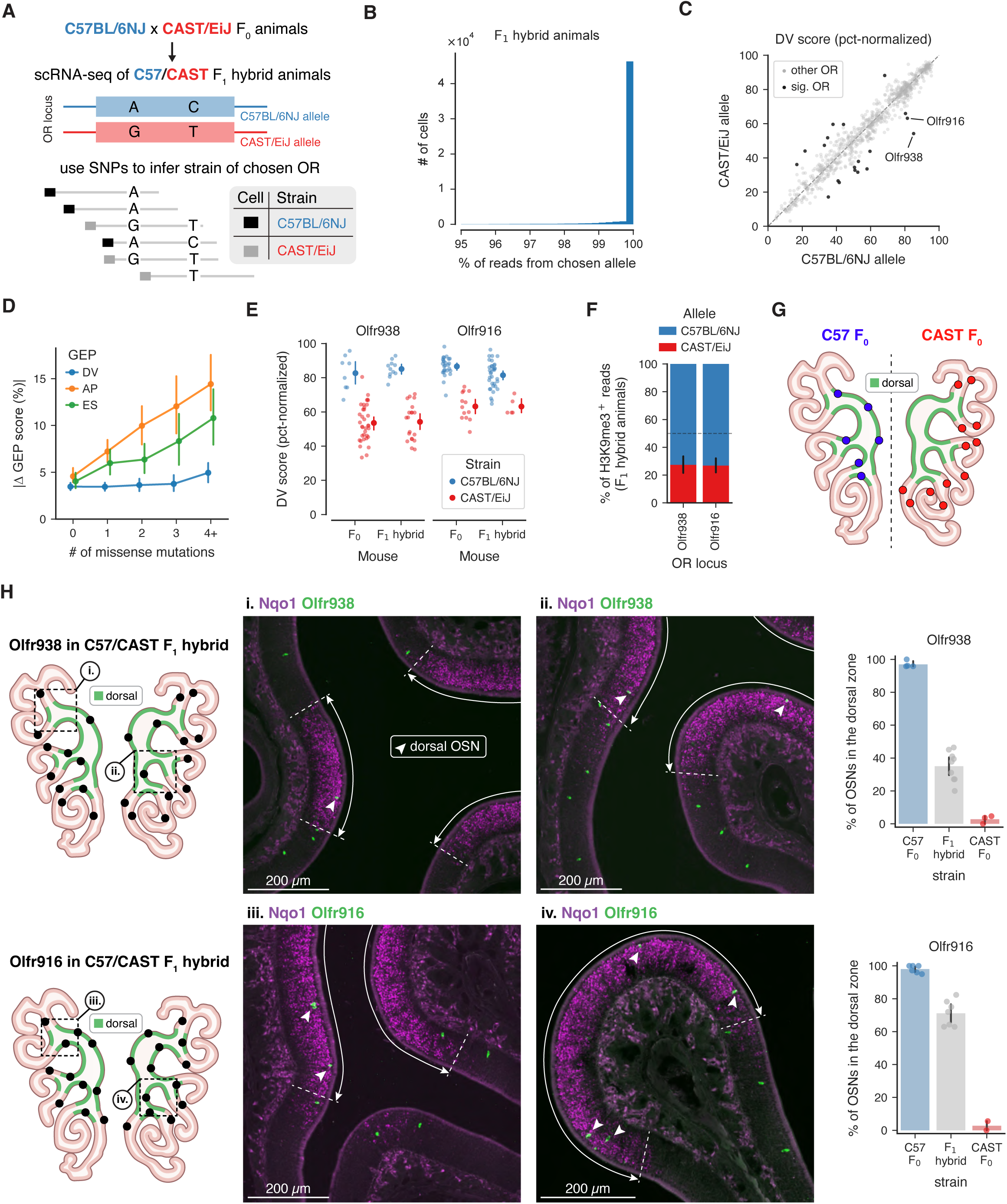
OSN spatial locations are controlled by genomic context. A. Experiment schematic depicting how the strain of the chosen OR was identified for individual OSNs from F_1_ hybrid animals, which were generated by crossing C57BL/6J and wild-derived CAST/EiJ mice. B. For mature OSNs expressing single ORs, the percent of reads that came from the chosen OR allele, out of all reads that mapped to either allele. Note the truncated x-axis (mean 99.6%). C. The DV score for each OSN subtype, evaluated separately for OSNs from F_1_ animals that chose either the C57BL/6J or CAST/EiJ allele of its cognate OR. OSN subtypes with significant changes in DV scores across strains, including OSNs expressing either Olfr938 or Olfr916, are highlighted in black (see Methods). D. The absolute change in percent-normalized GEP usages for either the DV or other activity-dependent GEPs for OSN subtypes whose ORs had the indicated number of missense mutations in their coding sequences. Error bars depict bootstrapped 95% confidence intervals across OSN subtypes. ES, environmental state activity GEPs; AP, anterior-posterior GEPs; usage of the ES and AP GEPs reflect differences in ligand-dependent and ligand-independent OR activity, respectively^54^. E. DV scores (percent-normalized across all OSNs) for individual OSNs expressing Olfr938 and Olfr916, separated by strain for F_0_ animals and based on the inferred strain of the chosen allele in F_1_ animals. Error bars depict bootstrapped 95% confidence intervals across OSNs. F. Differences in the levels of heterochromatin at the Olfr938 and Olfr916 loci in F_1_ animals (measured at the percentage of receptor reads using the CUT&RUN technique with antibodies against histones marked with H3K9me3); increased levels of heterochromatin in C57 compared to CAST alleles is consistent with a more-dorsal epithelial location. G. Schematic depicting the results of in situs for Olfr938 and Olfr916 in either C57BL/6J or CAST/EiJ F_0_ animals, with the dorsal region of the MOE in green. H. (Left) Schematic depicting the positions of OSNs expressing either Olfr938 or Olfr916 in F_1_ hybrid animals. (Middle) Insets showing in situs in the highlighted regions at the boundary between the dorsal and ventral regions of the MOE, with the indicated OR in green, the dorsal marker Nqo1 in magenta, the dorsal zone outlined in white, and dorsal ORs labeled with arrowheads. (Right) Summary quantification of the percentage of OSNs found in each animal that were located within the dorsal zone, with dots depicting individual replicates and error bars depicting bootstrapped 95% confidence intervals.

We wondered whether strain-specific differences in DV scores reliably predicted dorsoventral position differences in the epithelium. To address this question, we performed in situ hybridizations for *Olfr938* and *Olfr916*, two ORs with higher DV scores and correspondingly higher levels of heterochromatin in C57BL/6J mice compared to those in CAST/EiJ mice (Figures 6E–F). Consistent with the observed differences in their DV scores, *Olfr938* and *Olfr916* cells were exclusively located within the *Nqo1*^+^ dorsal region of the epithelium in C57BL/6J mice but were found in the *Nqo1^-^*ventral region in CAST/EiJ mice (Figure 6G). Furthermore, in situs for Olfr938 and Olfr916 in the F_1_ hybrids revealed that the same OR was found both within and outside of the *Nqo1*^+^ dorsal region— effectively residing in two “zones” simultaneously — matching the spatial positions of the two respective F_0_ animals (Figure 6H). Together, these receptor swap and F_1_ hybrid data argue that the spatial position of each OR is governed by noncoding elements in its genomic locus and suggest a model in which differential binding of transcription factors (e.g., NFI, RAR) whose presence or activity varies along the dorsoventral axis qualifies each OR for expression at particular positions in epithelial space, i.e. those positions with the appropriate DV scores.

### Epithelial spatial positions coordinate axon targeting to the olfactory bulb

The olfactory bulb contains a precise and stereotyped map in which each OSN subtype typically projects to one medial and one lateral glomerulus. The DV score represents a potentially parsimonious mechanism for coupling the expression of individual ORs (which are organized in epithelial space) to the expression of genes that guide OSN axons to their cognate glomeruli (which are organized in bulb space). Indeed, the DV GEPs include many genes that could facilitate axon guidance; while the DV genes *Robo2* and *Nrp2* are known to regulate dorsoventral glomerular targeting by OSN axons^12,44–46^, many genes in GEP_Dorsal_ and GEP_Ventral_ have not previously been implicated in OSN axon guidance and yet are organized into dorsoventral expression gradients, including cell surface molecules like *Ncam1*, *Mdga2*, *Plxna4*, *Plxnb3*, *Cdh3*, *Itgb1*, cytoskeletal regulatory molecules like *Rhoq*, *Cyth1*, *Septin3*, and at least a half dozen additional small GTPases or GTPase regulators (Figures 1C, S4B, Supplemental Table 1). Importantly, expression levels of *Nrp2* and the other DV-associated axon guidance genes were bidirectionally altered upon manipulations of RA signaling (Figure 4G), suggesting that their expression is fundamentally coupled to the gradients that define epithelial space.

OSN DV genes might therefore represent an organized transcriptional program that coordinates continuous variation in the spatial position of cell bodies in the olfactory epithelium with the elaboration of axons targeting specific glomerular positions in the olfactory bulb. To address this possibility, we took advantage of a recent spatial transcriptomics dataset in which the OSN subtypes that innervate each glomerulus were identified^72^. Coloring the identified glomeruli based on the DV score of its associated receptor revealed that DV scores varied smoothly across bulb glomeruli (Figure 7A); as a result, neighboring glomeruli had similar DV scores (Figures 7B–C). Furthermore, the 3D glomerular position of each glomerulus predicted the DV score of the innervating OSN subtype with high accuracy (rho=0.95, Figure 7D), and conversely the DV score of each OSN subtype predicted the 3D location of the innervated glomerulus with low median error (∼400 uM, 3–4 glomerular widths, Figure 7E). Thus spatially-organized variation in DV gene expression across OSN subtypes can at least in part explain the spatial organization of their projections to the bulb.

**Figure 7.**
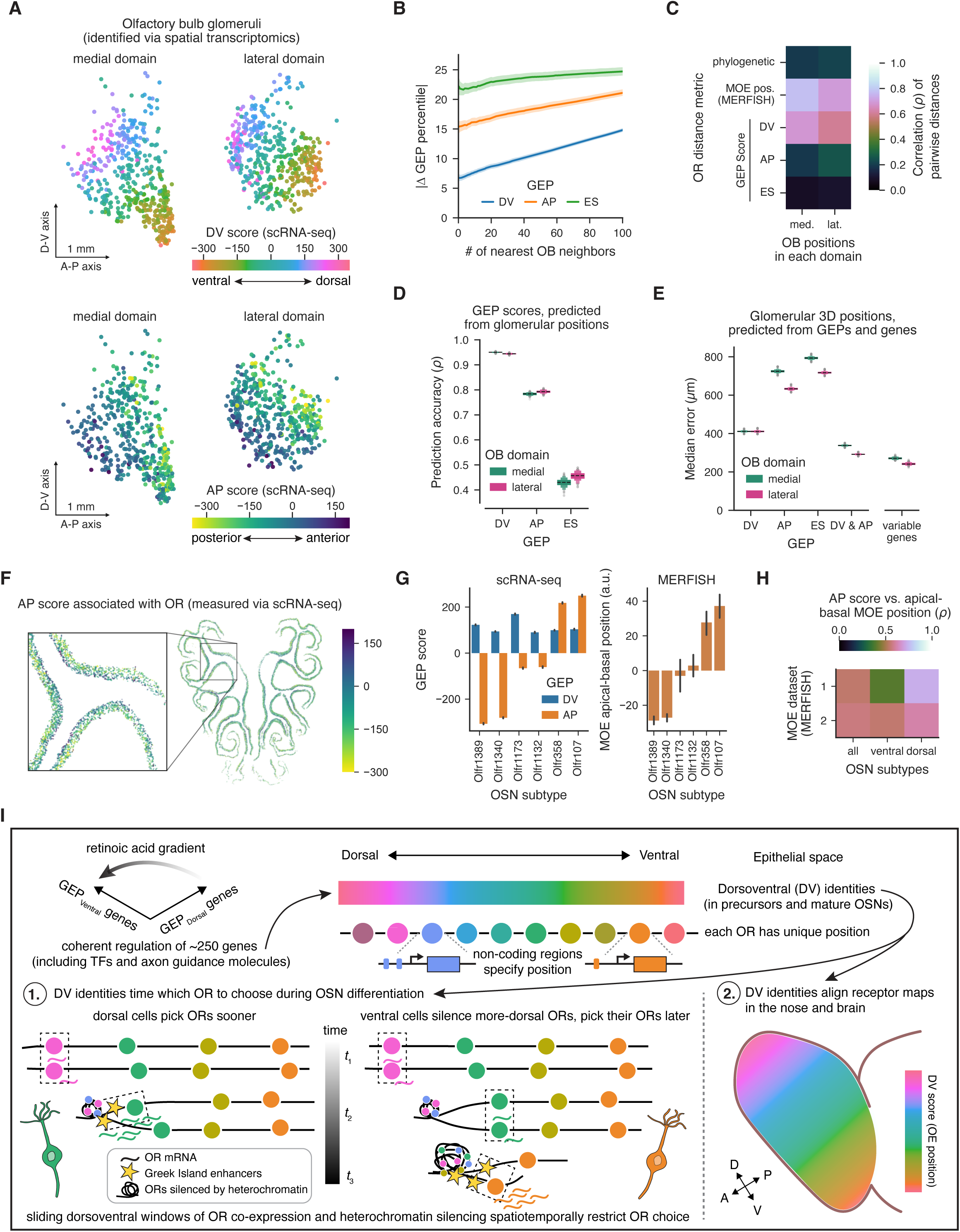
Olfactory bulb glomerular positions reflect epithelial DV scores. A. Glomerular positions in the medial and lateral domains of the olfactory bulb (OB), colored by the respective (top) DV and (bottom) AP scores associated with each OSN subtype. The OSN subtypes associated with each glomerular position was identified via a recent spatial transcriptomics dataset72. B. The average pairwise distance in GEP usage for the DV, AP, and ES GEPs among OSN subtypes located within the given number of neighboring glomeruli in the OB. ES, environmental state activity GEPs; AP, anterior-posterior GEPs; usage of the ES and AP GEPs reflect differences in ligand-dependent and ligand-independent OR activity, respectively54. C. Correlations between pairwise distances (across all OSN subtypes) of OB glomerular positions (measured as 3D distances in the indicated medial or lateral OB domain) and GEP usages for the DV, AP, and ES GEPs; MOE epithelial positions; and the phylogenetic distances of their respective OR proteins. D. The accuracy of regression models to predict the DV, AP or ES GEP scores using 3D glomerular positions. E. The accuracy of regression models predicting 3D glomerular positions using either the indicated GEPs or all 1,300 OSN variable genes. F. Example MERFISH section colored by the AP score (as measured via scRNA-seq) of each OSN (identified via the expressed OR, Data from Bintu et al.) G. (Left) DV and AP GEP usages for the indicated dorsal OSN subtypes. (Right) Apical-basal position of the OSNs expressing each OR. Error bars indicate means and 95% confidence intervals across OSNs. H. Correlation of the AP score for each OSN subtype with its apical-basal position as measured in two separate MERFISH datasets (1: this manuscript and 2: Data from Bintu et al.), for either all detected OSN subtypes or for those that were identified as either dorsal or ventral based on their DV scores. I. Summary of the results, depicting how RA signaling varies spatially within the OE to yield 1,000 different DV identities and OR positions, and how these DV identities coordinate both OR choice during OSN differentiation and the specific projection patterns of each OSN subtype to the OB.

We therefore wondered whether additional spatially organized gene expression programs might also participate in correctly guiding axons to glomeruli. In previous work we identified an additional pair of GEPs (GEP_Anterior_ and GEP_Posterior_, ∼180 total genes)^54^, which includes genes (like *Nrp1* and *Plxna1*) known to mediate axon targeting along the anteroposterior (AP) axis of the olfactory bulb^49,50,73^; these GEPs include many other additional genes relevant to axon guidance and cytoskeletal dynamics that could in principle participate in this process, including *Plxna3*, *Sema6c*, *Pcdh7*, *Cdh15* and *Nexn*. AP GEPs are used by OSNs in a mutually exclusive fashion, enabling the generation of an “AP score” that varies across subtypes, but unlike the DV GEPs, AP GEPs are only used by mature OSNs expressing functional ORs (Figure S4F)^54^. We find that the use of the AP GEPs smoothly varied across olfactory bulb glomeruli, although along an orthogonal spatial axis to that used by the DV GEPs (Figure 7A). However, the relationship between AP scores and glomerular positions was less predictive and more error-prone than that for DV scores (Figures 7D–E). Nevertheless, combining DV and AP GEPs predicted glomerular 3D positions with an error of only ∼300 µm, similar to the noise in the spatial position estimates themselves (Figure 7E, see Methods); this high level of performance was also similar to that observed for models trained with the entire set of ∼1,300 OSN variable genes, indicating that the DV and AP GEPs together effectively summarize the relevant transcriptional axes that map OSNs onto bulb position.

Given that DV scores explain OSN positions along the dorsoventral axis of the epithelium, we further wondered whether AP GEPs usage varies spatially across the epithelium. Surprisingly, the apical-basal positions of OSN subtypes within the epithelium systematically varied depending on their AP gene expression (Figures 7F–H, and S7M). The relationship between apical-basal position and AP scores was strongest for dorsal OSN subtypes (Figure 7H), which occupy precisely the bulb region in which epithelial position is least informative about glomerular position (Figures 2B, 7A, and S2J). Thus, the two cardinal axes that organize the olfactory bulb (DV and AP) are both topographically and transcriptionally represented in the epithelium itself.

## Discussion

The peripheral olfactory system has long been thought to balance determinism and randomness by limiting the expression of individual ORs to one of a handful of dorsoventral zones, while randomly distributing ORs within each zone^18,19^. Here we show instead that the olfactory epithelium harbors an organized map in which each of the ∼1100 receptors reliably and predictably occupies one of ∼1100 peaky (and therefore distinguishable) dorsoventral distributions along the dorsoventral axis of the epithelium. This pattern of expression reflects a developmental program in which spatial RA gradients are transformed into spatial transcriptional gradients in more than ∼250 genes, all of which are co-regulated and organized into a coherent transcriptional axis. This transcriptional axis — summarized as the DV score — allows us, for the first time, to clearly relate epithelial space to the key events during the differentiation and maturation of OSNs that ultimately lead to singular OR choice. We find that the differential expression of DV genes by OSN precursors correlates with the timing of receptor co-expression over pseudotime, the extent to which OR loci are heterochromatinized, and the identity of the chosen receptor. The relationship between DV gene expression and OR choice systematically varies after RA manipulations, demonstrating that RA spatial gradients play an important role in specifying OSN fates. Ultimately, heterochromatinization of OR loci — which are regulated by the DV genes NFIA/B/X — implement the transformation between spatial identity and OR choice, as weakening heterochromatin causes a precursor with a given DV score to choose from a different distribution of receptors than it would otherwise. The same transcriptional axis that builds a spatial map of OR identity in the epithelium likely coordinates the alignment of the receptor map in the nose with that in the bulb, as it includes a host of axon guidance-related genes and accurately predicts glomerular positioning. Together, these results demonstrate that precursors and mature OSNs harbor spatially specific transcriptomes, which both diversify and organize the peripheral olfactory system (Figure 7I).

Our data reveal, somewhat surprisingly, that development can solve the problem of reliably organizing ∼1100 different sensory channels in space. While relationships between OSN spatial positions and OR choice are highly predictable at the population level, they are noisy at the single neuron level, although the precise distribution of receptors from which a given precursor can select is tightly regulated by its spatial position. It is possible that the different alternative OR choices implemented by precursors of a given DV score reflects stochastic variation in interactions between OR enhancers located on different chromosomes and OR gene promoters, as recent work has demonstrated that different cells expressing the same OR have distinct nuclear architectures^74–78^. We find that dorsoventral position influences both the timing of OR co-expression and heterochromatin formation, enabling spatially appropriate OR choice. We speculate that the variability of the translation between space and OR choice may in part reflect noisiness in this timing mechanism; consistent with this possibility, the diversity of DV scores associated with each OSN subtype systematically increases in more ventral cells, suggesting that the accumulation of small timing errors may broaden the diversity of co-expressed receptors available for choice for ventral precursors (Figure S1I). Even though feedback processes are likely required to stabilize the singular expression of each functional OR, our observations suggest that OR choice is an open loop process that does not evaluate whether the spatially-optimal OR was chosen in each cell^25,48,59,74,79^.

What ultimately limits the expression of a particular OR to a given peaked dorsoventral distribution in the epithelium? Our analysis of OR promoters suggest biases in the presence of transcription factor binding site that may amplify the influence of spatially-organized gradients of transcription factor expression or activity (e.g., NFIA/B/X and RAR) to tune each OR’s propensity to be expressed and/or silenced pre-choice^79^ in precursors of a particular DV score, which ultimately allows OR choice to systematically vary as a function of spatial position. Although further work is required to dissect the *cis*-regulatory logic that links a given OR to the expression of DV genes reflecting its position in space, our results from F_1_ hybrid animals demonstrate that genomic sequences — rather than OR coding sequences — largely determine epithelial positioning, extending past work showing OR promoters can confer spatially restricted transgene expression^80–83^, and that the genomic locus from which an OR is expressed is related to glomerular targeting location in the olfactory bulb^25,84^. The broad mechanism we propose here — in which transcription factor binding sites upstream of each OR reads-out spatially-driven variation in transcription factor expression and activity — is reminiscent of the transcriptional logic that gives rise to the restricted combination of Drosophila *Or* (*dOR*) genes expressed in each sensillum^85,86^. However, the variability at the single cell level of the relationship between spatial position and mammalian OR choice observed here stands in contrast to the perfect determinism that characterizes *dOR* gene expression in the fruit fly.

We find no clustering or ordering of ORs in epithelial space based upon the particular odors to which they respond (with the notable exception of dorsal responses for acids). Thus, while there is a receptor map in the nose, its function is fundamentally unlike that of those in other sensory systems, where topographical organization relates directly to sensory tuning. Why, then, is receptor position spatially organized in the nose? We propose that the epithelial map is a necessary consequence of a developmental mechanism coordinated by DV genes whose function is to match OR choice to axon guidance programs that specifically target a given glomerular location in the bulb. Because DV genes bias but do not perfectly determine the OR chosen by a given OSN, OSNs expressing the same OR are distributed across locally overlapping regions of the epithelium, which demands OR-dependent axon guidance mechanisms (i.e., AP score genes, OR-dependent homophilic attraction and repulsion mechanisms) to reconcile variation in epithelial dorsoventral position with precision in glomerular locations. However, this cost is accompanied by a likely benefit, as the spatial distribution of each OSN subtype may contribute to the robustness of the olfactory system by rendering individual ORs resistant to local insults to the epithelium.

It has previously been demonstrated that precursors transplanted from the dorsal to ventral epithelium generate OSNs that express ventral markers and ORs, suggesting that spatial determinants influence OR choice^15^. Here we take advantage of pharmacology to identify RA as one such determinant, and further show that RA controls the coordinated expression of DV genes; although RA had been previously proposed to signal information about physical space (given the graded expression of e.g., ALDH1A2), testing this hypothesis has been challenging given the pleiotropic roles played by individual RA-related genes in olfactory stem cells^12,65,87–89^. Indeed, our data suggest that the function of many individual genes in OSN differentiation and fate determination may be obscured by either redundancy or pleiotropy, and that instead subtle variation in the coordinated expression of many or all of the ∼250 genes making up the DV score underlies e.g., OR choice and glomerular targeting. Future work will be required to determine how RA influences the graded levels of expression apparent in each of the DV genes and to isolate to specific cell types in which it acts, to address the possibility that in adult tissues there are additional spatial determinants that influence choice, and to explore the relationship between RA and developmental mechanisms that organize the epithelium during its initial ontogeny during embryogenesis^90,91^.

It is important to note that the dorsal-most region of the OE may indeed constitute the one *bona fide* “zone” in the nose, as it follows different rules than the remainder of the epithelium (i.e., it expresses unique marker genes, uses and harbors a variety of distinct, transcriptionally-defined precursor types that give rise to mature OSNs expressing not just Class II ORs, but also Class I and TAAR-type receptors^58,92,93^, and appears less sensitive to RA manipulations than the ventral epithelium); additional work will be required to assess how DV genes differentially influence fates among OSNs expressing non-canonical OR types like the TAARs and those few ORs that are organized into epithelial patches^94–96^. We note that the NFI transcription factors, which are absent in the dorsal epithelium, are essential for heterochromatin formation and appropriate OR choice in the ventral epithelium^32^; it may be that their absence contributes to the relatively broad spatial distributions of dorsal zone receptors.

Recent work has demonstrated that, over long timescales, odor experience can influence the number of mature OSNs present in the nose that respond to an experienced odor^97–100^. However, given a model in which OSN precursors randomly chose their receptors, it is not clear how odor responses in mature neurons might somehow bias OR choice towards the required receptor in stem cells, which themselves cannot detect odorants. Our observation that OSN precursors of a given DV score and therefore spatial identity sit immediately underneath the more mature OSNs to which they give rise suggests possible mechanisms through which such a coupling could be achieved. We speculate that odor responses in mature OSNs generate a signal that increases proliferation and differentiation of underlying stem cells, which would be expected to generate a population of mature OSNs whose expressed OR reflects their shared dorsoventral position in the epithelium. Such a signal could emanate from mature OSNs and diffuse to the stem cell layer, or could be propagated indirectly via adjacent sustentacular cells, which span the full height of the epithelium. Given our data that single OSN precursors give rise to more than one OSN subtype, a main prediction of this model is that the odor experience will recruit not just OSNs that respond to the experienced odors, but also OSNs that have similar DV scores but express receptors for odors different than those that were experienced; future work will be required to test this hypothesis.

## Acknowledgements

We thank Catherine Dulac for CNGA2 knockout mice, and members of the Datta lab for helpful feedback on the manuscript. We thank Jingwen Araki and Julia Martz for laboratory assistance, and Euipoom Yoon for illustrations. We acknowledge the Bauer Core Facility at Harvard University, the Flow Cytometry Core and Spatial Technologies Unit (STU) at Beth Israel Deaconess Medical Center, and the Neurobiology Imaging Facility (NIF) and Microscopy Resources on the North Quad (MicRoN) cores at Harvard Medical School. Portions of this research were conducted on the O2 High Performance Compute Cluster at Harvard Medical School. SRD is supported by R01DC021669, R01DC021422 and funding from the Tan-Yang Center. DHB was supported by an NSF Graduate Research Fellowship and by NIH grant F31DC019017.

## Author Contributions

D.H.B, T.T., and S.R.D conceived and designed the study. D.H.B and T.T. performed the experiments. C.T. and D.K. helped perform the F_1_ hybrid experiments, and R. L. and L.T.K helped perform the clonal lineage tracing and RA experiments, respectively. N.K. and M.K. provided the olfactory bulb spatial transcriptomics data, M. E-D-A. provided the HP1 swap mice, and B.B. generated an additional MERFISH dataset. S.R.D. acquired funding and supervised the study; T.B generated the swap mice and provided additional supervision. D.H.B and S.R.D wrote the manuscript, with input from the other authors.

## Supplemental figure legends

**Figure S1.**
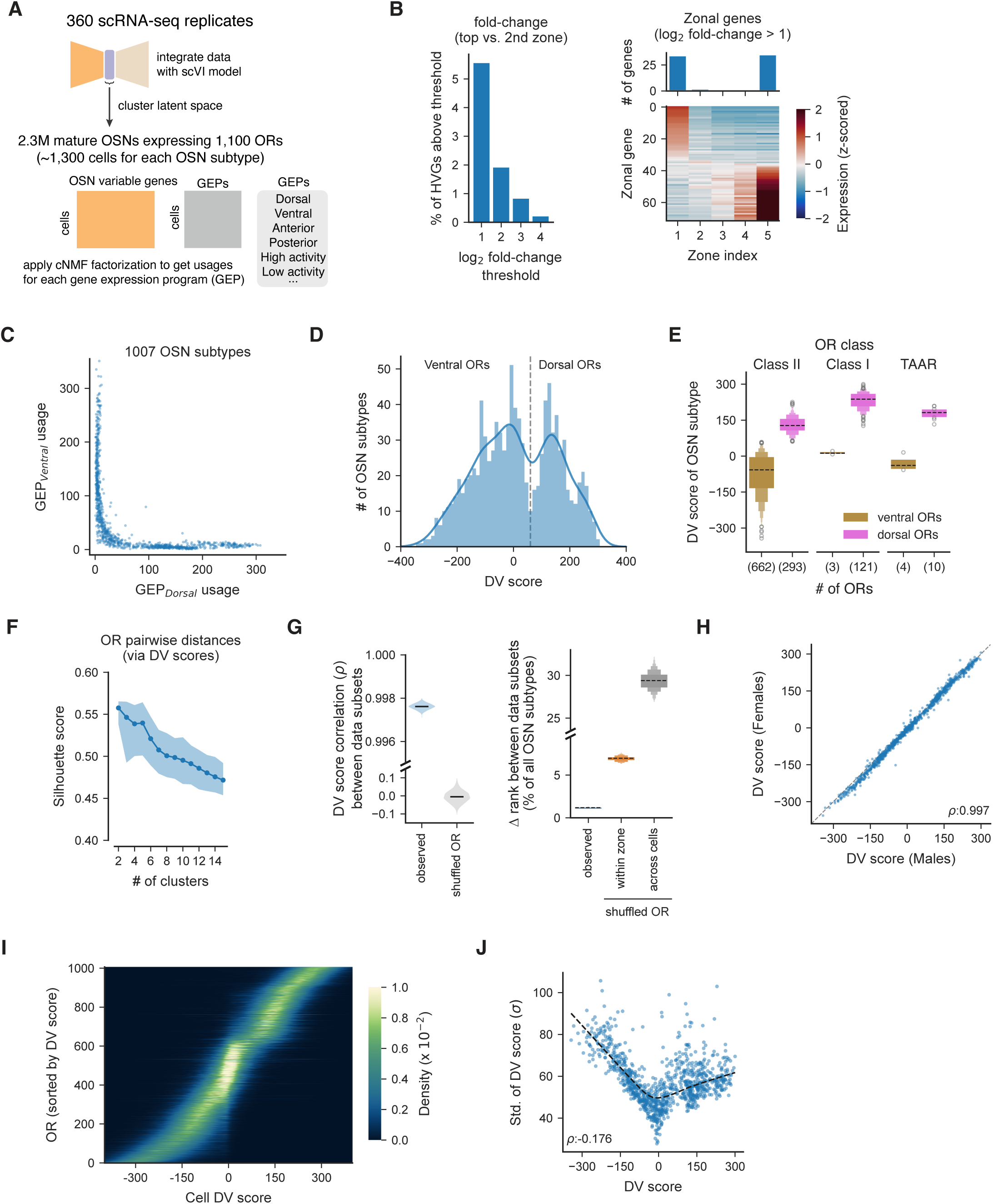
DV scores vary across OSN subtypes in a stereotyped manner, related to Figure 1. A. Schematic depicting approach for integrating single-cell RNA-sequencing (scRNA-seq) data from hundreds of replicates. Mature OSNs were identified by clustering the latent space of a scVI variational autoencoder model, and gene expression programs (GEPs) were identified using consensus non-negative matrix (cNMF), as previously described^54^. B. (Left) The fraction of the ∼1,300 OSN highly variable genes (HVGs) that showed enriched expression (log_2_ fold-changes in the top zone over the second-highest zone above the indicated thresholds, thresholded as indicated on the X axis) for OSN subtypes in a single zone, using discretized zonal indices, as previously inferred for each OR^21^. (Right) The z-scored expression across OSN subtypes of zonally-restricted HVGs (using a threshold of log_2_ fold-change > 1). C. The mean usage of GEP_Dorsal_ and GEP_Ventral_ by each OSN subtype (defined as the set of OSNs singularly expressing a given OR), for the 1007 OSN subtypes with at least 150 cells in our dataset. D. Histogram depicting the distribution of mean DV scores across OSN subtypes (after DV scores were binned into 61 equal-width bins). OSN subtypes were categorized as dorsal if their mean DV score was greater than 60 (see Methods). E. The distribution of DV scores for each OSN subtype, across different OR classes and separated for OSN subtypes that were categorized as dorsal or ventral (with the number of ORs in each category indicated); TAARs = Trace amine-associated receptors. F. Silhouette scores (quantifying clustering quality) from hierarchical clustering of the pairwise distance matrix of DV scores between OSN subtypes (number of clusters indicated on the x-axis, dark line and shading indicates the mean and 95% confidence of the mean across restarts in which equal numbers of OSNs were sampled per subtype). G. (Left) The correlation of OSN subtype DV scores across different subsets of cells belonging to that subtype, compared to the correlation when subtype labels were shuffled across cells; subsets generated by randomly sampling 150 cells, splitting the sample into two subsets, and computing the correlation. (Right) The mean absolute difference in ranks of the DV score for the data on the left, plotted as a percentage of a total of number of OSN subtypes, and compared to a model in which ORs were shuffled across cells of a given zone. H. DV scores for each OSN subtype, as measured in male and female mice. I. Heatmap depicting, for each OR, kernel density estimates of the distribution of DV scores across the set of OSNs expressing that OR. J. The standard deviation in the DV score for each OSN subtype, as a function of each subtype’s DV score.

**Figure S2.**
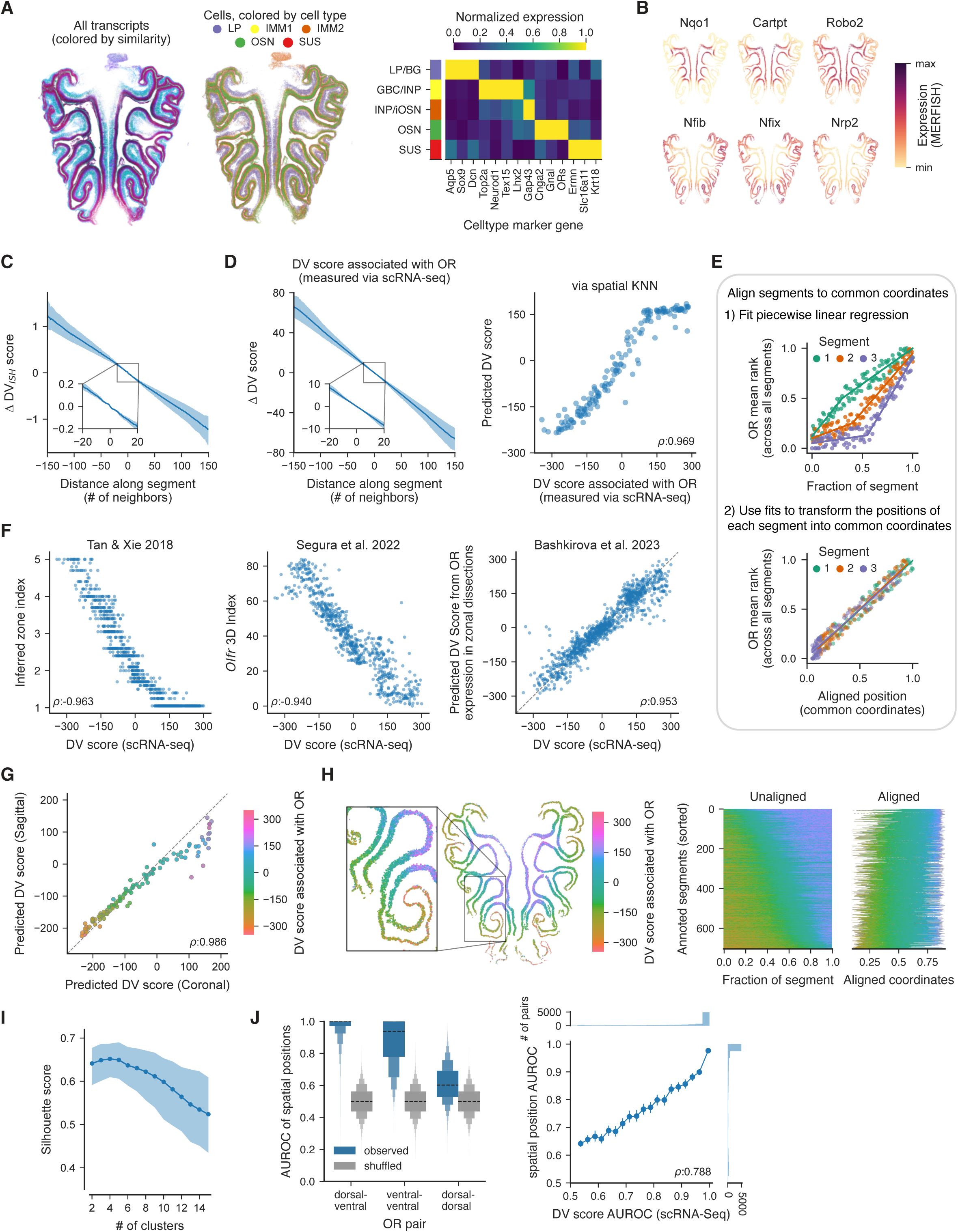
DV scores predict spatial positions across datasets, related to Figure 2. A. (Left) All transcripts from the MERFISH experiment, aligned to a common template and colored by their transcriptional similarity via Baysor. (Middle) Cells, segmented via Proseg, clustered based on their gene expression, and colored by their corresponding cell type labels. LP, lamina propria. IMM, immature cells (GBCs, INPs, and iOSNs). (Right) Heatmap of max-normalized marker gene expression, as measured via MERFISH, for each cell type. B. Expression of the indicated example DV genes, as measured via MERFISH. C. (Left) The observed change in DV_ISH_ scores (i.e., the estimate of the DV score obtained from a subset of the DV genes that were queried via MERFISH) as a function of changes in spatial position within an epithelial segment. Shading is mean ± 95% confidence intervals across epithelial segments. Pullout demonstrates the continuity of changes in the DV_ISH_ scores as a function of physical space. D. (Left) Similar to (C), but depicting the change in DV scores of the OSN subtypes (as measured via scRNA-seq) at increasing distances along each epithelial segment. (Right) Predictions (y-axis) of the DV score for each MERFISH-identified OSN subtype based upon the DV scores (x-axis) of the nearest neighbor OSNs from other OSN subtypes, using a spatial k-nearest neighbors regression approach with the DV scores associated with the detected OR as measured by scRNA-seq. E. Schematic depicting how individual epithelial segments were aligned to a common coordinate system using the positions of the identified ORs in a manner that was agnostic to the DV score of their respective OSN subtypes (see Methods). F. (Left and middle) Comparison of the DV scores (as identified via scRNA-seq using the expression of non-OR genes for each OSN subtype), to the OR spatial indices described in past reports that measured the expression of OR genes via bulk RNA-sequencing in microdissected regions of the MOE^20,21^. (Right) Results of support vector regression models trained to predict the DV score associated with each OR using the levels of OR expression measured recently via bulk RNA-sequencing in microdissected samples^32^. G. Comparisons of predicted DV scores for each OSN subtype across separate MERFISH coronal and sagittal MOE datasets, with DV scores for each OSN predicted using the DV scores of nearby neighboring cells (see Methods). H. Analysis of OR distributions in a separate MERFISH dataset from Bintu et al. (see Methods). (Left) the DV scores associated with the ORs detected in an example section. (Right) DV scores associated with each OR in epithelial segment from all replicates, before and after the alignment procedure. I. Silhouette score from hierarchical clustering of the pairwise distance matrix (in common coordinate space) of the spatial positions of OSN subtypes, with the indicated number of clusters, with the mean and 95% confidence interval across restarts indicated. J. (Left) The discriminability of the spatial positions of indicated types of OR pairs, as measured via area under the receiver-operator curve (AUROC) analysis. (Right) Comparison of the AUROC of the DV scores for each OR pair with the AUROC of their spatial positions. AUROCs were binned by their values for the DV score AUROC, and error bars depict the mean and 95% confidence interval of the mean for each bin. The correlation was calculated for all pairs. Histograms depict the number of pairs with each AUROC value; note most AUROC values are close to 1.

**Figure S3.**
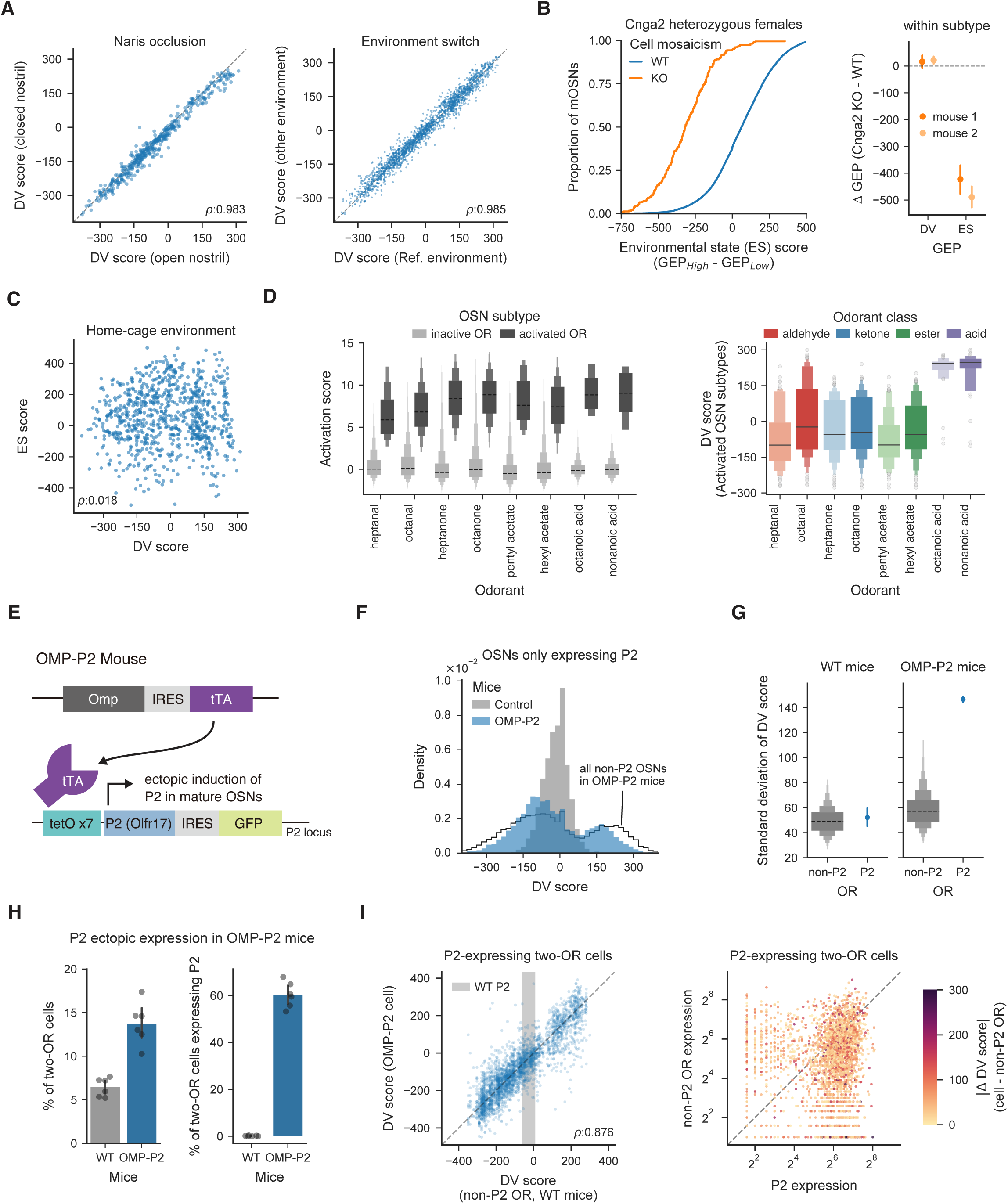
DV scores reflect cell identity not OR expression, related to Figure 3. A. (Left) DV scores for each OSN subtype for OSNs from open or occluded nostrils after 4 weeks of chronic naris occlusions. (Right) DV scores for each OSN subtype from mice housed in separate olfactory environments, whose odorants differentially activate the OR repertoire. B. (Left) Cumulative distribution of environmental state (ES) scores, which reflect ongoing OR-dependent activity and are computed as the difference between the GEP_High_ and GEP_Low_ activity GEPs^54^, for OSNs derived from Cnga2 heterozygous females expressing either the WT or loss of function allele of Cnga2. (Right) within OSN subtype change (KO – WT) in GEP usage for the DV and ES GEPs; note that ES scores depend upon CNGA2, while DV scores do not. C. DV and ES scores for each OSN subtype, for data from mice housed in a standard home-cage environment; the lack of relationship indicates that DV scores are likely activity-independent. D. Act-seq-based measurements of acute odor-driven activity (quantified as the Activation Score, which summarizes acute odor-driven increases in ∼200 immediate early and other genes^54^), for mice exposed to the indicated odorants, depicting Activation Scores for responsive and non-responsive OSN subtypes (left), and the distribution of DV scores for activated OSN subtypes (right); note that only the acids exhibit significant spatial clustering of their responses, likely because acids are largely detected by Class I ORs in the dorsal-most zone. E. Schematic depicting induction of the ectopic expression of the P2 OR in mature OSNs in the OMP-P2 mouse. F. Histogram depicting the DV scores for OSNs expressing P2, from either the entire integrated dataset (grey) or from OMP-P2 mice (blue), as well as the distributions of DV scores across all non-P2-expressing OSNs in OMP-P2 mice (black line). G. The distribution of standard deviations of the DV score for OSNs expressing P2 and non-P2 ORs from wild-type (WT) mice or OMP-P2 mice. Error bars indicate bootstrapped 95% confidence intervals. H. (Left) Quantification of the percent of OSNs expressing two ORs in WT and OMP-P2. (Right) The percent of these two-OR cells containing P2 (and another non-P2 OR). Dots indicate individual replicates and error bars represent the mean and bootstrapped 95% confidence interval of the mean. I. (Left) The DV score for two-OR cells in OMP-P2 mice expressing P2 and a non-P2 OR, plotted as a function of the DV score associated with the non-P2 OR in control mice. Shaded gray area depicts the interquartile range of the DV score for OSNs in control mice that singularly express P2. (Right) The expression of P2 and the non-P2 OR in OSNs expressing two ORs, colored by the difference in DV scores between each cell and the respective OSN subtype expressing the non-P2 OR in control animals.

**Figure S4.**
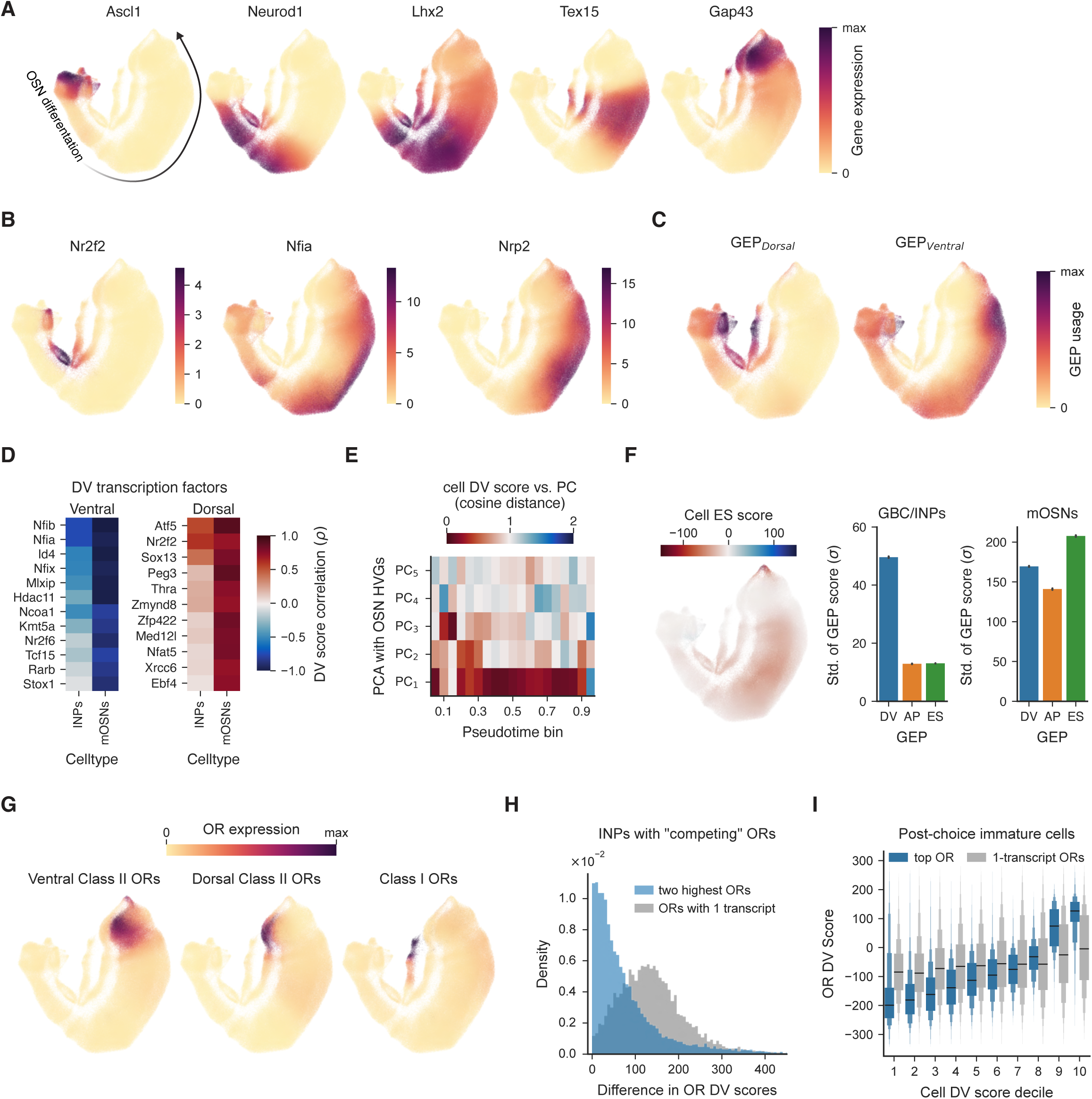
Variation in gene expression during OSN differentiation, related to Figure 3. A. UMAP plot depicting the indicated marker genes across the differentiating OSN dataset (individual dots are single cells, shading is normalized gene expression); note, the majority of iOSNs and all mOSNs are excluded in this dataset. For visualization purposes, gene expression (the number of unique molecular identifiers (UMIs) in the sequencing data for each indicated gene), was smoothed based on the nearest neighbor graph (see Methods). B. Similar to A), but for the expression of example DV genes in differentiating OSNs. C. GEP usages for GEP_Dorsal_ and GEP_Ventral_ for each differentiating cell, calculated using the GEP weights for each gene as identified in mature OSNs. D. The correlation of the cellular expression of each of the indicated transcription factors (all of which participate in the DV score and are therefore DV genes) with the DV scores of each cell (with data derived from either the mature OSN or differentiating OSN datasets). E. The similarity (cosine distance) of the cell DV score and the top principal components of gene expression, where PCA was performed using OSN variable genes for cells in each pseudotime bin. Note that OR expression does not begin until psuedotime > 0.4. F. (Left) UMAP depiction of the usage of ES score GEPs in differentiating OSNs. (Right) The standard deviation in GEP scores for DV, environmental state (ES), and anteroposterior (AP) scores. ES scores reflect ongoing OR-dependent activity and are computed as the difference between the GEP_High_ and GEP_Low_ activity GEPs; AP scores were computed as the difference between GEP_Anterior_ and GEP_Posterior_, which include a handful of genes implicated in OSN targeting along the anteroposterior axis of the olfactory bulb and which reflects OR-dependent but odor-independent activity (as recently described^54^, and see Figure 7). G. UMAP plots depicting the maximum levels of expression for the indicated types of ORs. Note the separate developmental trajectories adopted for differentiating OSNs that eventually choose each type/class of OR. H. For cells expressing multiple “competing” ORs expressed at moderate levels (three or more UMIs), the difference in DV scores of expressed ORs (as determined in mature OSNs) compared to similar differences amongst the ORs expressed in the same cells at the lowest expression level (i.e., one UMI per cell). I. For post-choice cells that express only a single OR (defined here as having three or more unique molecular identifiers (UMIs)), the DV score associated with the chosen OR (determined in mature OSNs) compared to the mean DV score associated with all those ORs expressed at low levels (only one UMI per cell), plotted as a function of each cell’s DV score (split into deciles).

**Figure S5.**
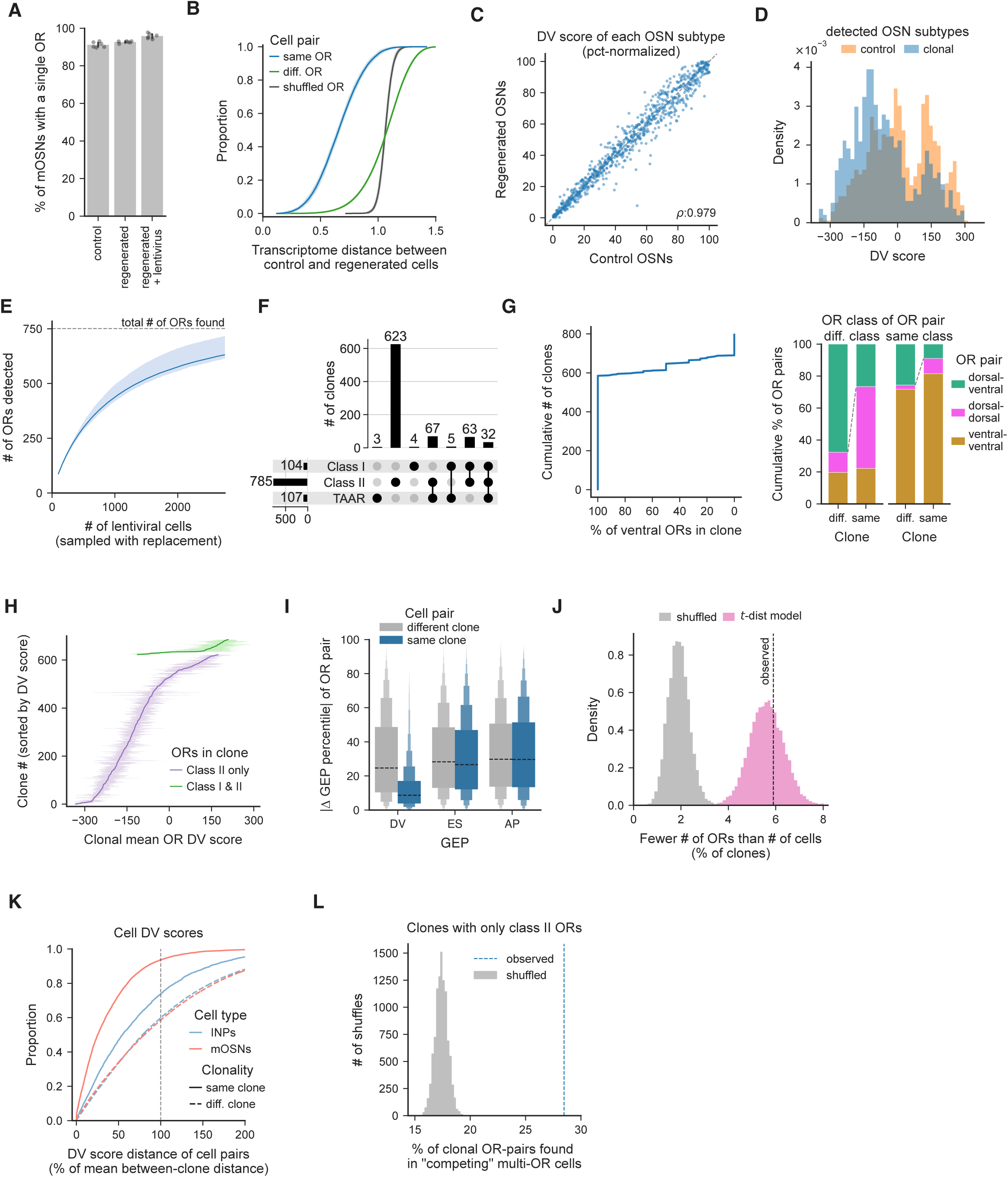
OSN clones have restricted cell fates, related to Figure 3. A. The percent of mature OSNs expressing a singular OR in either WT control mice or in regenerated cells post-methimazole from the lentiviral lineage tracing experiment. Dots indicate individual replicates and error bars indicate bootstrapped 95% confidence interval of the mean. B. The transcriptome distance, defined as the cosine distance of GEP usages, between control and regenerated cells that express either the same or different ORs, or when OR labels are shuffled across OSNs. Shaded error indicates the mean and bootstrapped 95% confidence interval of the mean. C. The percent-normalized DV scores associated with each OSN subtype in control and regenerated conditions. D. The distribution of DV scores for all OSN subtypes in control conditions and the distributions of DV scores of ORs detected in lentiviral-infected cells in the clonal lineage tracing experiment. E. Cumulative number of ORs detected as a function of the number of barcoded cells. Line and shaded error indicates the mean and bootstrapped 95% confidence interval of the mean. F. UpSet plot depicting the number of clones that contained OSNs collectively expressing each combination of OR classes. The total number of cells expressing each receptor type is indicated on the bottom Y axis. The bar graph indicates the number of clones harboring OSNs expressing receptors of the dotted classes (below each bar). Note that individual clones can harbor mature OSNs expressing ORs from different classes. G. (Left) The cumulative number of clones, as a function of the percent of ORs in the clone that were ventral. Note, most clones exclusively contain ventral ORs. (Right) For pairs of ORs from either the same or different receptor classes (out of class I ORs, class II ORs, or TAARs) and for pairs of cells that were or were not clonally-related, the cumulative percent of pairs in which the two ORs were both dorsal or ventral (or one was dorsal and the other was ventral). Note, clonally-related OR pairs were more likely to both be either dorsal or ventral. H. For each clone, the mean and bootstrapped 95% confidence intervals of the DV scores (as measured in control conditions) associated with the OSN subtypes of that clone. Clones are sorted by the mean DV score associated with their expressed ORs. I. The distances in DV, AP, and ES scores (as measured in control conditions) of OSN subtypes found in pairs of cells, colored by whether each pair was clonally-related or not. ES scores reflect ongoing OR-dependent activity and are computed as the difference between the GEP_High_ and GEP_Low_ activity GEPs; AP scores were computed as the difference between GEP_Anterior_ and GEP_Posterior_, which include a handful of genes implicated in OSN targeting along the anteroposterior axis of the olfactory bulb and which reflects OR-dependent but odor-independent activity (as recently described^54^, and see Figure 7). J. The percent of clones with fewer ORs than cells (thus contained multiple cells expressing the same OR) for the observed data, for data with shuffled clonal labels, and for clones generated using the observed *t-*statistics to sample ORs (see Methods). K. The cumulative distribution of the distance in the DV scores of pairs of INPs or mature OSNs that either came from same or different clones. L. The percent of clonally-related pairs of mature OSNs expressing class II ORs in which the different ORs were found to be co-expressed within single differentiating OSNs at the “competing” phase (two ORs, each expressed with three or more UMIs), compared to the percentage found when clonal labels were permutated across OSNs.

**Figure S6.**
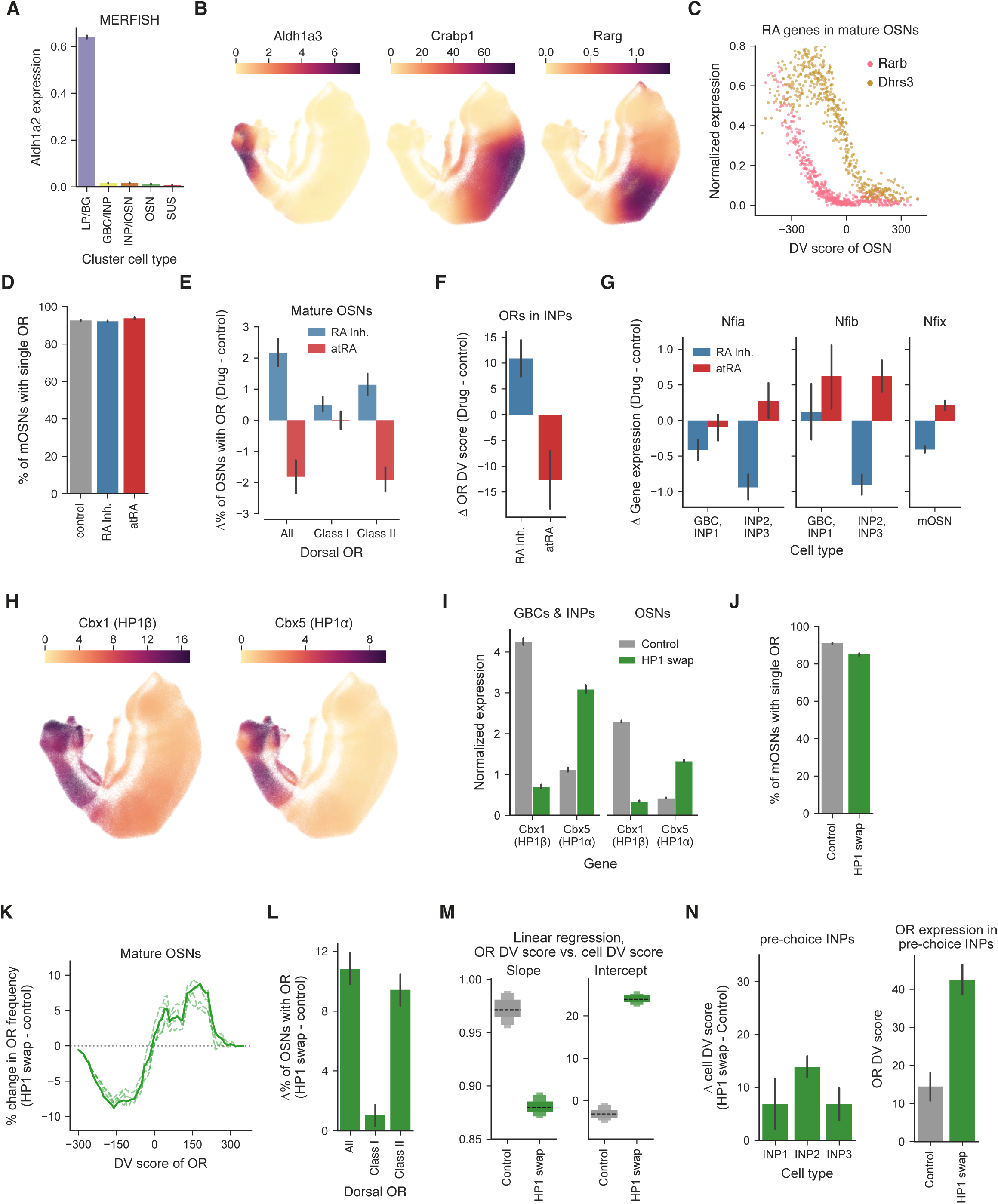
RA signaling components vary spatially to influence OSN identities, and heterochromatin binding influences OR choice, related to Figures 4 and 5. A. Expression of *Aldh1a2*, as measured via MERFISH, in each cell type (log-normalized number of transcripts per cell). LP/BG, lamina propria and Bowman’s gland cells. B. Expression of the genes involved in RA signaling in the differentiating OSN dataset, including the retinaldehyde dehydrogenase 2 (*Aldh1a3*), the cellular retinoic acid binding protein *Crabp1* in immature cells, and the nuclear retinoic acid receptor gamma (*Rarg*). Heatmaps indicate the number of unique molecular identifiers (UMIs) detected for the indicated gene, smoothed by the nearest neighbor graph (see Methods). C. Normalized gene expression of the RA-related genes *Rarb* and *Dhrs3* in mature OSNs, as a function of the DV score of each OSN. Cells were binned into equal-frequency bins and each dot depicts the average expression in each bin. D. The percent of mature OSNs post-methimazole-induced regeneration that expressed single ORs, for animals treated with either control vehicle, the RA inhibitor (RA Inh.) WIN 18,446, or atRA, with error bars indicating 95% confidence intervals of the mean. E. For mice given each drug, the change in frequency (relative to control mice) of the percent of cells expressing dorsal ORs, plotted for all dorsal ORs and separated by OR class. Error bars depict bootstrapped 95% confidence intervals. F. Change in the DV scores associated with each OR in mature OSNs for the ORs expressed in pre-choice INP cells. For cells expressing multiple ORs the DV scores associated with each OR were weighted by their expression levels. G. The expression of the NFI family transcription factors *Nfia, Nfib*, and *Nfix* in the indicated cell types relative to their expression in control mice, for mice given either the RA inhibitor or atRA. H. Similar to (B) but for the expression of HP1β and HP1α in differentiating OSNs. I. The Normalized expression of HP1β and HP1α in either the control or HP1 swap mice, in either differentiating GBCs and INPs or in mature OSNs. J. Similar to (D) but for the percent of mature OSNs from control or HP1 swap mice that express a single OR. K. Change in frequency of ORs with given DV scores for data from HP1 swap relative to control mice. Dashed lines indicate individual replicates. L. Similar to (E) but for the change in the percentage of cells expressing dorsal ORs in HP1 swap mice relative to control mice. M. Bootstrap estimates of the slope and intercept for robust linear regression models fit on the cell to OR DV score mapping in each condition. The observed change in slope in the HP1 swap mice indicates that the relationship between cell and OR-associated DV scores has changed. N. For pre-choice INPs, (left) the mean change in the cell DV scores (HP1 swap – control) for the cells at the indicated stage or (right) the mean DV scores associated with the ORs (as measured in mature OSNs) for the ORs expressed in INPs from each condition, where for cells expressing multiple ORs the DV scores associated with each OR were weighted by their expression. Error bars depict 95% confidence intervals of the mean.

**Figure S7.**
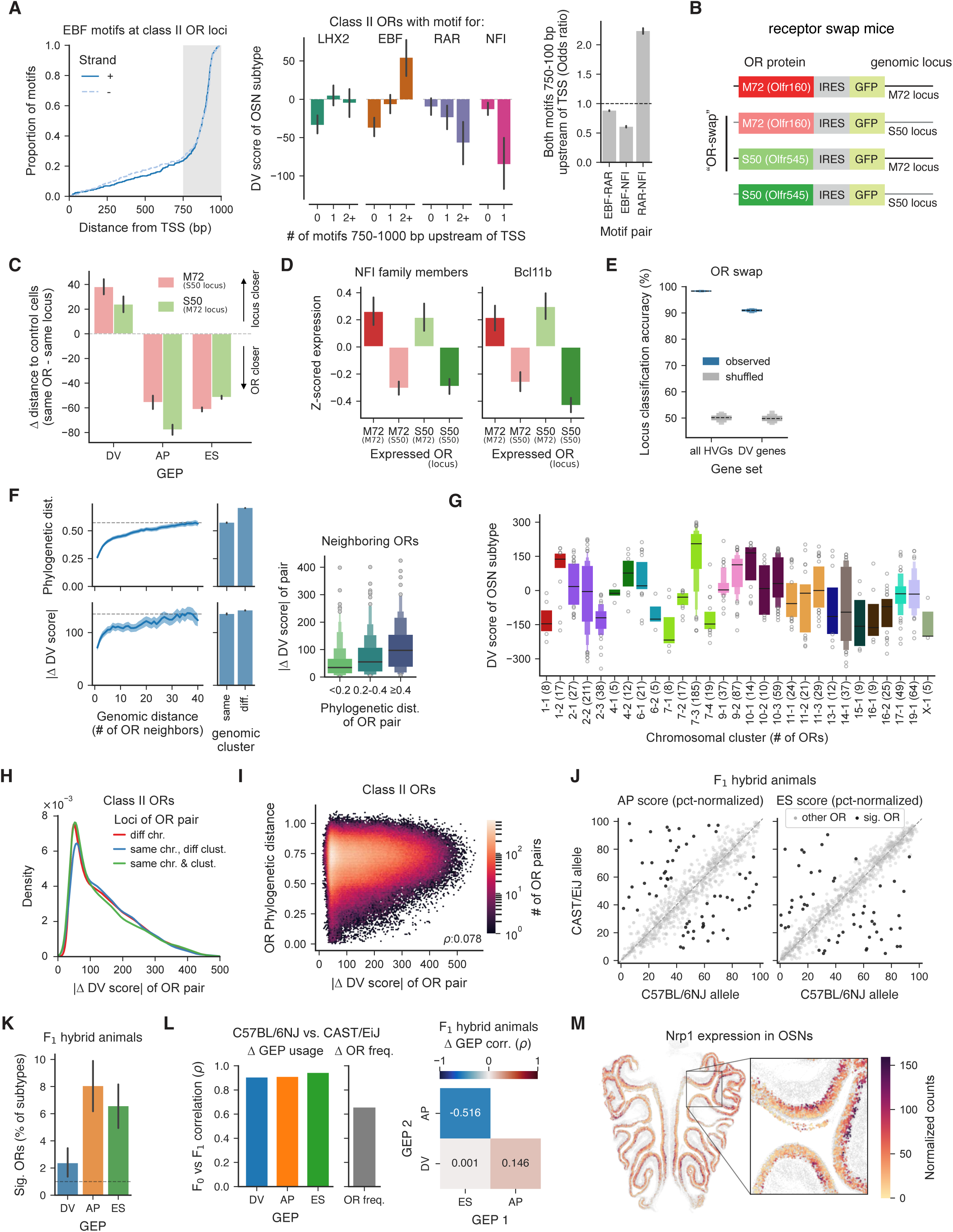
DV scores are a genomic property of each OR, and AP scores predict apical-basal epithelial position, related to Figures 6 and 7. A. (Left) Cumulative proportion of EBF motifs in the 1kb region upstream of each OR’s transcriptional start site (TSS). Note the density of motifs in the 750-1,000 bp region upstream of each TSS. (Middle) The mean and 95% confidence interval of the DV score for OSN subtypes whose ORs have the indicated number of motifs for each transcription factor in the upstream 750–1,000 bp region. (Right) The odds ratio of having the given motif pair in the same OR promoter region. B. Schematic of the receptor swap experiment, in which OSNs expressed the M72 or S50 OR from either the M72 or S50 genomic locus. Note that although these mice are referred to as receptor “swaps” in mice in which M72 (or S50) is expressed from the S50 (or M72) locus, there remains M72 (or S50) expressed from its original locus in the genome. As a consequence, in these mice there are two separate genomic locations associated with the same receptor. C. The distances in GEP usages for each cell from the OR swap mice to the mean of the respective control animal (in which ORs were expressed from their endogenous loci). The difference in distances to the same OR vs. same locus control animal is shown for each GEP. ES scores reflect ongoing OR-dependent activity and are computed as the difference between the GEP_High_ and GEP_Low_ activity GEPs; AP scores were computed as the difference between GEP_Anterior_ and GEP_Posterior_, which include a handful of genes implicated in OSN targeting along the anteroposterior axis of the olfactory bulb and which reflects OR-dependent but odor-independent activity (as recently described^54^, and see Figure 7). D. The average expression of the NFI family and Bcl11b transcription factors, both of which are enriched in ventral OSNs, across the animals examined in the OR swap experiments. E. Accuracy of support vector machine classifiers at distinguishing the genomic locus of each cell, using either all HVGs or the DV genes, for the observed data and data in which the genomic locus was shuffled across cells. F. (Left) For OSN subtypes whose ORs are located in the same genomic cluster, the mean phylogenetic distance (top, cophenetic distance) or distance in DV scores (bottom), plotted as function of genomic distance, as measured via the number of functional OR genes separating the ORs of each OSN subtype. (Right) The distribution of DV score distances for pairs of OSN subtypes whose ORs are the nearest genomic neighbors, separated based on the phylogenetic distance of the OR pair. G. The distribution of DV scores associated with each OSN subtype, for OSN subtypes whose ORs are located in the indicated chromosomal clusters. Note, class I ORs are found in cluster 7-3 and TAARs are located in cluster 10-1. H. Kernel density estimate of the distance in DV scores for pairs of OSN subtypes whose ORs are located either within the same or different chromosomal clusters. I. Phylogenetic distances (i.e., cophenetic distance) of OR pairs as a function of their associated DV scores. J. The AP and ES score for each OSN subtype, evaluated separately for cells from F_1_ animals that chose either the C57BL/6J or CAST/EiJ allele for a given OR. OSN subtypes with significant changes in GEP usages across strains are highlighted in black. Note that this strain-specific variation is expected to be the consequence of coding changes in the receptor itself (see Figure 6D). K. The percent of OSN subtypes with strain-specific changes for each GEP. L. (Left) For OSN subtypes with significant changes for each GEP, the correlation between the change in GEP usages (CAST – C57 allele) in F_1_ hybrid animals and the change in GEP usages across F_0_ animals (CAST – C57 mice), and the correlation in the change in OR frequency across alleles in F_1_ animals with that across strains in F_0_ animals. (Right) For each GEP pair, the correlation in the change in GEP usage across strains, for the set of OSN subtypes with significant changes in either GEP from each GEP pair. M. The expression of the GEP_Posterior_ axon guidance gene *Nrp1*, as measured directly via MERFISH. Gene expression was normalized based on the total number of transcripts detected in each OSN.

## Supplementary Materials

### Supplementary Tables

**Table S1. Dorso-ventral (DV) related genes and associated gene categories, related to** Figure 1.

**Table S2. Probe sets for in situ hybridization chain reaction (HCR) experiments, related to** Figure 6.

### Resource Availability

#### Lead Contact

Further information and requests for resources and reagents should be directed to and will be fulfilled by the Lead Contact, Sandeep R. Datta.

#### Materials availability

This study did not generate new unique reagents.

#### Data and code availability

Raw and processed next generation sequence data have been deposited at the NCBI Sequence Read Archive (SRA, accession SRP318630) and Gene Expression Omnibus (GEO, accession GSE173947) databases. The code for calculating GEP usages of OSNs can be found at https://github.com/dattalab/Tsukahara_Brann_OSN. Code for analyzing DV scores can be found at https://github.com/dattalab/Brann_olfactory_dorsoventral.

### Experimental model and study participant details

#### Animals

C57BL/6J mice were obtained from Jackson Laboratory (Stock No. 000664). Mice expressing OMP-IRES-tTA and tetO-P2-IRES-GFP (OMP-P2 mice) were obtained from the Lomvardas lab and maintained in the Datta Lab^1,2^. Cnga2 knockouts animals (Jackson Laboratory Stock No. 002905) were obtained from the Dulac lab and maintained by breeding heterozygous females with C57BL/6J males^3^. HP-1 swap mice (mice with loxP-HP1β-STOP-loxP-HP1α alleles at the HP1β locus crossed to Foxg1-Cre [Jackson Laboratory Stock No. 029690]^4^ such that knockout of HP1β is rescued via the expression of HP1α) were generated by Martín Escamilla-Del-Arenal using standard homologous recombination, and will be described in a separate forthcoming manuscript. Wild-derived CAST/EiJ mice were obtained from Jackson Laboratory (Stock No. 000928), and CAST/C57 F_1_ hybrids were generated in lab by crossing CAST males with C57 females. Mice of either sex between 6–16 weeks-old were used for experiments. Mouse husbandry and experiments were performed following institutional and federal guidelines and were approved by Harvard Medical School’s Institutional Animal Care and Use Committee.

### Method details

#### Naris occlusion, environment swap, and Act-seq activity manipulations

Data from wild-type mice housed in home-cage environments, or from mice that either underwent transient naris occlusion for a week or were housed in novel olfactory environments, was all previously generated^5^. Act-seq was performed as previously described, by exposing mice to monomolecular odorants on filter paper for two hours prior to OSN dissociation and scRNA-seq^5^. The odorants that were analyzed in further detail were heptanal, octanal, heptanone, octanone, pentyl acetate, hexyl acetate, octanoic acid, and nonanoic acid, which were all diluted in dipropylene glycol (DPG) based on their vapor pressures to give a nominal concentration of 500 ppm.

#### Methimazole-induced regeneration, intranasal lentiviral injections, and retinoic acid experiments

The main olfactory epithelium (MOE) was ablated and induced to regenerate via a single intraperitoneal injection of methimazole (50 mg/kg) in adult mice (>6 weeks old), as described elsewhere^6^. Intranasal lentiviral injections were performed 36-48 hours after methimazole injection, at timepoints in which the OSNs have sloughed off and the epithelium is a single monolayer of activated horizontal basal stem cells (HBCs). Mice were anaesthetized with Ketamine/Xylazine (100/10 mg/kg), and intranasal lentiviral injections were performed via a Hamilton syringe connected to thin PE-10 tubing; 15-25 µL of virus (at a titer of 1–10 x 10^8^ TU/mL) was injected over the course of 2-3 minutes. scRNA-seq was performed 4–6 weeks after methimazole injection, at timepoints under which much of the epithelium has regenerated and the infected progenitors have differentiated to give rise to clones of newly-born cells, many of which are OSNs.

Manipulations of retinoic acid signaling were performed in adult mice that received a single dose of methimazole (50 mg/kg). Starting 72 hours after methimazole injection, mice received either the ALDH1A2 inhibitor WIN 18,446 (5 mg/kg) or all trans-Retinoic Acid (atRA, 5 mg/kg). WIN 18,446 was dissolved in DMSO and atRA was dissolved in 5 % DMSO (dissolved in corn oil), and drugs were delivered subcutaneously. Mice within each cohort received either drug or vehicle daily for 3.5 weeks after methimazole-induced regeneration, and scRNA-seq was performed after 4 weeks, and animals receiving each drug or vehicle control were dissociated and sequenced together. The effects of the RA inhibitor WIN 18,446 were verified via observation of the testes^7^.

#### Single-cell RNA-sequencing and data processing

Isolation of single cells from the MOE and fluorescence-activated cell sorting (FACS) was performed as described^5^. In brief, mice were euthanized via carbon dioxide inhalation, and the epithelial tissue of the MOE was dissected. Individual mice were used for each replicate. Cells were dissociated with Papain and DNase-I (Worthington), incubated at 37°C for 60 minutes, triturated and then washed and resuspended in Hibernate-A medium supplemented with fetal bovine serum; these solutions also contained transcriptional and translation inhibitors (5 mg/mL of Actinomycin D, 10 mg/mL of Anisomycin and 10 mM of Triptolide, all obtained from Sigma). FACS was performed to enrich for OSNs expressing fluorescent reporters in the OMP-P2 and lentiviral experiments. In all other experiments, FACS was performed to enrich for live singlets.

scRNA-seq was performed using the 10x genomics platform using the default protocols (CG000315 Rev E) for the Chromium Next GEM Single Cell 3ʹ Kit v3.1. For each replicate, cells were loaded at a concentration predicted to yield 10,000 cells, or to obtain the maximum number of cells given the measured concentrations when the total number of was less than 10,000. For the OMP-P2 experiments, GFP-positive and GFP-negative cells were mixed and loaded together in a single lane of the Next GEM chip. In the lentiviral experiments, Venus-positive cells and negative cells from each mouse were not mixed and were loaded into separate lanes. For experiments with genetic or experimental controls (like the CNGA2, RA, and HP1 swap experiments), control and experiment animals were processed together, loaded onto the same chips, and were sequenced together.

The resulting libraries were quantified via qPCR and were sequenced with paired-end sequencing on the Illumina NovaSeq or NextSeq platforms (with Read1 = 28 cycle, Index (i7) = 10 cycle, Index (i5) = 10 cycle, Read2 = remaining number of cycles). Demultiplexed fastq files were processed in a manner similar to that previously described^5^, using a custom Nextflow pipeline that ran the Cell Ranger (version 6.1.2) count pipeline, as well as additional post-processing steps to identify and remove multi-mapped reads and regenerate the resulting cell by gene matrix. The Ensembl GRCm39 mouse genomic index (version 105) was used with a custom GTF file containing extended 3’ UTR annotations for some ORs and TAARs, to facilitate their identification in the 3’-biased scRNA-seq data. Cell doublets and low-quality cells were excluded, and OSNs were identified via iterative subclustering of the latent spaces of scVI models, as previously described^5,8^, based on their expression of known marker genes and ORs.

#### Clonal lineage tracing

Clonal lineage tracing was performed using a lentiviral library with ∼10M DNA barcodes at the 3’ UTR side of transcripts for Venus fluorescent protein and upstream of the polyA sequence (Cellecta CloneTracker XP). To amplify the lentiviral barcodes from the RNA transcripts of in each cell, a primer (5’-<u>GTGACTGGAGTTCAGACGTGTGCTCTTCCGATCT</u>CCGACCACCGAACGCAACGCACGCA-3’) whose 5’ end overlapped with the Truseq Read 2 (underlined) and whose 3’ end was complementary to a constant region upstream of the barcode was spiked in (final concentration 0.5 µM) during the initial RT and cDNA amplification steps; this approach increased the yield of the lentiviral barcodes compared to targeted amplification from the already-amplified cDNA libraries. Amplified barcode fragments were purified from the cDNA library and barcode sequencing libraries were constructed by performing PCR on these fragments with the Library Index PCR primers (which bind to Truseq); the resulting barcode libraries were mixed and sequenced together with the final 10x libraries and were subsequently demultiplexed computationally.

#### Spatial profiling via MERFISH

MERFISH was performed using the Vizgen MERSCOPE platform. A custom 300-plex codebook, which contained nearly 200 ORs and 100 non-OR genes consisting of cell type marker genes, DV-related genes, and other highly-variable OSN genes, was designed. Bulk sequencing and scRNA-seq was used to identify suitable ORs and genes, and the total abundance of the gene panel was ∼7600 FPKM to avoid optical crowding. ORs were chosen on the basis of their DV scores and frequency of expression to obtain a set of ORs predicted to uniformly span the MOE at relatively-higher frequencies. Due to their high abundance, additional probes for OSNs (*Omp, Calm2*), and sustentacular cells (*Cbr2, Cyp2g1*) were imaged separately and sequentially (along with the staining for DAPI, polyT, and cell boundaries).

Fresh frozen 10 µm coronal samples from the MOE from young (4-5 week) mice were processed according to the manufacturer’s user guides (Vizgen 91600002 Rev E and 91600112 Rev C). In brief, tissue section, fixation, permeabilization, and autofluorescence quenching were performed used the guidelines for fresh frozen tissue. However, the RNA anchoring steps from the formalin-fixed, paraffin-embedded (FFPE) tissue sample preparation guide were added to help the thin MOE tissue adhere to the MERSCOPE slides. Cell boundary staining, gel embedding, clearing, and probe hybridization steps were performed without modification from the fresh-frozen sample preparation guide. Samples were imaged on the MERSCOPE instrument using the 300-plex imaging kit, and transcripts were decoded using the MERLIN pipeline provided by Vizgen.

#### CUT&RUN

CUT&RUN was performed to measure the levels of H3K9me3 in C57/CAST F_1_ animals, using the reagents and instructions from a commercial kit (EpiCypher CUTANA CUT&RUN Kit, Version 4). In brief, the MOE was dissociated as in the scRNA-seq experiments, and 500k unsorted cells were used for each reaction, and two technical replicates were used for the cells from each animal. Cells were incubated overnight with ConA beads and 0.5 µg of a Rabbit polyclonal H3K9me3 antibody (Abcam ab8898), and on the following day permeabilized cells were incubated with pAG-MNase to target the digestion of DNA, which was purified with SPRIselect beads. E. coli spike-in DNA was used to normalize the yields across samples.

Sequencing libraries were prepared from 5 ng CUT&RUN DNA, or the entire yield for replicates where the yield was <5 ng. Some experiments also included a H3K4me3 positive control (EpiCypher 13-0041), which yielded the expected enrichment for transcriptional start sites, IgG negative control (EpiCypher 13-0042) antibodies, which showed minimal enrichment, but these were omitted for subsequent experiments given the low background in the H3K9me3 CUT&RUN data. Sequencing libraries were generated using the NEBNext DNA library kit, following the instructions and formulations in the EpiCypher CUT&RUN library prep kit (14-1002). CUT&RUN libraries were pooled and sequenced on the Element AVITI machines with the Cloudbreak Freestyle sequencing kit (2×75 cycles).

#### In situ hybridizations

In situ hybridization for genes in the MOE was performed via the Hybridization Chain Reaction (HCR) method^9^, using 16 µm cryosections from either fresh or fixed frozen MOE tissue, from 4 to 6-week-old animals. Fixed tissue was decalcified in 0.45M EDTA and equilibrated in 30% sucrose prior to freezing. In situs were performed according to manufacturer instructions for HCR v3.0 (Molecular Instructions, Rev. 4), except the proteinase K solution was changed to 1 µg/mL or omitted entirely. Split DNA oligo probes for ORs (∼15–20 probes per gene) were manually designed using regions in each OR’s cDNA sequence that were homologous between C57BL/6J and CAST/EiJ animals but lacked homology to other ORs. Probes for the dorsal marker *Nqo1* and reference ORs whose positions remaining constant across strains were also designed in a similar manner and used as reference landmarks to identify the positions of ORs of interest. The sequences of oligo probes can be found in Supplemental Table 2.

Sections were counterstained with DAPI and confocal imaging was performed using a Nikon Ti inverted spinning disk microscope equipped with a Yokogawa CSU-W1 Spinning disk (50 μm pinhole), a Plan Fluor 40x/1.3 NA oil objective, and 405, 561, and 640 nm laser lines. Tiled images of the MOE were captured using an Andor Zyla 4.2 Plus sCMOS monochrome camera and Nikon Elements Acquisition Software (AR 5.02). OSNs expressing each OR were manually counted, and the fraction of OSNs within the dorsal zone was assessed by comparing to positions of each OSN relative to Nqo1^+^ OSNs. Note, although imaging and quantification was performed across entire sections, schematics depict the results from F_0_ data due to space limitations and due to the difficulty in visualizing the expression of single ORs across the entire MOE.

### Quantification and statistical analysis

#### Data integration of scRNA-seq replicates to extract mature OSNs

Nearly 5 million cells from over 360 replicates were uniformly processed using a custom Nextflow pipeline to run Cell Ranger and custom postprocessing steps for each replicate. These replicates included the 150 replicates (and 780 thousand mature OSNs) described in a previously published dataset^5^, as well as over 200 additional replicates (containing another 1.4 million OSNs). The additional data consisted of the mice used as part of the experimental conditions described herein (e.g. the lentiviral experiments), as well as additional unpublished datasets that were generated for other work, including from mice that had been exposed to monomolecular odorants, as well as from smaller numbers of mice housed in different environments conditions, and control and knockouts for various genetic manipulations. Mice of both sexes were used for experiments. Note, although some of these lines had not been backcrossed to C57BL/6J backgrounds, few differences in DV scores or gene expression were observed across replicates, even though the entire dataset consisted of 360 replicates from many separate experiments.

A combination of unsupervised and semi-supervised methods was used to map the cells from all replicates into a common latent space to perform cell type identification. The gene set used for this embedding were the top 3000 genes (excluding ORs, mitochondrial and ribosomal genes) identified using the Scanpy highly variable gene function, with the “seurat_v3” flavor. First, unsupervised learning was used to fit a scVI model (n_hidden: 128, n_latent: 30, n_layers: 2, gene_likelihood: “nb”, dropout_rate: 0.2, with a batch key to identify each replicate) on the raw counts of these genes in a smaller reference dataset of ∼250 thousand cells from wild-type animals housed in standard home-cage conditions. The resulting scVI latent embeddings from this dataset were clustered via the Leiden algorithm and cluster labels were manually annotated based on known cell type markers. The trained scVI model and the manually-annotated cluster labels from this subset were then used to train a semi-supervised model using scANVI. The trained scANVI model was then applied to query the entire dataset, resulting in a common embedding of all cells into a 30-dimensional latent space. Cell types in this common latent space were identified by transferring the labels from the manually-annotated reference subset, using previous weighted nearest neighbor approaches for label transfer of scANVI models^10,11^. For cells from each replicate, their resulting neighbors (in the 30-dimensional latent scANVI space) from the annotated subset were found using PyNNDescent, distances were converted into affinities using a gaussian kernel, and cell type probabilities and matching uncertainty were calculated for each cell.

Non-neuronal cells, as well as low-quality doublets and dying cells were excluded, and mature were extracted from the cell type labels of the integrated dataset. Due to the dissociation and FACS approaches used, roughly 50% of cells from most replicates were OSNs. OSNs were filtered to retain only cells expressing a single OR, and the resulting dataset of 2.3 million OSNs was used for downstream analyses. Except where noted, only cells expressing single ORs were used for analyses, and OR expression was defined with a threshold of at least 3 transcripts (unique molecular identifiers (UMIs)) that mapped to that OR.

#### Identification of zonally-restricted genes

Zonal indices for each OR were obtained from prior work^12,13^, and were discretized into five zones, with zone 1 being the most-dorsal zone and zone 5 the most-ventral. The mean for each subtype for each variable gene was calculated and genes whose expression was on average a log-fold higher among the subtypes in its maximum versus second-highest zone were considered as zonally-restricted genes, and the number of genes for each zone were computed. Note, no genes were enriched in intermediate zone ORs (e.g. zone 2–4), consistent with gradients of gene expression that monotonically increase or decrease with the dorsoventral score.

#### Identification of the dorsoventral (DV) score

Consensus non-negative Matrix Factorization (cNMF) was used to decompose the transcriptomes of each OSN into a smaller set of interpretable factors^5,14^. The same NMF factors (referred to as gene expression programs (GEPs)) that were previously identified were used, and the gene loadings for these GEPs were applied to all datasets^5^. Importantly, cNMF was performed on a set of ∼1,300 highly variable genes (HVGs) that did not include any OR genes and thus the resulting gene loadings and GEP usages do not depend on the expressed ORs. These GEPs included two associated with neuronal activity, (GEP_High_ and GEP_Low_), two containing dorsal and ventral genes (GEP_Dorsal_ and GEP_Ventral_), and two relating to anteroposterior positions (GEP_Anterior_ and GEP_Posterior_). These GEPs were used in an orthogonal manner and their differences in usage was summarized into single metrics: the environmental state (ES) score = GEP_High_ - GEP_Low_, the dorsoventral DV score = GEP_Dorsal_ - GEP_Ventral_, and the anteroposterior (AP) score = GEP_Anterior_ - GEP_Posterior_) to capture the continuous variation across these 3 independent axes (ES, DV, and AP). The 248 genes (out of the set of highly variable genes) whose expression across OSNs was correlated with the DV scores (R^2^ > 0.5 for linear regression to predict the DV score from each gene and spearman’s rho > 0.5 at the OSN subtype level) were consider as DV genes and were used for downstream analyzes. DV genes were categorized into 10 different functional groups (transcription factors, axon guidance and cell adhesion, cellular metabolism, cell signaling, cell growth, cytoskeletal, GTP-activity, RNA-related, ion channels and transporters, and synaptic genes) via a combination of GO pathway analysis, the presence of protein features like DNA binding domains, or via manual annotation. The full list of DV genes and their associated categories (for the 162 belonging to the above categories) can be found in Supplemental Table 1.

As described above, GEP usages and DV scores were calculated at the cell level using the gene loadings for each GEP. A small fraction (3%) of cells had DV scores that were zero, which can occur for cells that had little expression of either dorsal- or ventral-related genes and because NMF identifies usages for all GEPs at once. For analyses that required comparisons at the cell level, the DV scores of these cells were imputed using a multilayer perceptron (MLP) trained to predict the DV score of mature OSNs using the expression of the 248 DV genes in each cell. The MLP (with two hidden layers of size 50 and 10) was trained in PyTorch with the Adam optimizer (learning rate 0.01) and gave predictions that were highly correlated (rho > 0.99) at both the cell and OR level for training, testing, and validation data.

Mature OSNs expressing a given OR (i.e. those from the same OSN subtype) had similar DV scores. Therefore, the DV score associated with each OSNs expressing each OR (also referred to as an OR DV score) was calculated by taking the mean DV score across all the OSNs expressing a given OR in the entire integrated dataset. Except where noted, for all downstream analyses the DV score of the expressed OR in each cell was considered as the OR DV scores from the entire dataset and thus could be compared to each cell DV’s score, especially in experiments that decoupled a given cell’s DV score from that of its expressed OR. In some experiments, DV scores were percentile normalized at the cell level via sklearn’s QuantileTransformer. This transformer was fit on data from all cells to maintain the distribution of DV scores for cells from each subtype.

##### OR classes

Only ORs with functional protein activity (as assessed by their singular expression in mature OSNs). ORs with mean DV scores above 60 were considered as dorsal; this threshold was set such that ORs that had previously been classified as being part of the most-dorsal zone (zone 1) were considered as dorsal with respect to their DV scores. ORs were clustered based on their amino acid sequences and class I ORs were identified as the large cluster of ORs in the resulting phylogenetic tree that share homology with fish ORs and were located in a single genomic cluster on Chromosome 7. Phylogenetic distances were defined as the cophenetic distance of the resulting phylogenetic tree. Most rodent odorant receptors are class II ORs, which are located in multiple genomic clusters of genes that are scattered across the genome. ORs were considered part of the same genomic cluster if they were located on contiguous regions of the chromosome (separated less than 3 Mb from the start of previous OR gene). Neighboring ORs (i.e. those at neighboring genomic locations) were more likely to be phylogenetically similar. A small fraction of cells also express trace amine-associated receptors (TAARs). Except where specified, analyses used OSNs expressing either class I, class II, or TAARs though most results reflect properties of class II ORs, which make up the majority (∼87%) of all ORs. The set of 1007 OSN subtypes with at least 150 cells (median 1,500 OSNs per subtype) was used for most analyses.

##### Consistency of the DV score

Correlations of the DV scores across replicates were computed at the OSN subtype level, by comparing the mean for each subtype in a given replicate to that from all other data. To evaluate the consistency across data subsets, 150 cells were sampled for each OSN subtype, the mean for each OSN subtype was calculated separately for the first versus second half of cells for each subtype, the spearman’s correlation between data halves was evaluated, and this process was repeated 1,000 times. The spearman’s correlation reflects changes in the rank order of OSN subtypes and the absolute change in ranks was also evaluated on each restart, as well as for data in which OR labels were either permuted across cells whose ORs shared the same zonal index (discretized into 5 zones as above) or permuted across all cells. The mean for each OSN subtype was also evaluated separately for data from male and female animals.

The uniqueness of the DV score for each OSN subtype was determined using linear regression. Across 100 restarts, cross-validation was performed using five stratified folds for training and testing. The mean for each OSN subtype was computed using the training data and in the testing data the observed DV score for each OSN was compared to that of the either the mean of its respective OSN subtype in the training data or the mean of the *n*^th^ nearest OSN subtypes (as determined based on the means of each OSN subtype in the training data); the difference in median absolute errors was computed for models with *n* ranging from 0–200.

For computing deviations from the mean and area under the receiver operator curve (AUROC), equal numbers of OSNs (150) were sampled for each OSN subtype on each of 1,000 restarts, and AUROC values were calculated empirically for all pairs of OSN subtypes (_1007_C_2_ = 506,521 pairs). For comparisons to shuffled data, OSN labels were shuffled across the subsampled cells on each restart. The difference in DV scores for each cell from the respective mean of its corresponding OSN subtype was also summarized via the median absolute deviation (MAD).

#### MERFISH analyses

##### Cell type identification in MERSCOPE data

Data from different epithelial sections were aligned to a common template using STalign^15^, an approach that relies on diffeomorphic mapping. MERFISH gene expression was also visualized using Baysor^16^, in its segmentation-free configuration. In brief, local neighborhoods were evaluating by evaluating the nearest transcripts (across all genes) for every gene, reducing the dimensionality of this data via principal component analysis (PCA) and Uniform Manifold Approximation and Projection (UMAP)^17^, and converting the resulting low dimensional embedding into colors for visualization purposes. Because OSNs are small and densely packed and difficult to segment via DAPI or cell boundary staining alone, cell segmentation from the Vizgen MERlin pipeline (via cellpose) was further refined using Proseg^18^, a probabilistic approach that helps to better match transcripts to cells. The resulting log-normalized cell by gene matrix from Proseg was processed in Scanpy. PCA was performed and the top 30 PCs were kept. Leiden clustering was performed on a random subset of 100,000 cells (on a nearest neighbor graph with n=30 neighbors and leiden resolution=1.2) and the labels were transferred to all cells using the weighted nearest neighbor approache described above for the scRNA-seq data. Clusters were annotated and merged based on their expression of known marker genes (e.g for OSNs, immature precursors, and non-neuronal genes); the spatial distribution of these genes and their respective clusters were also evaluated, and matched prior knowledge. As indicated, downstream analyses that used a specific cell type (e.g. INPs or mesenchymal lamina propria cells) used only the cells and or transcripts found within those cells of a given cluster. Similar spatial segregation was also observed at the transcript level when the Baysor local neighbors for each transcript were clustered using KMeans clustering (with k=3–10 clusters).

##### OSN subtype identification

OSNs expressing single ORs were identified in an iterative process, using the spatial locations of OR transcripts. First, only transcripts in the OSN layer (identified via the clustering described above) were considered. Next, to avoid false positives from background expression (or errors in spot decoding) OR transcripts were filtered for each gene to only keep those with at least 3 other transcripts within a 10 µm radius (the average size of an OSN), and the resulting transcripts were then clustered into individual cells using the DBSCAN algorithm (eps=5 and min_samples=3). The set of OSNs was further refined by removing 0.1% of cells in which the median pairwise distance between OR transcripts within the cell was greater than 10 µm. OSNs were identified separately for each sample, using the STaligned coordinates. Four OR genes out of ∼200 showed nonspecific binding across the MOE (and were also decoded in damaged areas and areas with high autofluorescence) and were not considered for downstream analyses. Only ORs detected in at least 20 different OSNs were used analyses.

##### DV_ISH_ scores and DV scores in MERFISH data

The DV_ISH_ score was computed directly using the MERFISH-measured DV gene expression for each cell. To infer DV_ISH_ scores in OSNs, the total expression of each gene for each cell (all transcripts within a 10 µm radius of the cell centroid) were total-count normalized. Next, rather than applying cNMF directly (because cNMF was fit on scRNA-seq data and the MERFISH panel only contained a subset of all HVGs), PCA was performed on the total-count normalized expression of the DV genes that were part of the MERFISH panel and the first PC was taken to be the DV_ISH_ scores for each cell. DV_ISH_ scores smoothly varied across the MOE and matched the underlying gradients of individual DV genes. DV_ISH_ scores were calculated in a similar manner for INPs, using the set of INP cells that were located basally to OSNs that did not have detectable OR transcripts and the counts of the nearest 100 transcripts to each cell. The DV score for each cell was taken to be the DV score associated with its expressed OR, as measured via scRNA-seq.

Predictions of the DV score associated with the OR expressed in each cell were made in a cross-validated manner by holding out all the cells of a given OSN subtype at a time using either the DV_ISH_ score (linear regression with the top PC of DV expression, where PCA was fit on the training data) or the DV score associated with the ORs expressed in neighboring cells (with KNeighborsRegressor with n_neighbors=10 weighted by distance). Predictions were summarized at the OSN subtype level. Similar predictions were also observed using the nearest neighbor regression approach on a smaller sagittal dataset.

##### Alignment of MERFISH epithelial segments

The complex 3D geometry of the nose made it difficult to compare data across sections, or across turbinates directly. An alignment procedure was developed to map data into common coordinate systems. Subsections of the epithelium, like the septum, had previously been shown to contain ORs from multiple zones. Using the expression of known dorsal and ventral ORs the boundaries of all segments of the epithelium that spanned from dorsal to ventral in the STaligned coordinates were manually annotated. Then, for each segment, the ranked distance of each cell from the most dorsal vertex of the polygon boundary of the segment was computed. These positions, expressed in terms of the fractional length along each segment, were considered as the “unaligned” coordinate system for downstream analyses.

If ORs are arranged continuously across the epithelium, then their positional ranks (along the dorsal to ventral axis of each segment) should be a good proxy for distance, assuming OR ranks are preserved across segments on average even if such ranks change with different rates across segments (e.g. due to differences in the total length of segments or their location within the MOE). The mean rank for each OR across all segments was calculated and used to develop a mapping between position (i.e. the fractional length along each segment) and the mean OR rank of the OR expressed at each position and these curves were fit with piecewise linear functions (with at most 5 knots and constrained to increase monotonically constraints) via the pwlfit library. The resulting piecewise linear regression thus gives a mapping, for each segment, between the positional coordinates of that segment and the common mean OR ranks; these predicted values were taken to be the new coordinates for each cell, thereby mapping each segment into a common coordinate space. Given the monotonicity constraint of this approach, the rank ordering of cells was relatively preserved by this approach. Accordingly, the overall mean OR ranks changed little by this approach, but the resulting positions in the new coordinate system were less confounded by differences in absolute positions across segments and better captured the underlying latent “position” of each cell, and in the aligned dataset the mean OR rank at a given position was more correlated. While this annotation and alignment procedures was done independently of the DV scores, both the mean unaligned and aligned positions were well-correlated with the DV score.

The resulting unaligned or aligned positions for the cells from each segment were used for downstream analyses. AUROC analysis was performed between pairs of OSN subtypes, as described in above for the scRNA-seq data. For analyses that looked at pairwise distances between cells, only pairs of cells within the same segment were considered, and pairs were accumulated and summarized across all segments. Dorsal ORs had broader and more intermingled spatial positions, which became compressed during our alignment procedure (since the mean ranks in the dorsal part of the epithelium were often flat). To avoid any confounding results, subsequent analyses were performed, where indicated, using only cells expressing ventral class II ORs.

Regression was also used to predict the position of each cell in the aligned coordinate system using each cell’s DV_ISH_ score. Predictions were performed using a Histogram-based Gradient Boosting Regression Tree (HistGradientBoostingRegressor with loss= “absolute_error”, max_leaf_nodes=15, max_depth=15). Equal numbers of cells were subsampled for each OSN subtype (for the set of OSN subtypes with at least 20 cells). The regressor was trained with five-fold cross-validation across 250 restarts and the median absolute error in the predicted positions on each restart was computed. The observed accuracy was compared to models trained on data in which DV_ISH_ scores were shuffled across cells within an OSN subtype or data in which the positions of each cell were shuffled across all cells.

#### Comparison of DV scores with other spatial datasets

##### Bulk RNA-seq

DV scores were compared to two previous measurements, via bulk RNA-sequencing, of the spatial positions of ORs within the epithelium. One approach measured OR RNA levels in micro-dissected subregions of the epithelium^12^, whereas the other performed RNA tomography and measured OR RNA levels in cryosections across all three axes of the epithelium^19^. These spatial indices were compared to the mean DV score for OSNs expressing each OR in wild-type data (as measured via GEP usages, which are calculated using non-OR genes) for the sets of ORs detected both in these datasets and via scRNA-seq.

DV scores were also predicted based on the expression of ORs in manually-dissected “zonal” subregions of the MOE from a third dataset^20^. The raw fastq files for these experiments were obtained from the NIH sequence read archive (SRA, dataset SRP285789). Bulk RNA-seq experiments were uniformly processed and mapped to the same mouse index used for scRNA-seq experiments (Ensembl GRCm39 v105) using a Nextflow pipeline (nf-core/rnaseq) that ran STAR and RSEM to quantify the bulk RNA abundance of genes in each sample^21–23^. The DV score for each OSN subtype was predicted as a function of OR abundance in the zonal dissections using support vector machine regressors (C=500 and kernel=“rbf”), trained via 5-fold cross validation across 1,000 restarts.

##### External MOE MERFISH dataset

MERFISH data from the MOE were generated in the Dulac lab and were publicly released (CC BY 4.0) as part of the NIH’s Brain Research through Advancing Innovative Neurotechnologies (BRAIN) Initiative - Cell Census Network (BICCN)^24^. Preprocessed data, which indicated the decoded olfactory receptor gene detected in each of ∼3.3 million cells, were downloaded from the Brain Imaging Library (BIL, dataset ID: ace-bag-tin)^25^. This MERFISH dataset consisted of data from 10 mice in which coronal sections were probed with two OR gene panels that each expressed ∼500 different ORs (and ∼1,100 ORs in total). Analyses were performed on 49 sections from 8 mice, excluding 2 mice whose sections had lower quality. The epithelial sections were manually segmented as above, and the resulting 704 sections were also aligned to common coordinate system using the global OR ranks and the ranking of the cells in each segment. The resulting ranks were also well-correlated with the DV score. Lastly, to predict the DV scores of each OR in a segmentation-free manner, the same nearest-neighbor approach described above for the other MERFISH dataset was performed in which all the OSNs of a given subtype were held out and the DV scores associated with the held-out OR was predicted using the DV scores associated with the ORs expressed in the nearest neighbors spatially. Predictions were summarized at the OSN subtype level.

#### Comparison of AP scores with MERFISH data

AP scores were also evaluated at the OSN subtype level in both MERFISH datasets using the AP scores associated with each OR in the integrated scRNA-seq dataset. To relate AP scores to epithelial positions, the apical and basal axes of each epithelial segment were manually annotated and the difference in the distance to the apical and basal axis (i.e. the side of the epithelium closer to the lumen/sustentacular cell layer and the side closer to the mesenchymal and basal stem cell layer, respectively) was computed for each OSN. Only OSNs whose distances to both axes were within the bottom 99^th^ percentile were included, to exclude outliers (likely false-positives outside of the epithelium or damaged areas of the tissue). The mean apical-basal distance for each OSN subtype was computed, which were then compared to the associated DV and AP scores for each subtype. The correlation in AP scores and apical-basal distances was also evaluated separately for dorsal and ventral ORs. The apical-basal positions of a subset of ORs were also manually verified via in situ hybridizations.

#### DV scores in activity manipulations and mice with ectopic OR expression

##### Naris occlusion and environment swaps

Data from wild-type mice housed in home-cage environments, or from mice that either underwent transient naris occlusion for a week or were housed in novel olfactory environments, were all previously generated^5^. DV scores were evaluated separately for each subtype for each condition (e.g. across open vs. closed nostrils or for mice house in each environment), for OSN subtypes found in both conditions. In mice housed in home-cage environments, the DV score for each subtype was also compared to each subtype’s ES score, which summarizes the chronic activity induced by the odorants in the home-cage.

##### CNGA2 mice

scRNA-seq was performed on female mice heterozygous for the CNGA2 allele. Because CNGA2 is X-linked its expression is mosaic in female heterozygotes because individual cells inactive either the wild-type or loss of function allele. Cells with the loss of function allele were identified either based on SNPs in the *Cnga2* cDNA that distinguished it from the C57 allele, or via reads that mapped to lacZ. Cnga2 loss of function cells had decreased ES scores, consistent with the usage of that GEP being downstream of OR-driven activity.

##### Act-seq

Identification of activated OSN subtype in the Act-seq experiments was performed as previously described^5^. In brief, activated OSN subtypes were identified via immediate early gene induction, and the activation score for each subtype in the odor condition was calculated using a previous set of acutely-responsive “activation genes.” Activation scores were consistent across replicates. The DV scores for each activated OSN subtype were calculated using the entire set of cells in our integrated dataset.

##### OMP-P2 mice

P2 cells from OMP-P2 mice were identified via the expression of the P2 receptor, which was expressed in a large fraction of cells across the entire epithelium of mature mice. Two-OR cells were defined as those in which both ORs were expressed with at least 3 UMIs; many of these cells contained P2 in OMP-P2 mice, and P2 was often expressed at higher levels than the other non-P2 OR. DV scores in P2-expressing cells in the OMP-P2 mice were compared to those in our integrated dataset expressing P2 or to those expressing the non-P2 OR (for two-OR cells).

##### DV scores in “OR Swap” mice

Cells expressing each OR in the OR-swap mice were identified based on the 3’ UTR associated with each OR transcript, which was unchanged in these mice and therefore identified the genomic locus of the expressed OR for each cell. The crossed design of the OR swap experiment (in which cells express either S50 or M72 receptors from each of the S50 and M72 genomic locus) facilitated the identification of genes that followed the locus or the expressed OR. Classification was performed using cross-validated SVM models (with kernel=“rbf”) trained on the expression of either the ∼1,300 HVGs or the ∼250 DV genes in each cell (with a feature selection step that kept the top 100 genes from the training data). Models were trained to predict the genomic locus of each cell using a subset of cells from each mouse and were tested on held-out cells.

#### Gene and OR expression in differentiating OSNs

Differentiating OSNs were also identified based on the transferred cell type labels of the scANVI latent space. Both mature and immature OSNs were excluded to obtain a dataset of differentiating cells in the process of choosing ORs. Iterative subclustering was performed on the latent space of new scVI models (n_hidden: 128, n_latent: 10, n_layers: 1, gene_likelihood: nb, dropout_rate: 0.1, with a batch key for each replicate and the number of genes as a continuous covariate) trained on the top variable genes from this dataset. In this process, any remaining doublets, low-quality cells, additional non-neuronal cells that were not excluded by the above filtering steps, and the small subset of cells from the non-canonical Emx1^+^ lineage that gives rise to Gucy1b2^+^ and Gucy2d^+^ sensory neurons were removed and the scVI models were retrained until the dataset consisted of only differentiating cells. The resulting differentiation dataset contained ∼400k cells and densely captured cells from the earliest globose basal stem cells to late INP/early immature OSN cells that express single ORs. To remove the effects of the cell cycle on downstream analyses, cells scores for the G2/M and S phases were computed (scanpy’s score_genes_cell_cycle) using published marker gene sets^26^. Any gene whose expression was correlated with either G2/M or S score (pearson’s R > 0.2) was removed and the remaining 2773 were used to train a new scVI model with the same training parameters as listed above. Cells were clustered based on the nearest neighbor graph of the resulting 10-dimensional scVI latent space and this latent space was also used to reduce the dimensionality of the data to two dimensions for visualization purposes via UMAP (using a KNN-graph with 100 neighbors); these UMAPs are shown in the figures of the differentiating dataset.

##### Pseudotime analyses

Smooth differentiation trajectories were visible in the resulting UMAP projections, and thus an approach was designed to pseudotemporally order the cells. An approximate KNN graph was constructed based on the cosine similarity in the scVI latent space, using k=20 neighbors. A vector the size of the number of cells of the dataset was constructed. This vector was all zeros except for a single GBC cell, which was given a starting weight of 1. This vector was then multiplied by the kNN graph (weighting each cell by 0.1 and the influence of its nearest 20 neighbors by 0.9), renormalized from 0 to 1, and this procedure was repeated 1500 times until the values at each iteration remained stable. The resulting vector was rank-ordered, and these ranks were considered the pseudotime values for each cell; similar pseudotime values were also observed starting from other cells. A similar KNN-smoothing approach was used to smooth gene expression and GEP usages across neighboring cells, following procedures described previously^27^. Here the starting vector was the gene expression or GEP usages for each cell and the smoothing procedure was only run for 5 iterations, which results in locally-weighted values for each cell. The smoothed expression of example genes are shown in the UMAP representation, and the locally-smoothed version of each cell’s DV score was used for all downstream analyses, to help counteract the fact that fewer DV genes were expressed pre-choice and thus estimates at the cell level were noisier. Gene expression as a function of pseudotime was also fit via Generalized Additive Models using pyGAM. The alignment of DV score with axes of gene expression was assessed for cells of a given pseudotime bin (width = 0.1 and steps of 0.05) by taking the cosine distance of the DV score and the top principal components, where PCA was performed using the log-normalized expression of all OSN highly-variable genes in each pseudotime bin.

##### OR expression as a function of pseudotime

OR expression was considered with different thresholds. To identify multi-OR cells, a threshold of 1 UMI was used. Such a threshold likely induces some false positives (from e.g. background “soup” expression), given that both non-neuronal cells and GBCs sometimes expressed ORs at this 1 UMI threshold. Therefore, analyses of OR expression were restricted to cells expressing class II ORs with pseudotime values of at least 0.4, the value at which OR expression was consistently observed across cells. At subsequent pseudotime values, cells transiently expressed multiple ORs at the 1 UMI threshold. Cells with “competing” ORs were defined as those expressing multiple ORs at 3+ UMIs each (the threshold used for singular expression in mature OSNs). For cells expressing multiple ORs, the mean DV score of the co-expressed ORs was computed by weighting the DV scores associated with each OR (as measured in mature OSNs) by their expression levels (so an OR expressed at 2 UMIs would be weighted twice as much as one with 1 UMI). For individual ORs the median (and other percentiles) of the pseudotime values of all OSNs expressing that OR was computed, both for OSNs expressing each OR at any level, as well as for those expressing at either low (1–2 UMIs) or moderate levels (3+ UMIs).

To assess the correlation between each cell’s DV score and that associated with the OR, the correlation was evaluated both for cells at different differentiation stages as well as for OR expressed in those cells at either the highest, second-highest, or lowest (1-UMI) levels. Across 1,000 restarts, single ORs among the ORs expressed at each level in each cell (e.g. one of the 1-UMI ORs was picked for each cell) and the correlation was evaluated on each restart separately for cells at each differentiation stage (pre-choice with lots of low expression, cells with “competing” ORs, and cells expressing a single OR at high levels and others at lower levels).

#### Clonal lineage tracing

##### Identification of clonally-related cells in lentiviral data

To identify cells expressing barcodes and to extract the lentiviral barcode, raw sequencing reads that contained the constant start of the barcode (i.e. those that came from the amplified barcode libraries) were extracted from the BAM files. Only barcode reads with at least 10 reads for a given UMI were evaluated. The barcode library contained two variable regions (that were 14 and 30 bp long, respectively). Barcodes with hamming distances smaller than that observed between the whitelist of barcodes (3 for the 14 bp region and 5 for the 30 bp region) were collapsed and barcodes that were part of the whitelist were included in downstream analyses. Cells that expressed a single barcode (e.g. one barcode accounted for all of the barcode reads/UMIs for that cell) were considered for downstream analyses, and cells from the same replicate that expressed the same lentiviral barcode were considered as clonally-related. The resulting list of barcode-expressing cells from each experiment did not have any overlapping barcodes across mice, suggesting that the lentiviral libraries were sufficiently diverse to uniquely label each clone in each mouse. However, the overall UMIs per barcode were relatively low, and, even with the targeted amplification, many of the cells in the Venus^+^ libraries did not have any detectable barcodes. This might reflect false positives from FACS, or transcriptional silencing of barcode loci, especially given the extensive heterochromatinization of OSNs; the fluorescent signal from many infected cells was also quite weak, which is also consistent with silencing of the integrated cassette. Therefore, the modal number of cells per barcode was one, and the mode across clones with at least two cells was two. Nevertheless, results held true across clone sizes, further indicating that “small” clones are likely undersampled (due to the e.g. difficulties in barcode detection or in capturing all cells during the single-cell dissociated), but not fundamentally different than those with more cells. Additionally, the OSNs identified following methimazole-induced regeneration expressed single ORs at normal levels and had GEP usages that was similar to that of control OSNs that expressed the same OR. Similar results were observed for the lentiviral-infected cells, which expressed a wide array of ORs.

##### Clonal restrictions

Clones expressing at least two OSNs, with each singularly expressing any OR or only class II ORs, as indicated, were considered for downstream analyses. DV scores for the expressed OR were calculated by using the DV score associated with that OR in the entire integrate dataset. The clonal mean OR DV was calculated by averaging the DV scores across clonally-related OSNs. Differences between the DV scores of clonally related ORs were calculated between all pairs of cells from the same clone and were either summarized across clones or across all pairs. Shuffles for all analyses were performed by shuffling clonal labels across cells. Clonal data was simulated based on the observed clonal restriction in DV scores by sampling, for each of the cells in the observed clones, cells from the wild-type data whose ORs had DV scores that matched the observed *t-*distribution of deltas from the clonal mean (e.g. ORs with DV scores similar to the clonal mean DV score had the highest probability of being chosen); the probabilities were also scaled by the observed frequency of each OR in the wild-type data. The percent of clones that had multiple cells that expressed the same OR (and therefore fewer number of ORs than cells) were then analyzed for both the observed and simulated data.

DV scores were also evaluated at the cell level, in which each cell’s DV scores was obtained by taking the difference in GEP_Dorsal_ and GEP_Ventral_ usage irrespective of the expressed OR. To test whether restrictions at the cell level were stronger than that at the OR level, the MAD in cell DV scores within a clone was compared the MAD obtained when DV scores were shuffled across cells of a given OSN subtype (irrespective of their clonal identity) or shuffled across all cells.

INP gene expression was related to OR DV scores, using the subset of clones that contained both INP cells and OSNs. For the cells in these clones support vector machine regression models (SVR with C=50 and kernel=“rbf”) were fit on the INP gene expression (where the input to the linear models was the top 250 genes in the training data, reduced to 30 dimensions via PCA) to predict the mean OR DV score of the clonally-related ORs expressed in the sister cells. The regression was performed via cross-validation, leaving out all the cells from a clone; additionally, the top genes and PCA transformation was identified using only the cells of the training data to avoid any data leakage. Lastly, comparisons to the clonal lineage data were performed only for clones expressing at least two ventral class II ORs that were also detected in the MERFISH data.

#### Retinoic acid manipulations

Data from adult mice given either the ALDH1A2 inhibitor WIN 18,446 (RA inh.) or all trans-Retinoic Acid (atRA) during methimazole-induced regeneration were analyzing relative to their respective vehicle controls. Cell types were identified via clustering the latent space of scVI models and cNMF GEP loadings were applied to each cell, as described above for the integrated dataset. Analyses were either performed at the cell level, using the observed DV score for each cell, or at the OR level, using the DV score associated with each OR as measured using all cells in the integrated datasets. To assess distributional shifts, the median DV score across all cells or the number of cells expressing ORs with given associated DV scores were compared between the data from each condition and its respective control; given that fewer cells express the most ventral ORs to begin with, the changes in OSN frequency were also expressed as log_2_ fold-changes. Because RA manipulations changed the overall distributions of cells without altering the relationship between cell identity and OR choice, distributional shifts were observed when evaluating all OSN subtypes at the cell level, but minimal changes were observed across conditions for cells of a given OSN subtype. Changes in gene expression were evaluated by z-scoring the expression of each DV gene across all cells and the mean change in z-scored expression was evaluated for cells from drug versus control conditions. Analyses were performed on mature OSNs expressing single ORs, except where, as noted, INPs were used to evaluate changes in DV scores and OR expression in differentiating cells pre-choice.

#### HP1 swap experiments

scRNA-seq was performed as described above in the HP1 swap and control animals (i.e. with or without the Foxg1-Cre allele) in young (4–6 week) animals of either sex. Cell type identification, DV scores, and changes to the cell and DV scores in the HP1 swap animals were evaluated in a similar manner to those in the RA manipulations. To assess changes in OR choice for cells of a given DV scores, cells were binned based on their cell DV scores and the percent-normalized DV score associated with each OR was computed for cells in each bin. The cell vs OR DV score mapping was evaluated using robust linear regression (HuberRegressor), and equal numbers of cells were sampled for each OSN subtypes, for subtypes detected in both control and HP1 swap animals. To evaluate changes in the cell to OR DV score mapping, a nearest neighbor approach was used, and a nearest neighbor regression model (KNeighborsRegressor) was fit on all cells from the control mice to predict the DV score associated with the ORs expressing in cells of a given DV score. This model was then applied to the cell DV scores of the HP1 mice, and the residuals from this model, which capture the difference in the associated percentile-normalized DV score of the OR detected in each HP1 swap cell with that predicted based on each cell’s DV score, were evaluated and summarized for HP1 swap cells of a given DV score.

#### Allelic expression in C57/CAST F_1_ hybrid animals

scRNA-seq was performed in F_1_ hybrid animals generated by crossing wild-derived CAST/EiJ female mice with C57BL/6J males. CAST/EiJ mice have on average single nucleotide polymorphisms (SNPs) every 150 bp. Because OR expression is monogenic and monoallelic, the reads that mapped to the chosen OR were used in each mature OSN to infer the strain of the chosen OR. The GRCm39 coordinates of SNPs that distinguished CAST/EiJ and C57BL/6J were obtained from the Mouse Genome Project^28,29^, and SNPs in ORs of interested were also confirmed by evaluating bulk and scRNA-seq data from homozygous CAST/EiJ animals, generated in house and in past work (Ibarria-Soria). For each cell, any reads (with MAPQ > 30) that overlapped with homozygous SNPs in CAST/EiJ mice for its chosen OR were evaluated to assess what fraction of reads expressed the CAST/EiJ or C57BL/6J variant. Cells that had reads that overlapped at least one SNP and in which at least 80% of such reads came from a single allele (mean 99.7% of reads) were used for downstream analyses. Similar results were also obtained using Demuxalot to demultiplex the strain of each cell via the set of SNPs across all OR genes^30^. The strain of the chosen OR was able to be inferred in 81% of cells. The remaining 19% of cells had too few reads that overlapped SNPs, likely because they expressed ORs with minimal variation between strains in their 3’ UTRs. SNPs that led to missense mutations were annotated via SnpEff^31^.

OSN subtypes with significant changes in GEP usages were evaluated empirically using permutation testing, and subtypes with a mean change between strains larger than 1% of shuffles for that GEP were considered as significant. Similar changes were observed across F_0_ and F_1_ animals, as well as across individual samples; however, some OSN subtypes had few cells for either CAST/C57 (as OR frequency varies across strains) and were likely underpowered.

#### ChIP-seq and CUT&RUN data analyses

ChIP-seq data were previously generated and were reanalyzed^20,32^. The raw fastq files for ChIP-seq data for H3K9me3 and H3k79me3 marks were obtained from the SRA (datasets SRP285789 and SRP096660) and were reprocessed with the Ensembl GRCm39 version 105 genome, using the default parameters of the nf-core/chipseq Nextflow pipeline (v2.0.0)^21^. As part of this pipeline, reads were mapped to the genome with BWA, and the normalized read density was summarized at base-pair resolution with UCSC-bedGraphToBigWig. The density of reads across the entire coding and non-coding regions of each OR for each sample were averaged, and the reads for each OR locus (for class II ORs) were correlated with the DV score of OSNs singular expressed that same OR.

##### CUT&RUN in F_1_ hybrid animals

Demultiplexed fastq files from CUT&RUN samples were mapped to the Ensembl GRCm39 v105 genome via Bowtie2 (-I 10 -X 700 --no-mixed --end-to-end --no-discordant --very-sensitive)^33^. Spike-in reads from E. coli DNA were mapped to the E. coli genome. However, because F_1_ animals have within-animal controls (the reads mapped to each allele), the number of reads that mapped to the E. coli genome were not used for normalization. Duplicates were not removed, and data were combined across replicates. As in the F_1_ scRNA-seq data, the allele of each read was inferred for reads overlapping SNPs that distinguished CAST/EiJ and C57BL/6J. The number of reads that mapped to each allele were then summarized for given ORs across the entire genomic coordinates for that gene (i.e. from TSS to TES).

#### OR promoter analyses

The presence of motifs in the 1kb upstream of the TSS for each OR (i.e. the promoter region) were evaluated using HOMER^34^, using the default motif file and threshold provided by HOMER for LHX2, NFI, EBF, and RARa. OR promoters were enriched for EBF binding motifs in the 750– 1000 bp upstream of the TSS, though the enrichment of EBF motifs in dorsal ORs and NFI/RAR motifs in ventral ORs was observed when evaluating the entire promoter region (since the majority of motif instances were in the 750–1000 bp region).

#### Olfactory bulb glomerular positions

The olfactory bulb glomerular map was generated via spatial transcriptomics using the 10x Visium platform, as recently described by Klimpert and colleagues^35^. In brief, putative glomeruli were identified using a Bayesian approach that used the expression of individual OR genes across the entire 3D volume to infer the location and number of glomeruli for each OR. The outputs of this model were manually curated to keep high-confidence identified glomeruli, and an axis of symmetry was identified that separated glomeruli from the medial and lateral domains. Importantly, the two hemibulbs are also symmetric, and once aligned to each other sister glomeruli expressing the same OR were found on average 230 µm away from each other (∼1–2 glomerular lengths), reflecting a combination of both biological variability and spatial errors in the glomerular detection and alignment procedures; such deviations are also similar to those from previous estimates^36^.

Glomeruli were identified based on OR gene expression and could thus be linked to the scRNA-seq or MERFISH data for the associated OSN subtype expressing that OR. First, OB positions were used to predict the scRNA-seq measured GEP scores for each OSN subtypes, via a cross-validated support vector machine regression model (SVR with C=150 and “rbf” kernel). Predicting GEP usages from glomerular positions is easier than the reverse, due to the higher dimensionality of the 3D positions and due to the fact that the DV score mapped onto an axis in the OB that was not fully aligned with the cartesian D-V axis but rather correlated with both the D-V and A-P position of glomeruli. Similar results were also obtained through canonical correlation analysis, which identified optimal rotations to align the GEP and glomerular spaces, further demonstrating that the DV score was well-aligned (rho > 0.9) with the top canonical correlation axis. Second, predictions of the 3D glomerular position for each OSN subtype was performed separately for glomeruli in each domain. Elastic-net-regularized linear regression models (alpha=1, l1_ratio=0.9) were trained on either GEP usage (e.g. GEP_Dorsal_, GEP_Ventral_, and the DV score) or all ∼1,300 variable genes to predict the glomerular position along each axis and the 3D error of these predictions were evaluated for each glomerulus. For both models, five-fold cross validation was performed; this procedure was repeated 100 times, and the distribution of the accuracies and or errors across restart was reported.

#### Software used for data analysis

Single-cell analyses were performed in python (versions 3.6–3.9) using the Scanpy and scvi-tools packages^37,38^, as well as custom-written scripts using the open-source python scientific stack (pysam, SciPy, NumPy, scikit-learn, umap-learn, pandas, statsmodels, matplotlib, seaborn, and numba). cNMF was performed using modified versions of the code from https://github.com/dylkot/cNMF.

#### Statistical testing

Hypothesis testing was performed using non-parametric statistical tests, except in the cases where p-values were calculated empirically using resampling-based permutation tests. The precision of sample statistics and regression trend lines were evaluated using bootstrapping, and, except where noted, plots and error bars depict the mean and the 95% confidence intervals of the mean across 1,000-10,000 bootstraps. Throughout the paper, a non-parametric version of the box plot (also known as a letter-value plot) was used to represent multiple quantiles and tails of large distributions of data (e.g. to summarize across OSN subtypes or sets of ORs or OR pairs) in an agnostic way that does require setting bandwidth parameters as in violin plots or kernel density estimates. Like a conventional box plot, the largest box represents the interquartile range (25–75 percentile) and the median (dotted-line). Subsequent boxes recursively represent exponentially-smaller quantiles (the 12.5–25 and 75–87.5 percentiles, then 6.25–12.5 and 87.5–93.75 percentiles, then 3.125-6.25 and 93.75–96.875 percentiles, and so forth.

## References

1. Kaas, J.H. (1997). Topographic maps are fundamental to sensory processing. Brain Res Bull 44, 107–112. 10.1016/s0361-9230(97)00094-4.

2. Knudsen, E.I., Lac, S., and Esterly, S.D. (1987). Computational Maps in the Brain. Annual Review of Neuroscience 10, 41–65. 10.1146/annurev.ne.10.030187.000353.

3. Brewer, A.A., and Barton, B. (2016). Maps of the Auditory Cortex. Annual Review of Neuroscience 39, 385–407. 10.1146/annurev-neuro-070815-014045.

4. Nauhaus, I., and Nielsen, K.J. (2014). Building maps from maps in primary visual cortex. Current Opinion in Neurobiology 24, 1–6. 10.1016/j.conb.2013.08.007.

5. Abraira, Victoria E., and Ginty, David D. (2013). The Sensory Neurons of Touch. Neuron 79, 618–639. 10.1016/j.neuron.2013.07.051.

6. Barnes, I.H.A., Ibarra-Soria, X., Fitzgerald, S., Gonzalez, J.M., Davidson, C., Hardy, M.P., Manthravadi, D., Van Gerven, L., Jorissen, M., Zeng, Z., et al. (2020). Expert curation of the human and mouse olfactory receptor gene repertoires identifies conserved coding regions split across two exons. BMC Genomics 21, 196. 10.1186/s12864-020-6583-3.

7. Monahan, K., and Lomvardas, S. (2015). Monoallelic Expression of Olfactory Receptors. Annual Review of Cell and Developmental Biology 31, 721–740. 10.1146/annurev-cellbio-100814-125308.

8. Murthy, V.N. (2011). Olfactory Maps in the Brain. Annual Review of Neuroscience 34, 233–258. 10.1146/annurev-neuro-061010-113738.

9. Imai, T., Sakano, H., and Vosshall, L.B. (2010). Topographic mapping--the olfactory system. Cold Spring Harb Perspect Biol 2, a001776. 10.1101/cshperspect.a001776.

10. Brann, D.H., and Datta, S.R. (2020). Finding the Brain in the Nose. Annual Review of Neuroscience 43, 277–295. 10.1146/annurev-neuro-102119-103452.

11. O’Leary, D.D.M., Yates, P.A., and McLaughlin, T. (1999). Molecular Development of Sensory Maps: Representing Sights and Smells in the Brain. Cell 96, 255–269. 10.1016/S0092-8674(00)80565-6.

12. Norlin, E.M., Alenius, M., Gussing, F., Hägglund, M., Vedin, V., and Bohm, S. (2001). Evidence for Gradients of Gene Expression Correlating with Zonal Topography of the Olfactory Sensory Map. Molecular and Cellular Neuroscience 18, 283–295. 10.1006/mcne.2001.1019.

13. Zhang, X., and Firestein, S. (2002). The olfactory receptor gene superfamily of the mouse. Nature Neuroscience 5, 124–133. 10.1038/nn800.

14. Bear, Daniel M., Lassance, J.-M., Hoekstra, Hopi E., and Datta, Sandeep R. (2016). The Evolving Neural and Genetic Architecture of Vertebrate Olfaction. Current Biology 26, R1039–R1049. 10.1016/j.cub.2016.09.011.

15. Coleman, J.H., Lin, B., Louie, J.D., Peterson, J., Lane, R.P., and Schwob, J.E. (2019). Spatial Determination of Neuronal Diversification in the Olfactory Epithelium. The Journal of Neuroscience 39, 814–832. 10.1523/jneurosci.3594-17.2018.

16. Schwob, J., and Gottlieb, D. (1986). The primary olfactory projection has two chemically distinct zones. The Journal of Neuroscience 6, 3393–3404. 10.1523/jneurosci.06-11-03393.1986.

17. Zapiec, B., and Mombaerts, P. (2020). The Zonal Organization of Odorant Receptor Gene Choice in the Main Olfactory Epithelium of the Mouse. Cell Reports 30, 4220–4234.e4225. 10.1016/j.celrep.2020.02.110.

18. Ressler, K.J., Sullivan, S.L., and Buck, L.B. (1993). A zonal organization of odorant receptor gene expression in the olfactory epithelium. Cell 73, 597–609. 10.1016/0092-8674(93)90145-G.

19. Vassar, R., Ngai, J., and Axel, R. (1993). Spatial segregation of odorant receptor expression in the mammalian olfactory epithelium. Cell 74, 309–318. 10.1016/0092-8674(93)90422-M.

20. Miyamichi, K., Serizawa, S., Kimura, H.M., and Sakano, H. (2005). Continuous and Overlapping Expression Domains of Odorant Receptor Genes in the Olfactory Epithelium Determine the Dorsal/Ventral Positioning of Glomeruli in the Olfactory Bulb. The Journal of Neuroscience 25, 3586–3592. 10.1523/jneurosci.0324-05.2005.

21. Tan, L., and Xie, X.S. (2018). A Near-Complete Spatial Map of Olfactory Receptors in the Mouse Main Olfactory Epithelium. Chemical Senses 43, 427–432. 10.1093/chemse/bjy030.

22. Ruiz Tejada Segura, M.L., Abou Moussa, E., Garabello, E., Nakahara, T.S., Makhlouf, M., Mathew, L.S., Wang, L., Valle, F., Huang, S.S.Y., Mainland, J.D., et al. (2022). A 3D transcriptomics atlas of the mouse nose sheds light on the anatomical logic of smell. Cell Reports 38. 10.1016/j.celrep.2022.110547.

23. Sullivan, S.L., Ressler, K.J., and Buck, L.B. (1995). Spatial patterning and information coding in the olfactory system. Current Opinion in Genetics & Development 5, 516–523. 10.1016/0959-437X(95)90057-N.

24. Iwema, C.L., Fang, H., Kurtz, D.B., Youngentob, S.L., and Schwob, J.E. (2004). Odorant Receptor Expression Patterns Are Restored in Lesion-Recovered Rat Olfactory Epithelium. The Journal of Neuroscience 24, 356–369. 10.1523/jneurosci.1219-03.2004.

25. Bozza, T., Vassalli, A., Fuss, S., Zhang, J.-J., Weiland, B., Pacifico, R., Feinstein, P., and Mombaerts, P. (2009). Mapping of Class I and Class II Odorant Receptors to Glomerular Domains by Two Distinct Types of Olfactory Sensory Neurons in the Mouse. Neuron 61, 220–233. 10.1016/j.neuron.2008.11.010.

26. Pacifico, R., Dewan, A., Cawley, D., Guo, C., and Bozza, T. (2012). An Olfactory Subsystem that Mediates High-Sensitivity Detection of Volatile Amines. Cell Reports 2, 76–88. 10.1016/j.celrep.2012.06.006.

27. Shykind, B.M., Rohani, S.C., O’Donnell, S., Nemes, A., Mendelsohn, M., Sun, Y., Axel, R., and Barnea, G. (2004). Gene Switching and the Stability of Odorant Receptor Gene Choice. Cell 117, 801–815. 10.1016/j.cell.2004.05.015.

28. Oka, Y., Kobayakawa, K., Nishizumi, H., Miyamichi, K., Hirose, S., Tsuboi, A., and Sakano, H. (2003). O-MACS, a novel member of the medium-chain acyl-CoA synthetase family, specifically expressed in the olfactory epithelium in a zone-specific manner. European Journal of Biochemistry 270, 1995–2004. 10.1046/j.1432-1033.2003.03571.x.

29. Yoshihara, Y., Kawasaki, M., Tamada, A., Fujita, H., Hayashi, H., Kagamiyama, H., and Mori, K. (1997). OCAM: A New Member of the Neural Cell Adhesion Molecule Family Related to Zone-to-Zone Projection of Olfactory and Vomeronasal Axons. The Journal of Neuroscience 17, 5830–5842. 10.1523/jneurosci.17-15-05830.1997.

30. Kobayakawa, K., Kobayakawa, R., Matsumoto, H., Oka, Y., Imai, T., Ikawa, M., Okabe, M., Ikeda, T., Itohara, S., Kikusui, T., et al. (2007). Innate versus learned odour processing in the mouse olfactory bulb. Nature 450, 503–508. 10.1038/nature06281.

31. Tan, L., Zong, C., and Xie, X.S. (2013). Rare event of histone demethylation can initiate singular gene expression of olfactory receptors. Proceedings of the National Academy of Sciences 110, 21148–21152. 10.1073/pnas.1321511111.

32. Bashkirova, E.V., Klimpert, N., Monahan, K., Campbell, C.E., Osinski, J., Tan, L., Schieren, I., Pourmorady, A., Stecky, B., Barnea, G., et al. (2023). Opposing, spatially-determined epigenetic forces impose restrictions on stochastic olfactory receptor choice. eLife 12, RP87445. 10.7554/eLife.87445.

33. Tan, L., Li, Q., and Xie, X.S. (2015). Olfactory sensory neurons transiently express multiple olfactory receptors during development. Molecular Systems Biology 11, 844. 10.15252/msb.20156639.

34. Hanchate, N.K., Kondoh, K., Lu, Z., Kuang, D., Ye, X., Qiu, X., Pachter, L., Trapnell, C., and Buck, L.B. (2015). Single-cell transcriptomics reveals receptor transformations during olfactory neurogenesis. Science 350, 1251–1255. 10.1126/science.aad2456.

35. Monahan, K., Horta, A., and Lomvardas, S. (2019). LHX2- and LDB1-mediated trans interactions regulate olfactory receptor choice. Nature 565, 448–453. 10.1038/s41586-018-0845-0.

36. Yusuf, N., and Monahan, K. (2024). Epigenetic programming of stochastic olfactory receptor choice. genesis *62*, e23593. 10.1002/dvg.23593.

37. Bashkirova, E., and Lomvardas, S. (2019). Olfactory receptor genes make the case for inter-chromosomal interactions. Current Opinion in Genetics & Development 55, 106–113. 10.1016/j.gde.2019.07.004.

38. Zapiec, B., and Mombaerts, P. (2015). Multiplex assessment of the positions of odorant receptor-specific glomeruli in the mouse olfactory bulb by serial two-photon tomography. Proceedings of the National Academy of Sciences 112, E5873–E5882. 10.1073/pnas.1512135112.

39. Soucy, E.R., Albeanu, D.F., Fantana, A.L., Murthy, V.N., and Meister, M. (2009). Precision and diversity in an odor map on the olfactory bulb. Nature Neuroscience 12, 210–220. 10.1038/nn.2262.

40. Strotmann, J., Conzelmann, S., Beck, A., Feinstein, P., Breer, H., and Mombaerts, P. (2000). Local Permutations in the Glomerular Array of the Mouse Olfactory Bulb. The Journal of Neuroscience 20, 6927–6938. 10.1523/jneurosci.20-18-06927.2000.

41. Mombaerts, P., Wang, F., Dulac, C., Chao, S.K., Nemes, A., Mendelsohn, M., Edmondson, J., and Axel, R. (1996). Visualizing an Olfactory Sensory Map. Cell 87, 675–686. 10.1016/S0092-8674(00)81387-2.

42. Mori, K., and Sakano, H. (2011). How Is the Olfactory Map Formed and Interpreted in the Mammalian Brain? Annual Review of Neuroscience 34, 467–499. 10.1146/annurev-neuro-112210-112917.

43. Ma, M., and Shepherd, G.M. (2000). Functional mosaic organization of mouse olfactory receptor neurons. Proceedings of the National Academy of Sciences 97, 12869–12874. 10.1073/pnas.220301797.

44. Takahashi, H., Yoshihara, S.-i., Nishizumi, H., and Tsuboi, A. (2010). Neuropilin-2 is required for the proper targeting of ventral glomeruli in the mouse olfactory bulb. Molecular and Cellular Neuroscience 44, 233–245. 10.1016/j.mcn.2010.03.010.

45. Cho, J.H., Prince, J.E.A., Cutforth, T., and Cloutier, J.-F. (2011). The Pattern of Glomerular Map Formation Defines Responsiveness to Aversive Odorants in Mice. The Journal of Neuroscience 31, 7920–7926. 10.1523/jneurosci.2460-10.2011.

46. Takeuchi, H., Inokuchi, K., Aoki, M., Suto, F., Tsuboi, A., Matsuda, I., Suzuki, M., Aiba, A., Serizawa, S., Yoshihara, Y., et al. (2010). Sequential Arrival and Graded Secretion of Sema3F by Olfactory Neuron Axons Specify Map Topography at the Bulb. Cell 141, 1056–1067. 10.1016/j.cell.2010.04.041.

47. Bozza, T., Feinstein, P., Zheng, C., and Mombaerts, P. (2002). Odorant Receptor Expression Defines Functional Units in the Mouse Olfactory System. The Journal of Neuroscience 22, 3033–3043. 10.1523/jneurosci.22-08-03033.2002.

48. Wang, F., Nemes, A., Mendelsohn, M., and Axel, R. (1998). Odorant Receptors Govern the Formation of a Precise Topographic Map. Cell 93, 47–60. 10.1016/S0092-8674(00)81145-9.

49. Sakano, H. (2020). Developmental regulation of olfactory circuit formation in mice. Development, Growth & Differentiation 62, 199–213. 10.1111/dgd.12657.

50. Nakashima, A., Takeuchi, H., Imai, T., Saito, H., Kiyonari, H., Abe, T., Chen, M., Weinstein, Lee S., Yu, C.R., Storm, Daniel R., et al. (2013). Agonist-Independent GPCR Activity Regulates Anterior-Posterior Targeting of Olfactory Sensory Neurons. Cell 154, 1314–1325. 10.1016/j.cell.2013.08.033.

51. Feinstein, P., and Mombaerts, P. (2004). A Contextual Model for Axonal Sorting into Glomeruli in the Mouse Olfactory System. Cell 117, 817–831. 10.1016/j.cell.2004.05.011.

52. Serizawa, S., Miyamichi, K., Takeuchi, H., Yamagishi, Y., Suzuki, M., and Sakano, H. (2006). A Neuronal Identity Code for the Odorant Receptor-Specific and Activity-Dependent Axon Sorting. Cell 127, 1057–1069. 10.1016/j.cell.2006.10.031.

53. Gussing, F., and Bohm, S. (2004). NQO1 activity in the main and the accessory olfactory systems correlates with the zonal topography of projection maps. Eur. J. Neurosci. 19, 2511–2518. 10.1111/j.0953-816x.2004.03331.x.

54. Tsukahara, T., Brann, D.H., Pashkovski, S.L., Guitchounts, G., Bozza, T., and Datta, S.R. (2021). A transcriptional rheostat couples past activity to future sensory responses. Cell 184, 6326–6343.e6332. 10.1016/j.cell.2021.11.022.

55. Kotliar, D., Veres, A., Nagy, M.A., Tabrizi, S., Hodis, E., Melton, D.A., and Sabeti, P.C. (2019). Identifying gene expression programs of cell-type identity and cellular activity with single-cell RNA-Seq. eLife 8, e43803. 10.7554/eLife.43803.

56. Chen, K.H., Boettiger, A.N., Moffitt, J.R., Wang, S., and Zhuang, X. (2015). Spatially resolved, highly multiplexed RNA profiling in single cells. Science 348, aaa6090. 10.1126/science.aaa6090.

57. Brunet, L.J., Gold, G.H., and Ngai, J. (1996). General Anosmia Caused by a Targeted Disruption of the Mouse Olfactory Cyclic Nucleotide–Gated Cation Channel. Neuron 17, 681–693. 10.1016/s0896-6273(00)80200-7.

58. Cichy, A., Shah, A., Dewan, A., Kaye, S., and Bozza, T. (2019). Genetic Depletion of Class I Odorant Receptors Impacts Perception of Carboxylic Acids. Current Biology 29, 2687–2697.e2684. 10.1016/j.cub.2019.06.085.

59. Fleischmann, A., Abdus-Saboor, I., Sayed, A., and Shykind, B. (2013). Functional Interrogation of an Odorant Receptor Locus Reveals Multiple Axes of Transcriptional Regulation. PLOS Biology 11, e1001568. 10.1371/journal.pbio.1001568.

60. Fletcher, R.B., Das, D., Gadye, L., Street, K.N., Baudhuin, A., Wagner, A., Cole, M.B., Flores, Q., Choi, Y.G., Yosef, N., et al. (2017). Deconstructing Olfactory Stem Cell Trajectories at Single-Cell Resolution. Cell Stem Cell 20, 817–830.e818. 10.1016/j.stem.2017.04.003.

61. Hussainy, M., Korsching, S.I., and Tresch, A. (2022). Pseudotime analysis reveals novel regulatory factors for multigenic onset and monogenic transition of odorant receptor expression. Scientific Reports 12, 16183. 10.1038/s41598-022-20106-w.

62. Bergström, U., Giovanetti, A., Piras, E., and Brittebo, E.B. (2003). Methimazole-Induced Damage in the Olfactory Mucosa: Effects on Ultrastructure and Glutathione Levels. Toxicol. Pathol. 31, 379–387. 10.1080/01926230390201101.

63. Whitesides, J., Hall, M., Anchan, R., and LaMantia, A.S. (1998). Retinoid signaling distinguishes a subpopulation of olfactory receptor neurons in the developing and adult mouse. Journal of Comparative Neurology 394, 445–461. 10.1002/(sici)1096-9861(19980518)394:4<445::aid-cne4>3.0.co;2-1.

64. Asson-Batres, M.A., and Smith, W.B. (2006). Localization of retinaldehyde dehydrogenases and retinoid binding proteins to sustentacular cells, glia, Bowman’s gland cells, and stroma: Potential sites of retinoic acid synthesis in the postnatal rat olfactory organ. Journal of Comparative Neurology 496, 149–171. 10.1002/cne.20904.

65. Peluso, C.E., Jang, W., Dräger, U.C., and Schwob, J.E. (2012). Differential expression of components of the retinoic acid signaling pathway in the adult mouse olfactory epithelium. Journal of Comparative Neurology 520, 3707–3726. 10.1002/cne.23124.

66. Kanata, E., Duffié, R., and Schulz, E.G. (2024). Establishment and maintenance of random monoallelic expression. Development 151, dev201741. 10.1242/dev.201741.

67. Clowney, E.J., LeGros, Mark A., Mosley, Colleen P., Clowney, Fiona G., Markenskoff-Papadimitriou, Eirene C., Myllys, M., Barnea, G., Larabell, Carolyn A., and Lomvardas, S. (2012). Nuclear Aggregation of Olfactory Receptor Genes Governs Their Monogenic Expression. Cell 151, 724–737. 10.1016/j.cell.2012.09.043.

68. Le Gros, Mark A., Clowney, E.J., Magklara, A., Yen, A., Markenscoff-Papadimitriou, E., Colquitt, B., Myllys, M., Kellis, M., Lomvardas, S., and Larabell, Carolyn A. (2016). Soft X-Ray Tomography Reveals Gradual Chromatin Compaction and Reorganization during Neurogenesis In&#xa0;Vivo. Cell Reports 17, 2125–2136. 10.1016/j.celrep.2016.10.060.

69. Bosch-Presegué, L., Raurell-Vila, H., Thackray, J.K., González, J., Casal, C., Kane-Goldsmith, N., Vizoso, M., Brown, J.P., Gómez, A., Ausió, J., et al. (2017). Mammalian HP1 Isoforms Have Specific Roles in Heterochromatin Structure and Organization. Cell Reports 21, 2048–2057. 10.1016/j.celrep.2017.10.092.

70. Hiragami-Hamada, K., Soeroes, S., Nikolov, M., Wilkins, B., Kreuz, S., Chen, C., De La Rosa-Velázquez, I.A., Zenn, H.M., Kost, N., Pohl, W., et al. (2016). Dynamic and flexible H3K9me3 bridging via HP1β dimerization establishes a plastic state of condensed chromatin. Nature Communications 7, 11310. 10.1038/ncomms11310.

71. Keane, T.M., Goodstadt, L., Danecek, P., White, M.A., Wong, K., Yalcin, B., Heger, A., Agam, A., Slater, G., Goodson, M., et al. (2011). Mouse genomic variation and its effect on phenotypes and gene regulation. Nature 477, 289–294. 10.1038/nature10413.

72. Klimpert, N., Kollo, M., Brann, D.H., Tan, C., Barry, D., Ma, Y., Schaefer, A.T., and Fleischmann, A. (2025). 3D spatial transcriptomics reveals the molecular domain structure of the mouse olfactory bulb. bioRxiv, 2025.2002.2019.639192. 10.1101/2025.02.19.639192.

73. Imai, T., Suzuki, M., and Sakano, H. (2006). Odorant Receptor□-Derived cAMP Signals Direct Axonal Targeting. Science 314, 657–661. 10.1126/science.1131794.

74. Pourmorady, A.D., Bashkirova, E.V., Chiariello, A.M., Belagzhal, H., Kodra, A., Duffié, R., Kahiapo, J., Monahan, K., Pulupa, J., Schieren, I., et al. (2024). RNA-mediated symmetry breaking enables singular olfactory receptor choice. Nature 625, 181–188. 10.1038/s41586-023-06845-4.

75. Pourmorady, A., and Lomvardas, S. (2022). Olfactory receptor choice: a case study for gene regulation in a multi-enhancer system. Current Opinion in Genetics & Development 72, 101–109. 10.1016/j.gde.2021.11.003.

76. Lomvardas, S., Barnea, G., Pisapia, D.J., Mendelsohn, M., Kirkland, J., and Axel, R. (2006). Interchromosomal Interactions and Olfactory Receptor Choice. Cell 126, 403–413. 10.1016/j.cell.2006.06.035.

77. Wu, H., Zhang, J., Jian, F., Chen, J.P., Zheng, Y., Tan, L., and Sunney Xie, X. (2024). Simultaneous single-cell three-dimensional genome and gene expression profiling uncovers dynamic enhancer connectivity underlying olfactory receptor choice. Nature Methods 21, 974–982. 10.1038/s41592-024-02239-0.

78. Tan, L., Xing, D., Daley, N., and Xie, X.S. (2019). Three-dimensional genome structures of single sensory neurons in mouse visual and olfactory systems. Nature Structural & Molecular Biology 26, 297–307. 10.1038/s41594-019-0205-2.

79. Lyons, David B., Magklara, A., Goh, T., Sampath, S.C., Schaefer, A., Schotta, G., and Lomvardas, S. (2014). Heterochromatin-Mediated Gene Silencing Facilitates the Diversification of Olfactory Neurons. Cell Reports 9, 884–892. 10.1016/j.celrep.2014.10.001.

80. Qasba, P., and Reed, R.R. (1998). Tissue and Zonal-Specific Expression of an Olfactory Receptor Transgene. The Journal of Neuroscience 18, 227–236. 10.1523/jneurosci.18-01-00227.1998.

81. Rothman, A., Feinstein, P., Hirota, J., and Mombaerts, P. (2005). The promoter of the mouse odorant receptor gene M71. Molecular and Cellular Neuroscience 28, 535–546. 10.1016/j.mcn.2004.11.006.

82. Vassalli, A., Rothman, A., Feinstein, P., Zapotocky, M., and Mombaerts, P. (2002). Minigenes Impart Odorant Receptor-Specific Axon Guidance in the Olfactory Bulb. Neuron 35, 681–696. 10.1016/S0896-6273(02)00793-6.

83. Zhang, Y.-Q., Breer, H., and Strotmann, J. (2007). Promotor elements governing the clustered expression pattern of odorant receptor genes. Molecular and Cellular Neuroscience 36, 95–107. 10.1016/j.mcn.2007.06.005.

84. Tsuboi, A., Yoshihara, S.-i., Yamazaki, N., Kasai, H., Asai-Tsuboi, H., Komatsu, M., Serizawa, S., Ishii, T., Matsuda, Y., Nagawa, F., and Sakano, H. (1999). Olfactory Neurons Expressing Closely Linked and Homologous Odorant Receptor Genes Tend to Project Their Axons to Neighboring Glomeruli on the Olfactory Bulb. The Journal of Neuroscience 19, 8409–8418. 10.1523/jneurosci.19-19-08409.1999.

85. Li, H., Li, T., Horns, F., Li, J., Xie, Q., Xu, C., Wu, B., Kebschull, J.M., McLaughlin, C.N., Kolluru, S.S., et al. (2020). Single-Cell Transcriptomes Reveal Diverse Regulatory Strategies for Olfactory Receptor Expression and Axon Targeting. Current Biology 30, 1189–1198.e1185. 10.1016/j.cub.2020.01.049.

86. Ray, A., Naters, W.v.d.G.v., Shiraiwa, T., and Carlson, J.R. (2007). Mechanisms of Odor Receptor Gene Choice in Drosophila. Neuron 53, 353–369. 10.1016/j.neuron.2006.12.010.

87. Hägglund, M., Berghard, A., Strotmann, J.r., and Bohm, S. (2006). Retinoic Acid Receptor-Dependent Survival of Olfactory Sensory Neurons in Postnatal and Adult Mice. The Journal of Neuroscience 26, 3281–3291. 10.1523/jneurosci.4955-05.2006.

88. Login, H., Håglin, S., Berghard, A., and Bohm, S. (2015). The Stimulus-Dependent Gradient of Cyp26B1+ Olfactory Sensory Neurons Is Necessary for the Functional Integrity of the Olfactory Sensory Map. Journal of Neuroscience 35, 13807–13818. 10.1523/jneurosci.2247-15.2015.

89. Paschaki, M., Cammas, L., Muta, Y., Matsuoka, Y., Mak, S.-S., Rataj-Baniowska, M., Fraulob, V., Dolle, P., and Ladher, R.K. (2013). Retinoic acid regulates olfactory progenitor cell fate and differentiation. Neural Development 8, 13. 10.1186/1749-8104-8-13.

90. LaMantia, A.-S., Bhasin, N., Rhodes, K., and Heemskerk, J. (2000). Mesenchymal/Epithelial Induction Mediates Olfactory Pathway Formation. Neuron 28, 411–425. 10.1016/s0896-6273(00)00121-5.

91. Yang, L.M., Huh, S.-H., and Ornitz, D.M. (2018). FGF20-Expressing, Wnt-Responsive Olfactory Epithelial Progenitors Regulate Underlying Turbinate Growth to Optimize Surface Area. Dev. Cell 46, 564–580.e565. 10.1016/j.devcel.2018.07.010.

92. Enomoto, T., Nishida, H., Iwata, T., Fujita, A., Nakayama, K., Kashiwagi, T., Hatanaka, Y., Kondo, H., Kajitani, R., Itoh, T., et al. (2019). Bcl11b controls odorant receptor class choice in mice. Communications Biology 2, 296. 10.1038/s42003-019-0536-x.

93. Iwata, T., Tomeoka, S., and Hirota, J. (2021). A class I odorant receptor enhancer shares a functional motif with class II enhancers. Scientific Reports 11, 510. 10.1038/s41598-020-79980-x.

94. Shah, A., Ratkowski, M., Rosa, A., Feinstein, P., and Bozza, T. (2021). Olfactory expression of trace amine-associated receptors requires cooperative cis-acting enhancers. Nature Communications 12, 3797. 10.1038/s41467-021-23824-3.

95. Fei, A., Wu, W., Tan, L., Tang, C., Xu, Z., Huo, X., Bao, H., Kong, Y., Johnson, M., Hartmann, G., et al. (2021). Coordination of two enhancers drives expression of olfactory trace amine-associated receptors. Nature Communications 12, 3798. 10.1038/s41467-021-23823-4.

96. Strotmann, J., Wanner, I., Helfrich, T., Beck, A., Meinken, C., Kubick, S., and Breer, H. (1994). Olfactory neurones expressing distinct odorant receptor subtypes are spatially segregated in the nasal neuroepithelium. Cell and Tissue Research 276, 429–438. 10.1007/BF00343941.

97. Liff, C.W., Ayman, Y.R., Jaeger, E.C.B., Lee, H.S., Kim, A., Albarracín, A.V., and Marlin, B.J. (2023). Fear conditioning biases olfactory stem cell receptor fate. eLife Sciences Publications, Ltd.

98. Vihani, A., Hu, X.S., Gundala, S., Koyama, S., Block, E., and Matsunami, H. (2020). Semiochemical responsive olfactory sensory neurons are sexually dimorphic and plastic. eLife 9, e54501. 10.7554/eLife.54501.

99. van der Linden, C.J., Gupta, P., Bhuiya, A.I., Riddick, K.R., Hossain, K., and Santoro, S.W. (2020). Olfactory Stimulation Regulates the Birth of Neurons That Express Specific Odorant Receptors. Cell Reports 33. 10.1016/j.celrep.2020.108210.

100. Hossain, K., Smith, M., Rufenacht, K.E., O’Rourke, R., and Santoro, S.W. (2025). In mice, discrete odors can selectively promote the neurogenesis of sensory neuron subtypes that they stimulate. bioRxiv, 2024.2002.2010.579748. 10.1101/2024.02.10.579748.

101. Monahan, K., Schieren, I., Cheung, J., Mumbey-Wafula, A., Monuki, E.S., and Lomvardas, S. (2017). Cooperative interactions enable singular olfactory receptor expression in mouse olfactory neurons. eLife 6, e28620. 10.7554/eLife.28620.

## Supplemental References

1. Fleischmann, A., Abdus-Saboor, I., Sayed, A., and Shykind, B. (2013). Functional Interrogation of an Odorant Receptor Locus Reveals Multiple Axes of Transcriptional Regulation. PLOS Biology 11, e1001568. 10.1371/journal.pbio.1001568.

2. Yu, C.R., Power, J., Barnea, G., O’Donnell, S., Brown, H.E.V., Osborne, J., Axel, R., and Gogos, J.A. (2004). Spontaneous Neural Activity Is Required for the Establishment and Maintenance of the Olfactory Sensory Map. Neuron 42, 553–566. 10.1016/S0896-6273(04)00224-7.

3. Brunet, L.J., Gold, G.H., and Ngai, J. (1996). General Anosmia Caused by a Targeted Disruption of the Mouse Olfactory Cyclic Nucleotide&#x2013;Gated Cation Channel. Neuron 17, 681–693. 10.1016/S0896-6273(00)80200-7.

4. Hébert, J.M., and McConnell, S.K. (2000). Targeting of cre to the Foxg1 (BF-1) Locus Mediates loxP Recombination in the Telencephalon and Other Developing Head Structures. Developmental Biology 222, 296–306. 10.1006/dbio.2000.9732.

5. Tsukahara, T., Brann, D.H., Pashkovski, S.L., Guitchounts, G., Bozza, T., and Datta, S.R. (2021). A transcriptional rheostat couples past activity to future sensory responses. Cell 184, 6326–6343.e6332. 10.1016/j.cell.2021.11.022.

6. Suzukawa, K., Kondo, K., Kanaya, K., Sakamoto, T., Watanabe, K., Ushio, M., Kaga, K., and Yamasoba, T. (2011). Age-related changes of the regeneration mode in the mouse peripheral olfactory system following olfactotoxic drug methimazole-induced damage. Journal of Comparative Neurology 519, 2154–2174. 10.1002/cne.22611.

7. Hogarth, C.A., Evanoff, R., Mitchell, D., Kent, T., Small, C., Amory, J.K., and Griswold, M.D. (2013). Turning a Spermatogenic Wave into a Tsunami: Synchronizing Murine Spermatogenesis Using WIN 18,4461. Biology of Reproduction 88, 40, 41–49. 10.1095/biolreprod.112.105346.

8. Finlay, J.B., Brann, D.H., Abi Hachem, R., Jang, D.W., Oliva, A.D., Ko, T., Gupta, R., Wellford, S.A., Moseman, E.A., Jang, S.S., et al. (2022). Persistent post–COVID-19 smell loss is associated with immune cell infiltration and altered gene expression in olfactory epithelium. Science Translational Medicine 14, eadd0484. doi:10.1126/scitranslmed.add0484.

9. Choi, H.M.T., Schwarzkopf, M., Fornace, M.E., Acharya, A., Artavanis, G., Stegmaier, J., Cunha, A., and Pierce, N.A. (2018). Third-generation in situ hybridization chain reaction: multiplexed, quantitative, sensitive, versatile, robust. Development 145. 10.1242/dev.165753.

10. Lotfollahi, M., Naghipourfar, M., Luecken, M.D., Khajavi, M., Büttner, M., Wagenstetter, M., Avsec, Ž., Gayoso, A., Yosef, N., Interlandi, M., et al. (2022). Mapping single-cell data to reference atlases by transfer learning. Nature Biotechnology 40, 121–130. 10.1038/s41587-021-01001-7.

11. Sikkema, L., Ramírez-Suástegui, C., Strobl, D.C., Gillett, T.E., Zappia, L., Madissoon, E., Markov, N.S., Zaragosi, L.-E., Ji, Y., Ansari, M., et al. (2023). An integrated cell atlas of the lung in health and disease. Nature Medicine 29, 1563–1577. 10.1038/s41591-023-02327-2.

12. Tan, L., and Xie, X.S. (2018). A Near-Complete Spatial Map of Olfactory Receptors in the Mouse Main Olfactory Epithelium. Chemical Senses 43, 427–432. 10.1093/chemse/bjy030.

13. Miyamichi, K., Serizawa, S., Kimura, H.M., and Sakano, H. (2005). Continuous and Overlapping Expression Domains of Odorant Receptor Genes in the Olfactory Epithelium Determine the Dorsal/Ventral Positioning of Glomeruli in the Olfactory Bulb. The Journal of Neuroscience 25, 3586–3592. 10.1523/jneurosci.0324-05.2005.

14. Kotliar, D., Veres, A., Nagy, M.A., Tabrizi, S., Hodis, E., Melton, D.A., and Sabeti, P.C. (2019). Identifying gene expression programs of cell-type identity and cellular activity with single-cell RNA-Seq. eLife 8, e43803. 10.7554/eLife.43803.

15. Clifton, K., Anant, M., Aihara, G., Atta, L., Aimiuwu, O.K., Kebschull, J.M., Miller, M.I., Tward, D., and Fan, J. (2023). STalign: Alignment of spatial transcriptomics data using diffeomorphic metric mapping. Nature Communications 14, 8123. 10.1038/s41467-023-43915-7.

16. Petukhov, V., Xu, R.J., Soldatov, R.A., Cadinu, P., Khodosevich, K., Moffitt, J.R., and Kharchenko, P.V. (2022). Cell segmentation in imaging-based spatial transcriptomics. Nature Biotechnology 40, 345–354. 10.1038/s41587-021-01044-w.

17. McInnes, L., Healy, J., and Melville, J. (2018). Umap: Uniform manifold approximation and projection for dimension reduction. arXiv preprint arXiv:1802.03426.

18. Jones, D.C., Elz, A.E., Hadadianpour, A., Ryu, H., Glass, D.R., and Newell, E.W. (2024). Cell Simulation as Cell Segmentation. bioRxiv, 2024.2004.2025.591218. 10.1101/2024.04.25.591218.

19. Ruiz Tejada Segura, M.L., Abou Moussa, E., Garabello, E., Nakahara, T.S., Makhlouf, M., Mathew, L.S., Wang, L., Valle, F., Huang, S.S.Y., Mainland, J.D., et al. (2022). A 3D transcriptomics atlas of the mouse nose sheds light on the anatomical logic of smell. Cell Reports 38. 10.1016/j.celrep.2022.110547.

20. Bashkirova, E.V., Klimpert, N., Monahan, K., Campbell, C.E., Osinski, J., Tan, L., Schieren, I., Pourmorady, A., Stecky, B., Barnea, G., et al. (2023). Opposing, spatially-determined epigenetic forces impose restrictions on stochastic olfactory receptor choice. eLife 12, RP87445. 10.7554/eLife.87445.

122. Ewels, P.A., Peltzer, A., Fillinger, S., Patel, H., Alneberg, J., Wilm, A., Garcia, M.U., Di Tommaso, P., and Nahnsen, S. (2020). The nf-core framework for community-curated bioinformatics pipelines. Nature Biotechnology 38, 276–278. 10.1038/s41587-020-0439-x.

22. Dobin, A., Davis, C.A., Schlesinger, F., Drenkow, J., Zaleski, C., Jha, S., Batut, P., Chaisson, M., and Gingeras, T.R. (2013). STAR: ultrafast universal RNA-seq aligner. Bioinformatics 29, 15–21. 10.1093/bioinformatics/bts635.

23. Li, B., and Dewey, C.N. (2011). RSEM: accurate transcript quantification from RNA-Seq data with or without a reference genome. BMC Bioinformatics 12, 323. 10.1186/1471-2105-12-323.

24. Bintu, B. (2021). Genome-Scale Imaging: From the Subcellular Structure of Chromatin to the 3D Organization of the Peripheral Olfactory System. Ph.D. (Harvard University).

25. Kenney, M., Vasylieva, I., Hood, G., Cao-Berg, I., Tuite, L., Laghaei, R., Smith, M.C., Watson, A.M., and Ropelewski, A.J. (2024). The Brain Image Library: A Community-Contributed Microscopy Resource for Neuroscientists. Scientific Data 11, 1212. 10.1038/s41597-024-03761-8.

26. Tirosh, I., Izar, B., Prakadan, S.M., Wadsworth, M.H., Treacy, D., Trombetta, J.J., Rotem, A., Rodman, C., Lian, C., Murphy, G., et al. (2016). Dissecting the multicellular ecosystem of metastatic melanoma by single-cell RNA-seq. Science 352, 189–196. 10.1126/science.aad0501.

27. Weinreb, C., Rodriguez-Fraticelli, A., Camargo, F.D., and Klein, A.M. (2020). Lineage tracing on transcriptional landscapes links state to fate during differentiation. Science 367, eaaw3381. 10.1126/science.aaw3381.

28. Keane, T.M., Goodstadt, L., Danecek, P., White, M.A., Wong, K., Yalcin, B., Heger, A., Agam, A., Slater, G., Goodson, M., et al. (2011). Mouse genomic variation and its effect on phenotypes and gene regulation. Nature 477, 289–294. 10.1038/nature10413.

29. Lilue, J., Doran, A.G., Fiddes, I.T., Abrudan, M., Armstrong, J., Bennett, R., Chow, W., Collins, J., Collins, S., Czechanski, A., et al. (2018). Sixteen diverse laboratory mouse reference genomes define strain-specific haplotypes and novel functional loci. Nature Genetics 50, 1574–1583. 10.1038/s41588-018-0223-8.

30. Rogozhnikov, A., Ramkumar, P., Shah, K., Bedi, R., Kato, S., and Escola, G.S. (2021). Demuxalot: scaled up genetic demultiplexing for single-cell sequencing. bioRxiv, 2021.2005.2022.443646. 10.1101/2021.05.22.443646.

31. Cingolani, P., Adrian, P., Lily, W.L., Melissa, C., Tung, N., Luan, W., J., L.S., Xiangyi, L., and and Ruden, D.M. (2012). A program for annotating and predicting the effects of single nucleotide polymorphisms, SnpEff. Fly 6, 80–92. 10.4161/fly.19695.

32. Monahan, K., Schieren, I., Cheung, J., Mumbey-Wafula, A., Monuki, E.S., and Lomvardas, S. (2017). Cooperative interactions enable singular olfactory receptor expression in mouse olfactory neurons. eLife 6, e28620. 10.7554/eLife.28620.

33. Langmead, B., and Salzberg, S.L. (2012). Fast gapped-read alignment with Bowtie 2. Nature Methods 9, 357–359. 10.1038/nmeth.1923.

34. Heinz, S., Benner, C., Spann, N., Bertolino, E., Lin, Y.C., Laslo, P., Cheng, J.X., Murre, C., Singh, H., and Glass, C.K. (2010). Simple Combinations of Lineage-Determining Transcription Factors Prime *cis*-Regulatory Elements Required for Macrophage and B Cell Identities. Molecular Cell 38, 576–589. 10.1016/j.molcel.2010.05.004.

136. Klimpert, N., Kollo, M., Brann, D.H., Tan, C., Barry, D., Ma, Y., Schaefer, A.T., and Fleischmann, A. (2025). 3D spatial transcriptomics reveals the molecular domain structure of the mouse olfactory bulb. bioRxiv, 2025.2002.2019.639192. 10.1101/2025.02.19.639192.

36. Zapiec, B., and Mombaerts, P. (2015). Multiplex assessment of the positions of odorant receptor-specific glomeruli in the mouse olfactory bulb by serial two-photon tomography. Proceedings of the National Academy of Sciences 112, E5873–E5882. 10.1073/pnas.1512135112.

37. Wolf, F.A., Angerer, P., and Theis, F.J. (2018). SCANPY: large-scale single-cell gene expression data analysis. Genome Biology 19, 15. 10.1186/s13059-017-1382-0.

38. Gayoso, A., Lopez, R., Xing, G., Boyeau, P., Valiollah Pour Amiri, V., Hong, J., Wu, K., Jayasuriya, M., Mehlman, E., Langevin, M., et al. (2022). A Python library for probabilistic analysis of single-cell omics data. Nature Biotechnology 40, 163–166. 10.1038/s41587-021-01206-w.

